# AADAT-Driven Metabolic Control of Malate and CoQ_10_ Shapes Immune Evasion in Triple-Negative Breast Cancer

**DOI:** 10.64898/2026.01.28.702389

**Authors:** Megha Chatterjee, Franklin Gu, Susmita Samanta, Uttam Rasaily, Sai Manohar Thota, Dana Varghese, Yunping Qiu, Lynette Ewura Esi Fordwuo, Hugo Villanueva, Mary Kathryn McKenna, Jun Hyoung Park, Weijie Zhang, Lin Tian, Liqun Yu, Badrajee Piyarathna, Yang Gao, Brian Wesley Simons, Sung Yun Jung, Balasubramanyam Karanam, Vasanta Putluri, Nagi Chandandeep, Nada Mohamed, Jaya Ruth Asirvatham, Deborah Jebakumar, Arundati Rao, Carolina Gutierrez, Angela R Omilian, Carl Morrison, Gokul M Das, Christine Ambrosone, Erin H Seeley, Benny Abraham Kaipparettu, Irwin J. Kurland, Nagireddy Putluri, Ahmed Elkhanany, Andrew A. Davis, Qian Zhu, Xiang H.-F. Zhang, Arun Sreekumar

## Abstract

Compared to other subtypes of breast cancer, triple-negative breast cancers (TNBC) have fewer treatment options and exhibit a worse prognosis. Through integrated transcriptomic, metabolomic, immunohistochemical, spatial, and clinical analyses, we identify the mitochondrial enzyme, α-aminoadipate aminotransferase (AADAT) as a previously unrecognized metabolic immune checkpoint in TNBC. *AADAT* mRNA and protein were significantly upregulated in human TNBC, and high AADAT expression was associated with reduced intra-tumoral CD8⁺ T-cell density and inferior survival. Genetic silencing of *AADAT* in orthotopic murine TNBC models curtailed primary tumor growth and distant metastasis in a CD8⁺ T-cell–dependent manner, enhanced effector T-cell activation, and sensitized tumors to dual PD-1/CTLA-4 blockade. Mechanistically, unbiased metabolomics showed increased malate levels after *AADAT* knockdown. Additionally, 4-hydroxyphenylpyruvate, an essential precursor for coenzyme Q_10_(CoQ_10_) biosynthesis, decreased following *AADAT* knockdown, suggesting an impaired mitochondrial electron transport chain. CoQ_10_ supplementation restored metabolic balance and reversed malate accumulation caused by *AADAT* knockdown, indicating that AADAT helps maintain CoQ_10_-supported redox homeostasis, thereby preventing malate buildup and export. Notably, malate addition directly boosted CD8⁺ T-cell oxidative metabolism, increased the NAD⁺/NADH ratio and reactive oxygen species, and augmented TNF-α and IFN-γ production. In vivo, malate supplementation in drinking water phenocopied AADAT knockdown, restored the response to paclitaxel plus anti–PD-1 therapy in multiple independent syngeneic TNBC models with de novo or acquired resistance to immunotherapy, reduced tumor burden, and prolonged survival. In patient cohorts, higher spatially clustered intra-tumoral malate is associated with co-localization of functional CD8⁺ T cells, decreased exhausted T-cell neighborhoods, and superior post-chemotherapy outcomes. These data position AADAT as a central metabolic orchestrator of immune escape in TNBC and nominate oral malate as a readily translatable adjuvant to reverse chemo-immunotherapy resistance in TNBC.

**Statement of Significance:** AADAT defines a metabolic-immune axis driving immune evasion and therapy resistance in triple-negative breast cancer. Blocking AADAT or administering oral malate reactivates CD8⁺ T-cell immunity and sensitizes chemo-immunotherapy-resistant tumors to these agents. These findings uncover a readily translatable metabolic vulnerability with potential to improve outcomes for patients with aggressive breast cancer subtypes.

## Introduction

Triple-negative breast cancer (TNBC) is a heterogeneous disease with poor prognosis due to a high propensity for early metastasis and recurrence following standard-of-care chemotherapy^1^. The presence of tumor-infiltrating lymphocytes (TILs), particularly cytotoxic CD8⁺ T cells, has been strongly associated with improved response to neoadjuvant chemotherapy and favorable survival outcomes in TNBC patients^2–4^. These findings have provided a rationale for the use of immune checkpoint blockade (ICB), including therapies targeting CTLA-4 and the PD-1/PD-L1 axis, to enhance anti-tumor immune responses in TNBC^5^. While the addition of ICB to chemotherapy has shown clinical benefit in a subset of patients—most notably in the KEYNOTE-522 trial, where pembrolizumab improved pathologic complete response (pCR) from 50% to 65% and increased overall survival from 81.7% to 86.6%—many patients remain unresponsive or develop resistance to treatment^6–9^. Moreover, the high incidence of immune-related adverse events^10,11^ underscores the need for more targeted, tolerable strategies to sensitize tumors to ICB.

A significant obstacle to optimizing immunotherapy is the immunosuppressive tumor microenvironment (TME), shaped in part by tumor-intrinsic metabolic programs ^12^. In TNBC, the metabolic crosstalk between tumor and immune cells profoundly influences antigen presentation, T cell infiltration, and effector function. For example, hypoxia-driven metabolic changes in the TME can reduce MHC expression ^13^, suppress interferon-gamma (IFN-γ) production ^14^, upregulate PD-L1 ^13^, and enhance aerobic glycolysis ^13^—all of which facilitate immune escape. Beyond hypoxia, many metabolic intermediates directly modulate immune cell differentiation and function^15–18^, and reprogramming these axes has emerged as a promising avenue to overcome therapeutic resistance.

Tryptophan catabolism through the kynurenine pathway (KP) illustrates a mechanism of immune evasion where the depletion of tryptophan and the buildup of immunosuppressive metabolites suppress cytotoxic T cells and promote regulatory T cells. The KP’s first enzymatic step, catalyzed by indoleamine 2,3-dioxygenase (IDO1) and tryptophan 2,3-dioxygenase (TDO2), has been associated with poor prognosis and immune suppression in breast tumors^19–21^. While most studies have focused on IDO1 and TDO2, recent evidence suggests that downstream KP enzymes, including kynureninase and kynurenine-3-monooxygenase, may also contribute to TNBC progression^22,23^. However, the immunoregulatory potential of other KP enzymes remains poorly defined.

In this context, we identified the mitochondrial enzyme alpha-aminoadipate aminotransferase (AADAT), also known as kynurenine aminotransferase, as a previously uncharacterized immunosuppressive regulator in TNBC. AADAT catalyzes the conversion of kynurenine to kynurenic acid ^24^ and, in our study, was found to be markedly upregulated in TNBC tumors. Beyond its canonical aminotransferase activity, AADAT also functions as a mitochondrial tyrosine transaminase^25^ that generates 4-hydroxyphenylpyruvate, an obligate precursor of 4-hydroxybenzoate and thus the benzoquinone headgroup of coenzyme Q_10_ (CoQ_10_), coupling KP flux to mitochondrial CoQ_10_ biosynthesis^26^. In mammalian cancer cells, stress-induced up-regulation of CoQ_10_ biosynthetic genes enhances oxidative phosphorylation and antioxidant capacity. At the same time, pharmacologic blockade of CoQ_10_ synthesis increases chemotherapy-induced cytotoxicity^27^.

Our research reveals a new link between AADAT and anti-tumor immunity in TNBC. Specifically, AADAT suppresses CD8+ T cell activity within the tumor microenvironment by lowering secreted malate levels and preserving CoQ_10_-dependent redox balance in the tumors. Increasing malate in the tumor microenvironment, either by blocking AADAT or by supplementing the metabolite, reprograms CD8+ T cell metabolism and function. This restores response to immune checkpoint inhibitors and chemotherapy in de novo or acquired immunotherapy-resistant TNBC and increases patient survival without added toxicity. The findings highlight malate as an essential immunoregulatory metabolite and suggest that metabolic reprogramming could be a valuable strategy to overcome immune resistance in aggressive breast cancers.

## Results

### AADAT Expression Correlates with Reduced CD8⁺ TILs and Poor Prognosis in TNBC

To investigate the role of the kynurenine pathway (KP, **Fig. 1a**) in immune escape, we examined publicly available breast cancer patient-derived microarray and IHC data on KP gene expression and CD8⁺ TILs in breast tumors^27^. As expected from an earlier study (28), higher levels of KP enzymes such as IDO1, IDO2, and HAAO were linked to increased CD8⁺ TILs (**Figs. S1a–c**). Notably, only Alpha-Amino Adipate Amino Transferase (AADAT) expression was uniquely associated with reduced CD8⁺ TIL infiltration (**Fig. 1b, c).**

**Figure 1.**
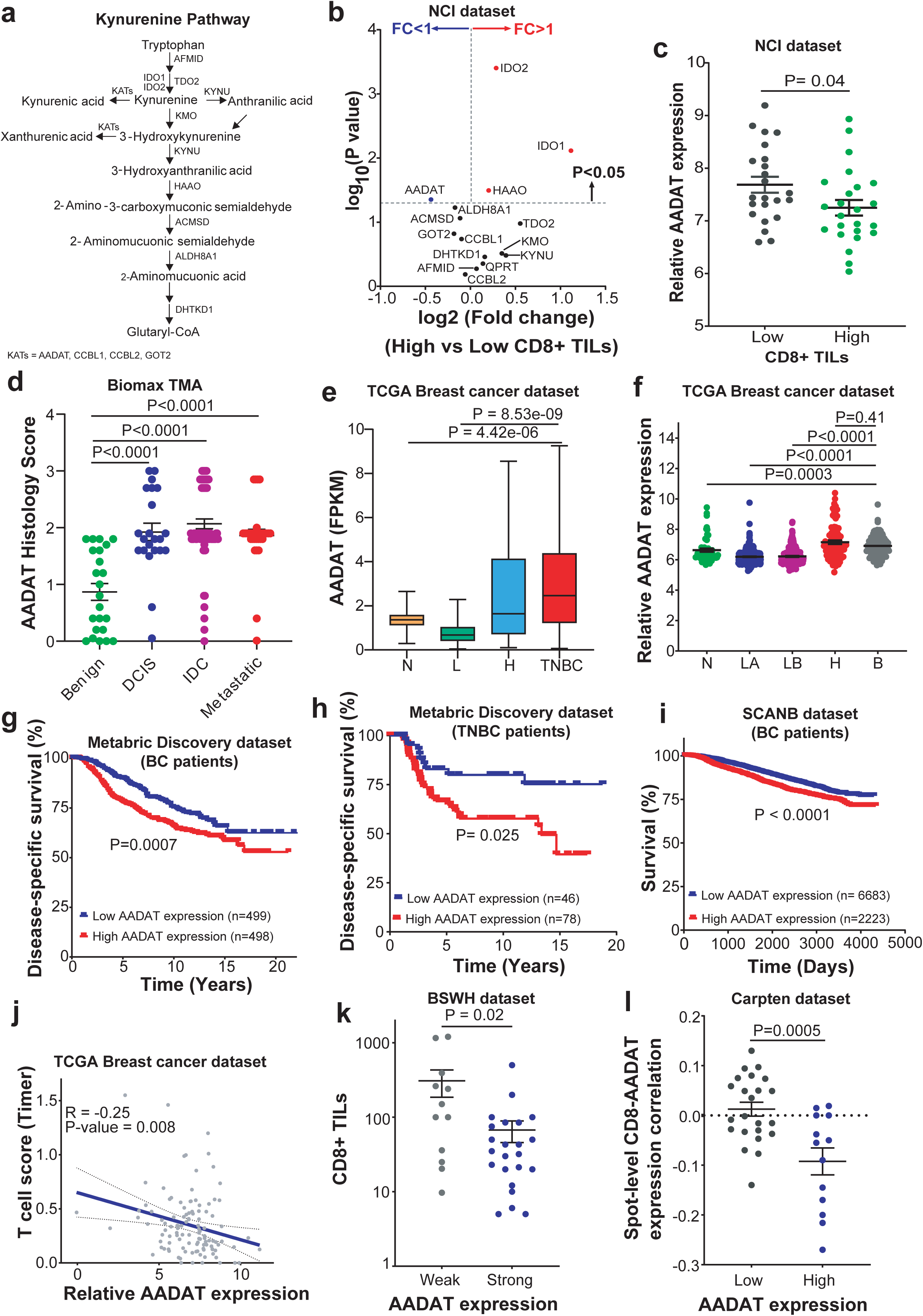
AADAT expression in aggressive breast tumors is associated with decreased CD8+ T cell infiltration and worse clinical outcomes. (a) Schematic of the KP based on annotated pathways in Metacyc (PWY-6309) and KEGG (hsa00380) databases. (b) Volcano plot representing changes in the expression of KP genes in breast tumors in the NCI dataset (GSE37751) with high (n = 24) vs. low (n = 23) intra-tumoral CD8+ TILs based on median CD8+ TIL count. Red circles, upregulated genes; blue circles, downregulated genes. (c) Dot plot showing *AADAT* mRNA expression in breast tumors from the NCI dataset (GSE37751), divided into two subgroups of low (black, n = 23) and high (green, n = 24) CD8+ TILs divided based on the median number of intra-tumoral CD8+ TILs. Data are presented as mean ± SEM. P-value computed using Student’s t-test. (d) Dot plot showing immunohistochemical staining of AADAT scored by a pathologist in benign breast tissues (n = 23), ductal carcinoma in situ (DCIS, n = 22), invasive ductal carcinoma (IDC, n = 67), and metastatic IDC samples (n = 32) in a human breast cancer TMA (BR2082a, US Biomax). Data are presented as mean ± SEM. P-value was computed using a two-tailed Mann-Whitney test. (e) Box plot showing *AADAT* mRNA expression in breast tumors from the TCGA dataset across four PAM50 subtypes: Normal-like (n = 113), Luminal (n = 575), HER2-enriched (n =37), and TNBC (n = 115). Data are presented as mean ± SEM. P-value computed using Student’s t-test. (f) Dot plot showing *AADAT* mRNA expression in breast tumors from the METABRIC Discovery dataset across five PAM50 subtypes: Normal-like (n = 58), Luminal A (n = 466), Luminal B (n = 268), HER2-enriched (n = 87), and Basal-like (n = 118). Data are presented as mean ± SEM. P-values were computed using a two-tailed Mann-Whitney test. (g) Kaplan-Meier survival curves of disease-specific survival in breast cancer patients from the METABRIC Discovery dataset, stratified based on the median expression value of *AADAT*. The log-rank (Mantel-Cox) test was used to determine the P-value. (h) Kaplan-Meier survival curves of disease-specific survival in triple-negative breast cancer (TNBC) patients from the METABRIC Discovery dataset stratified by the median expression value of *AADAT*. The log-rank (Mantel-Cox) test was used to determine the P-value. (i) Kaplan-Meier survival curves of disease-specific survival in breast cancer patients from the SCANB dataset obtained from the ULCAN portal (https://ualcan.path.uab.edu). The median expression value of *AADAT* stratified the samples in the dataset. The log-rank (Mantel-Cox) test was used to determine the P-value. (j) Correlation plot between TIMER T cell score and *AADAT* mRNA expression in the TCGA breast cancer dataset (n = 112). Pearson correlation coefficient and two-tailed P-value are indicated. Dot plots showing infiltration of CD8+ T cells in TNBC tumors in the Baylor Scott and White dataset with weak (n = 12) or strong (n = 23) AADAT immunohistochemical staining. Data are presented as mean ± SEM. P-values computed using the two-tailed Mann-Whitney U test. (l) Dot plots showing the correlation between CD8+ T cells and low (n = 23) and high (n = 13), expression of *AADAT* in the spatial TNBC transcriptomics dataset published by Bassiouni etal (GSE210616). Data are presented as mean ± SEM. P-values computed using the student’s t-test.

Tissue Microarray (TMA) analysis revealed that AADAT was elevated in malignant breast tumors relative to adjacent benign tissue, without differences between localized and metastatic disease (**Fig. 1d**). TCGA analysis confirmed that *AADAT* expression was increased in the more aggressive HER2-like and TNBC subtypes compared to Luminal and Normal tumors (**Fig. 1e**), findings corroborated in the METABRIC dataset^28^, where *AADAT* was enriched in HER2-enriched and Basal-like subtypes (**Fig. 1f**). Analysis of multiple publicly available gene expression datasets in The University of Alabama at Birmingham Cancer data analysis portal (UALCAN) ^29^ further demonstrated that *AADAT* expression is significantly higher in ER-negative or TNBC tumors compared to ER-positive tumors or adjacent benign tissues (**Figs. S1 d-j**). AADAT protein expression was also elevated in TNBC tumors compared to luminal and HER2+ breast cancers in the CPTAC dataset (**Fig. S1k**). Across datasets, higher *AADAT* expression predicted worse clinical outcomes in breast cancer and within TNBC (**Figs. 1g–i**). In TNBC samples, *AADAT* expression negatively correlated with predicted T cell infiltration using TIMER (Tumor Immune Estimation Resource ^30^) analysis (**Fig. 1j**), a finding confirmed by IHC on TNBC TMAs showing fewer CD3+ and CD8⁺ TILs in AADAT-high tumors (**Figs.1k, S2a**) and validated using a spatial gene expression dataset ^31^ (**Fig. 1l**). Gene set enrichment analysis on NanoString data ^32^ revealed enrichment of cytotoxic T cell–associated signatures, including allograft rejection and IFN-γ response, in TNBC samples with low *AADAT* expression (**Figs. S2 b-d**), supporting an inverse relationship between AADAT and CD8⁺ TILs.

### AADAT Promotes Tumor Growth and Suppresses Immune Infiltration in TNBC

To determine whether AADAT directly influences immune exclusion, we knocked down (KD) AADAT in E0771 and 4T1 cells using two independent shRNAs (shAADAT-1 and shAADAT-2) delivered by a lentiviral vector, along with reconstituting AADAT expression for rescue (shAADAT-2+AADAT). We confirmed KD and rescue through qPCR, peptide reaction monitoring (PRM) proteomics, and a functional assessment of the KYNA: KYN ratio (**Figs. 2a, S3a, b**). Remarkably, *AADAT* KD significantly reduced tumor initiation and delayed tumor growth following orthotopic injection of both E0771 and 4T1 cells into the mammary fat pad of syngeneic wild-type C57BL/6J and BALB/cJ female mice, respectively, which was rescued by re-expression of *AADAT* (**Figs. 2b, c, and S3c, d**). In CD8 KO mice, *AADAT* KD did not delay tumor initiation, indicating that CD8⁺ T cells are required for the observed delay (**Figs. S3e, f**). However, once established, *AADAT* KD tumors grew more slowly in CD8 KO mice (**Figs. 2d, e**), suggesting the presence of additional CD8+ T cell-independent mechanisms. Flow cytometry analysis of E0771 tumors revealed increased infiltration of immune cells, including monocytes, macrophages, B cells, and CD4⁺ T cells after *AADAT* KD (**Figs. S3 g-j**), indicating that AADAT impacts the immune landscape in TNBC. To study metastasis, we created an inducible AADAT KD in E0771 cell line (**Fig. 2f**). Spontaneous pulmonary metastasis, assessed one month following resection of size-matched primary tumors derived from the above-mentioned cells, was significantly decreased in mice with induced AADAT KD tumors compared to mice with control tumors that were not given doxycycline (**Fig. 2g**).

**Figure 2.**
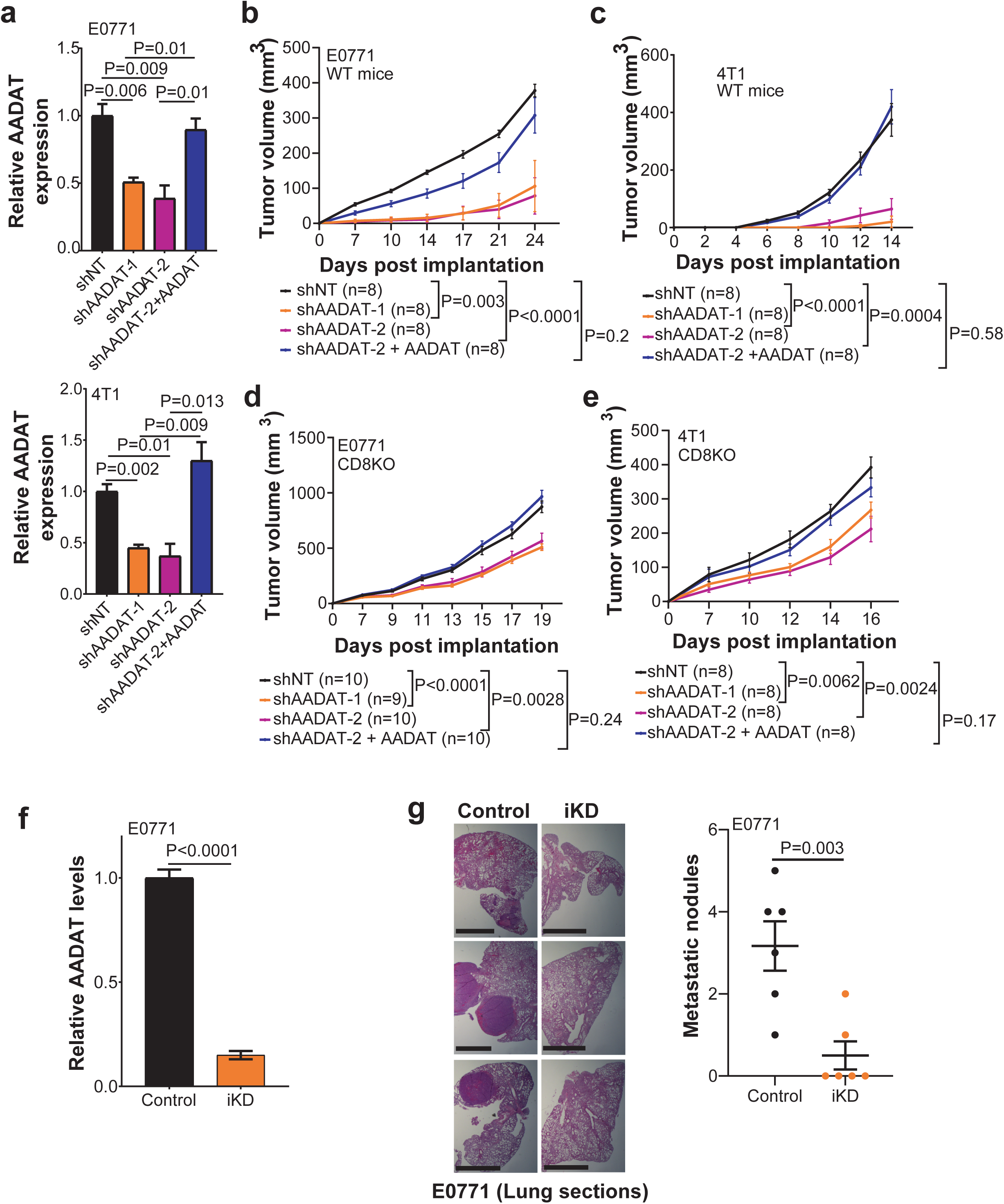
AADAT expression promotes tumor formation in two independent syngeneic immunocompetent mouse models. (a) RT-qPCR analysis for *AADAT* in E0771 and 4T1 cells stably transduced with control non-targeted shRNA (shNT), Aadat-specific shRNAs (shAadat), and an Aadat re-expression construct, each n=3 biological replicates. P-value computed using Student’s t-test. (b) Tumor growth analysis of wild-type C57BL/6J mice orthotopically implanted with the syngeneic E0771 cell lines described in panel a. P-values were calculated using multiple unpaired Student’s t-test. The indicated P-values are for the final study time point, i.e., Day 24. The n values are also indicated. (c) Tumor growth analysis of wild-type BALB/cJ mice orthotopically implanted with the syngeneic 4T1 cell lines stably transduced with either control non-targeted shRNA (shNT), Aadat-specific shRNAs (shAadat), and an Aadat re-expression, as described in panel a. P-values were calculated using multiple unpaired Student’s t-test. The indicated P-values are from the last study time point, i.e., Day 14. n values are shown. (d) Tumor growth analysis of C57BL/6J mice with CD8 knockout (CD8KO), orthotopically implanted with the syngeneic E0771 cell line (panel a). P-values were calculated using multiple unpaired Student’s t-tests. The indicated P values are from the final study time point, i.e., Day 19. (e) Tumor growth analysis of BALB/cJ mice with CD8 knockout (CD8KO), orthotopically implanted with the syngeneic E0771 cell line (panel a). P-values were calculated using multiple unpaired Student’s t-tests. The indicated P values are from the last study time point, i.e., Day 16. (f) RT-qPCR analysis for *AADAT* in E0771 cells containing doxycycline-inducible Aadat-specific shRNAs (iKD1) compared to uninduced control (control). The average of three biological replicates is shown. P-value computed using an unpaired two-tailed Student’s t-test. (g) Left panel: Representative H&E images of lung sections from wild-type C57BL/6J mice four weeks following resection of size-matched E0771 tumors expressing doxycycline-induced Aadat-specific shRNA (iKD1, n = 6, per treatment group). The primary tumors were resected at 300 mm^3^ followed by induction of AADAT KD for 28 days using Doxycycline and quantification of lung metastases. Scale bar: 1 mm. Right Panel: Metastatic nodules were visually quantified in lung sections in Control and iKD1 mice. P-values computed using an unpaired two-tailed Student’s t-test.

### AADAT Inhibition Sensitizes TNBC to Immune Checkpoint Blockade

To determine whether the immunomodulatory role of AADAT extends to clinical outcomes, we evaluated its association with response to immune checkpoint therapy. Analysis of a melanoma dataset (GSE91061) showed that low *AADAT* expression in patients was significantly associated with better immunotherapy response, indicated by a higher pathological response in patients with low versus high AADAT tumors (58% vs 33%, **Fig. 3a**). Based on this, we tested whether AADAT depletion could improve immunotherapy effectiveness in orthotopic TNBC mouse models. Inducible *AADAT* KD, combined with anti–PD–1 and anti–CTLA–4, significantly reduced tumor growth compared to monotherapy in the E0771 and 4T1 models (Figs. **3b–d**). Analysis of *AADAT* KD and control 4T1 tumors, shown in **Fig. 3c**, using imaging mass cytometry (IMC) with a panel of 14 antibodies **(Supplementary Table 1)** targeting murine immune and tumor cells (**Fig. S4**), revealed significantly increased interactions (see Methods for details) between tumor cells and CD8⁺ T cells in *AADAT* KD compared to control tumors (**Fig. 3e**). This suggests that targeting AADAT enhances immune-mediated tumor clearance.

**Figure 3.**
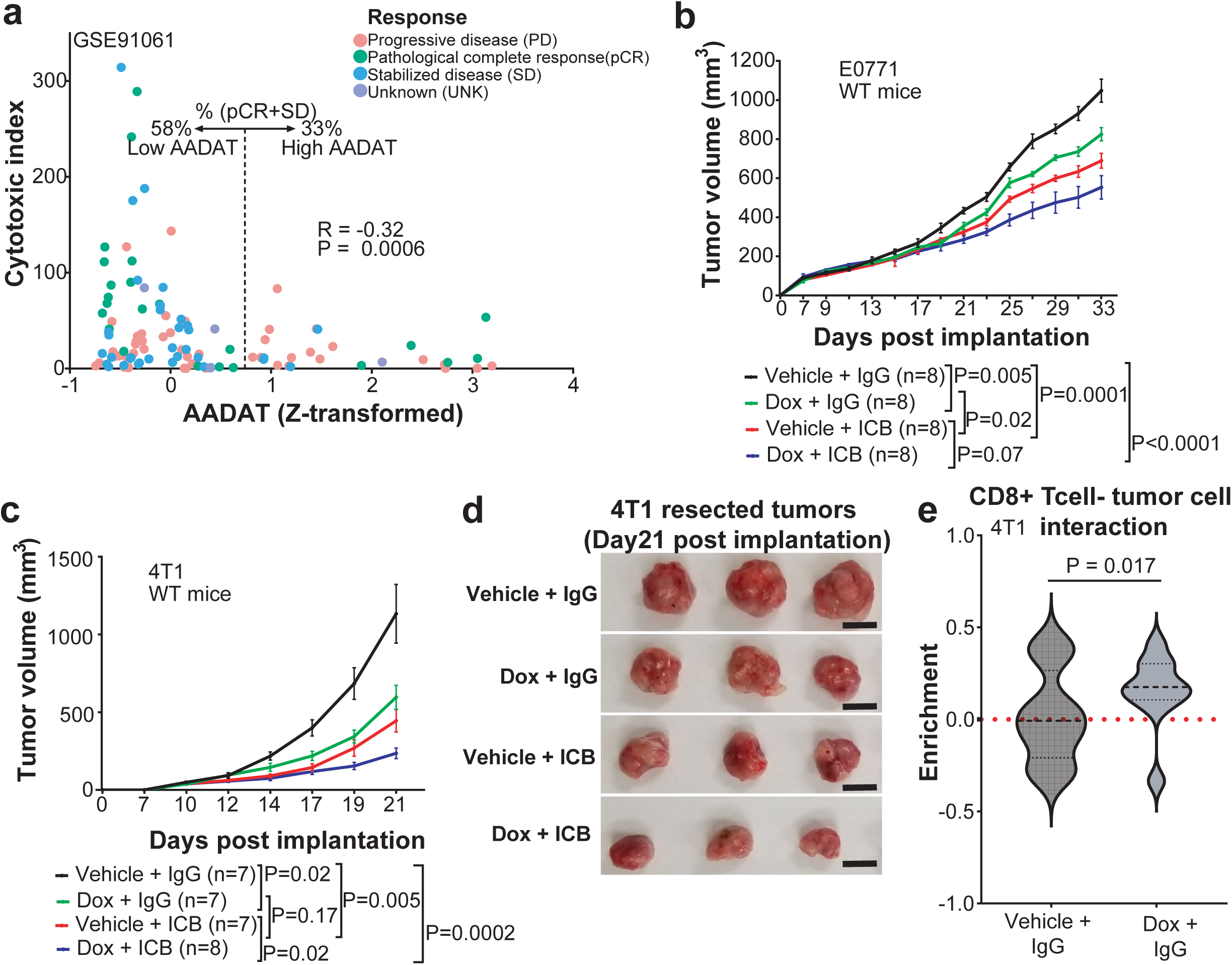
Inducible AADAT knockdown has an additive effect on the response of breast tumors to immune checkpoint blockade therapy. (a) Analysis of clinical and transcriptomic data (GSE91061) from melanoma patients treated with immunotherapy, divided into groups with high (n = 27) or low (n = 82) expression of *AADAT*, revealed increased response rates (pCR + SD) in the low *AADAT* group (58%) vs the high *AADAT* group (33%). Correlation is computed by performing the Spearman correlation test, and the corresponding R and two-tailed P value are indicated. (b) Tumor growth analysis of E0771 cells expressing doxycycline-inducible Aadat-specific shRNA (iKD1) orthotopically implanted into syngeneic wild-type C57BL/6J mice. The mice were treated with one of four regimens: 1) vehicle + IgG (control), 2) doxycycline + IgG (genetic KD of *AADAT*), 3) vehicle + ICB (immune checkpoint blockade therapy), and 4) doxycycline + ICB (*AADAT* KD + immune checkpoint blockade therapy). Multiple unpaired Student’s t-tests determined the P-value. The indicated P values are from the final study time point, i.e., Day 33. n values are shown. (c) Tumor growth analysis of 4T1 cells expressing doxycycline-inducible Aadat-specific shRNA, orthotopically implanted into syngeneic wild-type BALB/cJ mice. The mice were treated with either four regimens: 1) vehicle + IgG (control), 2) doxycycline + IgG (genetic KD of *AADAT*), 3) vehicle + ICB (immune checkpoint blockade therapy), and 4) doxycycline + ICB (*AADAT* KD + immune checkpoint blockade therapy). Multiple unpaired Student’s t-tests determined the P-values. The indicated P-values are from the last study time point, i.e., Day 21. The n values are shown. (d) Representative orthotopic 4T1 tumors resected 21 days after implantation from the experiment shown in panel c. Scale bar: 1 cm. (e) Violin plot illustrating the spatial co-localization and higher interaction of CD8+ T cells and tumor cells in the AADAT iKD and control tumors shown in panels c and d. CD8+ T cells and tumor cells were identified through imaging mass cytometry analysis of paraffin-embedded 4T1 tumors (Control; n=5 mice, 16 regions of interest or ROIs; AADAT*-*iKD1 n=4 mice, 10 ROIs) from the experiment described in panel c. Interactions are represented in Z-scores obtained from Giotto and the significance was calculated using a two-tailed t-test.

### AADAT reduces the buildup and release of immunostimulatory malate

Given that AADAT depletion enhances antitumor immunity, we next explored the metabolic basis for this phenotype by profiling the secretome of AADAT-deficient TNBC cells. Unbiased metabolomics revealed increased malate levels in conditioned media (CM) after *AADAT* KD in E0771-ova+ cells (**Fig. 4a**). To determine the clinical relevance of this finding, we examined the relationship between intra-tumoral malate levels, immune infiltration, and clinical outcomes in TNBC patients.

**Figure 4.**
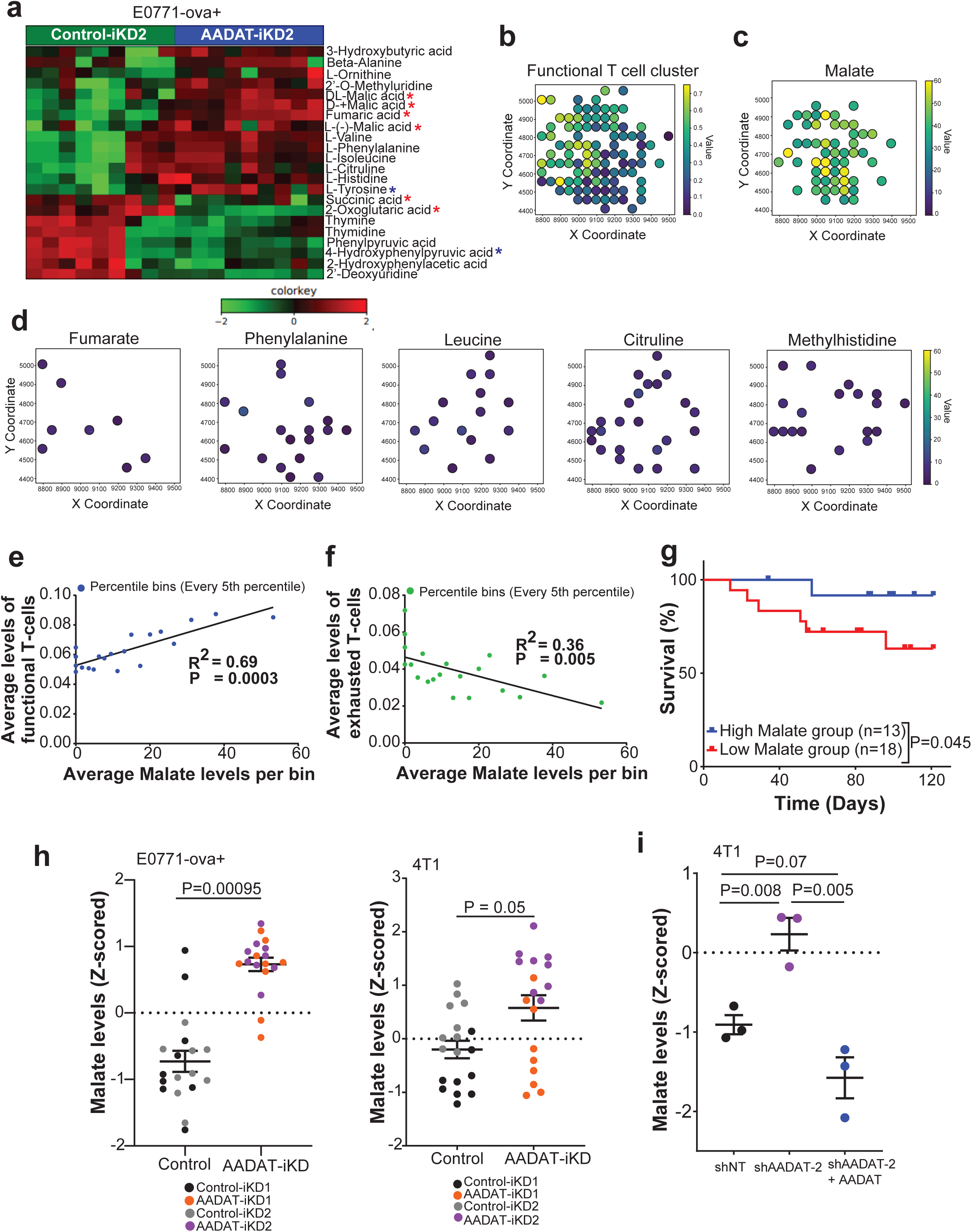
AADAT reduces the buildup and release of immunostimulatory malate. (a) Heatmap of significantly altered metabolites (P<0.05, FDR <0.25) from an unbiased metabolomics analysis of conditioned media (CM) from E0771-ova+ TNBC cells with doxycycline-induced KD of AADAT (AADAT-iKD2) or uninduced control (Control-iKD2, n=3 biological replicates, each with three technical replicates). Shades of red and green indicate metabolites that are significantly elevated or depleted, respectively (see color scale). Metabolites belonging to the tricarboxylic acid cycle are marked with a red asterisk. CoQ_10_ precursors are marked with a blue double asterisk. (b) Representative photomicrographs of a region of interest in a patient’s TNBC tumor show similar spatial clustering patterns for the CD8+ functional T cell cluster and (c) Malate, but not for (d) Fumaric acid, Phenylalanine, Leucine, Citruline, and Methylhistidine, all of which were elevated in the CM of E0771-OVA+ cells containing AADAT-KD (see Fig. 4a). Spatial profiles of metabolites and CD8+ functional T cells were derived from 31 TNBC tumors using two consecutive 5-micron sections of a Tissue Microarray, integrating data from imaging mass spectrometry (about 20-cell resolution) and imaging mass cytometry (single-cell resolution). Each dot represents a pixel indicating the levels (see scale bars) of metabolites or of a CD8+ functional T cell cluster. (e) Plot showing a positive correlation between average levels of functional T-cell clusters and malate. A total of 5245 aligned IMS-IMC spots from 31 TNBC patient samples were examined to explore the relationship between functional T-cells and malate levels at each location. For data visualization, spots were categorized into 10-percentile bins, and the mean values of both functional T cells and malate within each bin were plotted. A Pearson correlation coefficient was calculated, along with the P-value, using a one-tailed Student’s t-test. Results showed a significant positive correlation between functional T cells and malate across the binned spots in the TNBC tumors. (f) Same as in (e) but demonstrating a negative correlation between exhausted T cells and malate. (g) The average pixel intensity of malate, measured by imaging mass spectrometry (IMS), was assessed in each tissue core of the TNBC tissue microarray, and patients were stratified into high (n=13) and low (n=18) malate groups based on the median. An inset shows representative cores with high and low malate levels. The P-value was determined using a two-tailed unpaired Student’s t-test. (h) Kaplan-Meier plot illustrating disease-specific survival among TNBC patients, divided by median spatial malate levels. The P-value was determined using the log-rank (Mantel-Cox) test. The count of patients in each group is shown. (i) Dot plot showing Z-score normalized malate levels in conditioned media of E0771-ova+ and 4T1 cells, with two independent doxycycline-induced AADAT knockdowns (AADAT-iKD1 and AADAT-iKD2) compared to their uninduced controls (Control-iKD1 and Control-iKD2). Each group (iKD1 and iKD2) includes three biological replicates, each with three technical replicates. P values comparing induced KD to uninduced control within each group were calculated using an unpaired two-tailed Student’s t-test. The overall P value was obtained by combining the two group P values with Fisher’s method. (j) Similar to panel i, but in 4T1 cells stably transduced with either control non-targeted shRNA (shNT), Aadat-specific shRNAs (shAadat-2), or an Aadat re-expression construct (shAADAT-2+AADAT), each with three biological replicates. P values were determined using an unpaired two-tailed Student’s t-test.

Using high-resolution imaging mass spectrometry (IMS) and IMC on TNBC TMAs (n=31, **Supplementary Table 2** for clinical data), we quantified malate and immune cell markers **(Supplementary Table 3)** in their spatial context. We then aligned IMS malate spots with IMC spots, yielding 5,245 matched pairs. We analyzed the correlation between IMS malate spots and IMC spots for both functional (CD3, CD4, CD8a, CD45, CD45RO) and exhausted T cell markers (CD8a, Ki67, PD-L1, PD-1, CD152, VEGF) across 31 TNBC samples. **Figures 4b and 4c** show a very similar spatial pattern between functional T cell markers and malate. Notably, this spatial similarity is not seen with fumarate, phenylalanine, leucine, citrulline, and methyl-histidine (**Fig. 4d**), all of which were also elevated in the conditioned media (CM) following AADAT KD in E0771-ova+ cells (**Fig. 4a**). Furthermore, analysis revealed a positive correlation between malate levels and the IMC cluster of functional T cells, and a negative correlation with the cluster of exhausted T cells **(Figs. 4e, f**). However, no significant correlation was observed between the remaining metabolites mentioned above, except methyl histidine, and either the functional T-cell or the exhausted T-cell clusters **(Fig. S5).** Interestingly, TNBC patients with a higher number of intra-tumoral malate-enriched IMS spots had better 10-year survival after standard chemotherapy compared to those with fewer such spots (**Fig. 4g**). Additionally, Cox-Proportional Hazard analysis indicated a hazard ratio of 0.835 (P<0.05) for co-localized malate levels with functional T cells, after adjusting for age and cancer stage (**Supplementary Table 4**). These data suggest that malate serves as a biomarker of a “hot” immune microenvironment and may functionally enhance anti-tumor immunity in TNBC.

We next confirmed elevated levels of secreted malate in murine E0771-ova+ and 4T1 cells by targeted metabolomics, which either had inducible KDs of *AADAT* (**Figs. S6a, b**-AADAT expression levels and **Figs. 4 h-** malate levels) or a constitutive KD of AADAT (**Fig. 2a bottom panel**-AADAT expression levels **and Fig. 4i**-malate levels). Additionally, it was validated in human MDA-MB-231 TNBC cells with separate shRNA-mediated constitutive KDs of *AADAT* (**Fig. S6c**-AADAT expression levels and **Figs. S6d, e** malate levels). Significantly, re-expressing AADAT in 4T1 cells with a constitutive KD reversed malate elevation (**Figs. 2a bottom panel and 4i**), indicating AADAT’s direct role in regulating malate secretion in TNBC cells.

### AADAT regulates malate secretion by regulating CoQ_10_ redox homeostasis

Pathway analysis of the CM metabolome in E0771-ova+ cells with inducible *AADAT* KD, compared with uninduced control cells, revealed changes in amino acid metabolism, the TCA cycle, and CoA biosynthesis (**Fig. 5a**), suggesting mitochondrial stress. Consistent with this, E0771-ova+ cells with inducible AADAT KD and E0771 cells with constitutive AADAT KD showed significantly lower extracellular acidification rate (ECAR, **Figs. S6f-h**), oxygen consumption rate (OCR, **Figs. 5b,c, S6i**), and intracellular malate-to-oxaloacetate ratio (**Fig. 5d**, a surrogate for mitochondrial redox capacity ^33^ or NADH: NAD+ ratio, not seen in Dox control shown in **Fig. S6j**) compared to control cells. Moreover, unbiased metabolomic profiles of CM from *AADAT*-KD cells also revealed elevated malate levels, along with amino acids, including tyrosine, and decreased levels of TCA intermediates such as α-ketoglutarate (α-KG) and succinate, as well as the tyrosine breakdown product, 4-hydroxyphenylpyruvate, a necessary precursor for CoQ_10_ synthesis (**Fig.4a**). The pattern of decreased succinate and α-KG, along with reduced levels of the CoQ_10_ precursor 4-hydroxyphenylpyruvate, and decreased OCR seen with AADAT KD cells (**Figs. 5b, c**), suggests impaired mitochondrial oxidative metabolism, indicating TCA cycle stalling, disrupted electron transport, and redox imbalance. It also suggests that AADAT regulates CoQ_10_ levels, and its loss creates a redox bottleneck, due to reductive stress (**Fig. 5d**). These findings indicate a synergism between reversal of normal α-KG/malate antiporter activity due to high intracellular malate and elevated NADH/NAD+ levels caused by reduced CoQ_10_, promoting mitochondrial malate production via malate dehydrogenase 2 (MDH2), which leads to increased malate secretion. High levels of secreted malate create an immunostimulatory metabolic signal in the TME.

**Figure 5.**
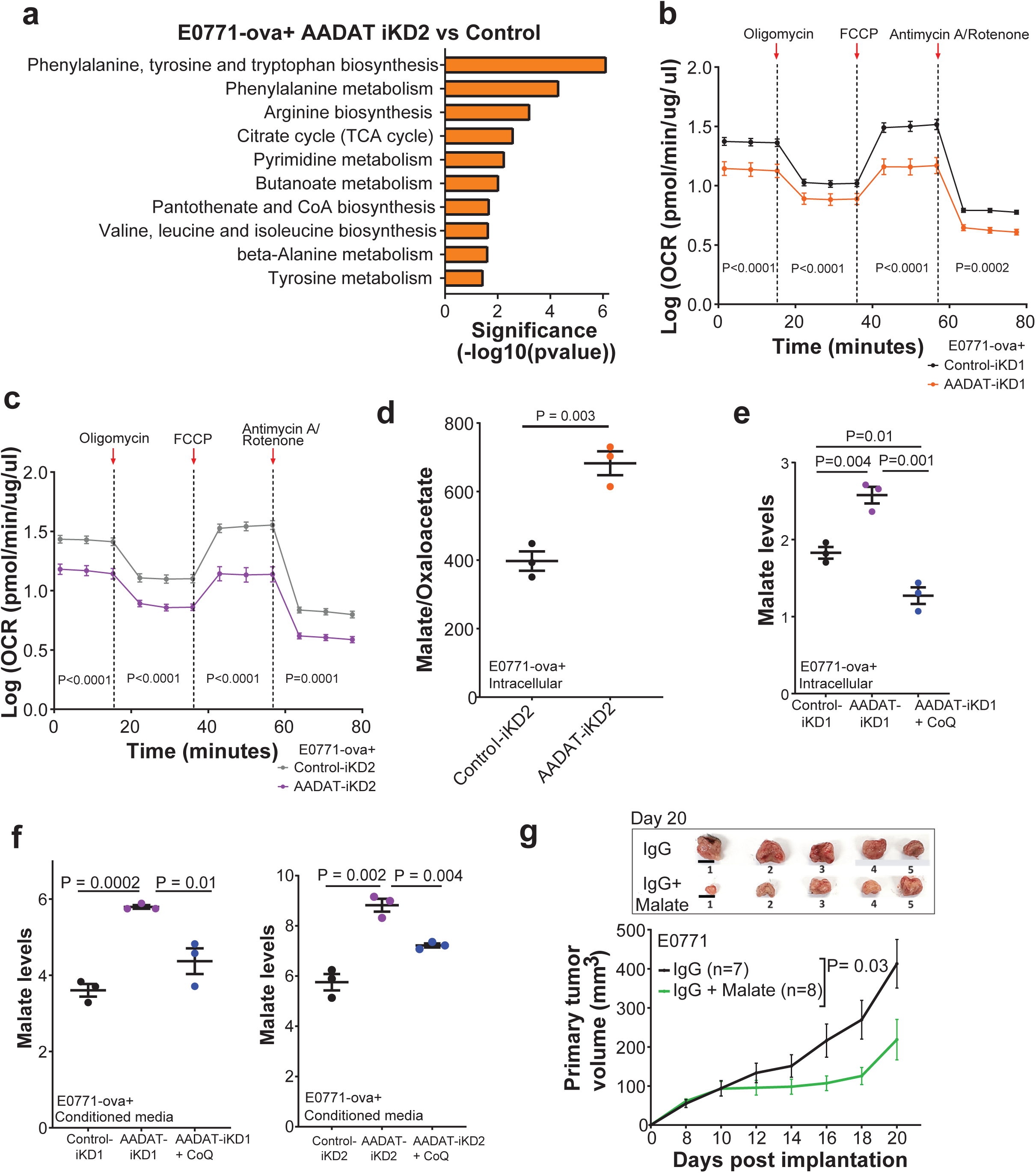
TNBC cells with AADAT knocked down exhibit altered mitochondrial activity that influences CoQ_10_ redox balance, leading to increased malate levels that suppress tumor growth. (a) Bar plot showing significantly enriched pathways (FDR<0.25) derived from the altered list of metabolites shown in panel a, comparing inducible knockdown of *AADAT* (AADAT-iKD2) with its matched uninduced control (Control-iKD2) in E0771-ova+ cells. (b,c) Line plot illustrating the oxygen consumption rate (OCR) measured with the Mito-Stress test on a Seahorse Bioanalyzer for E0771-ova+ cells with two independent doxycycline-induced *AADAT* knockdowns (AADAT-iKD1 and AADAT-iKD2), compared to their respective uninduced controls (Control-iKD1 and Control-iKD2). Three biological replicates were analyzed for each condition, each containing 9-10 technical replicates. P values were calculated using an unpaired two-tailed Student’s t-test. (d) Dot plot showing the Malate/Oxaloacetate ratio in E0771-ova+ cells with doxycycline-induced AADAT knockdown (AADAT-iKD2) versus uninduced controls (Control-iKD2). Three biological replicates were analyzed. P values were calculated using an unpaired two-tailed Student’s t-test. (e) Dot plot showing normalized intracellular malate levels in E0771-ova+ cells with doxycycline-induced AADAT knockdown (AADAT-iKD1), treated with or without Coenzyme Q10. (f) Same as in panel e, but in conditioned media from two independent doxycycline-induced AADAT knockdowns (AADAT-iKD1 and AADAT-iKD2). Three biological replicates were analyzed for each condition. P-values were calculated using an unpaired two-tailed Student’s t-test. (g) Tumor growth analysis of wild-type C57BL/6J mice orthotopically implanted with the syngeneic E0771 cell line treated with 5% malate (green, n=8) or IgG (black, control, n=7) in drinking water daily for 12 days, followed by 1% malate until the study endpoint. Statistical significance was determined using multiple unpaired Student’s t-tests. The indicated P value is for the last study time point, i.e., Day 20. The inset at the top shows representative photomicrographs of five tumors resected on Day 20 from each group. Scale bar: 1cm.

To confirm the link between CoQ_10_ levels and malate secretion in *AADAT* KD cells, we added CoQ_10_ to E0771-ova+ cells with inducible *AADAT* KD. As expected, CoQ_10_ supplementation reversed the increased intracellular and secreted malate levels observed with *AADAT* KD (**Figs. 5e, f, S6k)**.

We then investigated the impact of malate on TNBC tumor growth using C57BL/6J female mice orthotopically injected with E0771 cells. Malate supplementation in drinking water significantly reduced tumor growth in treated mice compared with untreated controls **(Fig. 5g),** demonstrating its tumor-reducing effects *in vivo*.

### Malate Recapitulates AADAT KD Effects on T Cell Activation

Based on our previous findings, we performed co-culture experiments to test the ability of activated CD8+ T cells isolated from OT-1 mice to kill murine TNBC cells (see methods section). The CD8+ T cells were then co-cultured with E0771-ova+ cells with or without inducible KD of *AADAT* (**Fig. 6a**). Additionally, we included a co-culture with uninduced E0771-ova+ and CD8+ T cells where the media was supplemented with 2.5 mM malate (**Fig. 6a**). Interestingly, CD8+ T cells showed significantly increased killing of E0771-ova+ cells with *AADAT* KD compared to uninduced controls (**Fig. 6b**). Furthermore, supplementation of malate also resulted in a similar level of CD8+ T cell killing on the E0771-ova+ control cells **(Fig. 6b).** Malate supplementation did not affect either tumor cells or T cells when cultured separately (**Figs. S7a-c**). Additionally, an unrelated metabolite like tryptophan did not alter the ability of CD8+ T cells to kill E0771-ova+ cells in a similar co-culture setup **(Fig. 7d).** Along the same lines, treatment with malate, but not fumarate, significantly enhanced the killing of patient-derived TNBC organoids (with baseline HER-2 levels) by HER-2-directed CAR-T cells, but not by non-targeting CAR-T cells (**Fig. S8).**

**Figure 6.**
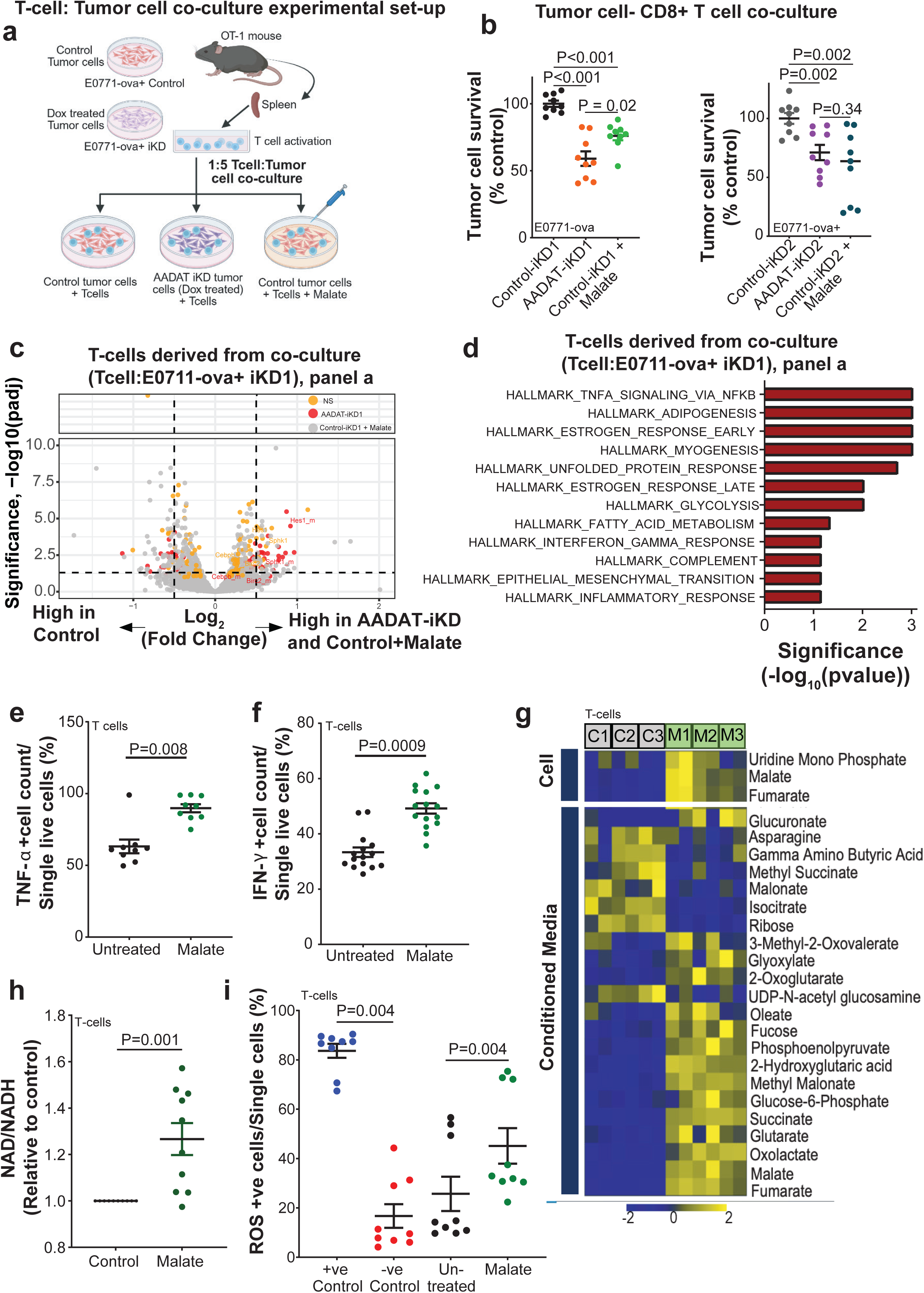
Malate enhances the inflammatory capacity of CD8+ T cells by boosting mitochondrial fitness. (a) Schematic of the co-culture showing CD8+ T cells from OT1-mice with E0771-ova+ cells that have an induced *AADAT* KD or uninduced controls. The uninduced controls are either treated with or without 2.5 mM malate. (b) The dot plot shows the percentage of surviving E0771-ova+ cells under conditions of induced *AADAT* KD or uninduced controls in a co-culture with CD8+ T cells derived from OT1-mice. The controls were either treated with or without 2.5 mM malate. Two independent inducible *AADAT* knockdowns (AADAT-iKD1 and AADAT-iKD2) are compared to their respective uninduced controls (Control-iKD1 and Control-iKD2). n=3 biological replicates, each with three technical replicates, were analyzed per condition. The P-value was calculated using a two-tailed unpaired Student’s t-test. (c) Volcano plot displaying differentially expressed genes (FDR<0.1, fold change ≥ 2) in CD8+ T cells from the co-culture with E0771-ova+-iKD1 cells (see panel b) and induced *AADAT* KD (yellow) or E0771-ova uninduced cells treated with 2.5 mM malate (red), compared to uninduced and untreated controls. Inflammatory genes are labeled. (d) Bar plot showing the results of hallmark pathways enriched by gene set enrichment analysis (GSEA, FDR<0.1) of differentially expressed genes in malate vs untreated CD8+ T cells, presented in panel b, red dots. (d) Bar plot showing common (e) Dot plot illustrating flow cytometry-derived intracellular expression of TNF-α in CD8+ T cells, treated with or without 2.5 mM malate. n=3 biological replicates, each with three technical replicates were analyzed for each condition. P-values were calculated using a two-tailed Wilcoxon matched-pairs signed rank test. (f) Same as in panel e, but for Interferon-γ. n=3 biological replicates, each with five technical replicates, were analyzed for each condition. P-values were calculated using two-tailed Wilcoxon matched-pairs signed rank test. (g) Heat map illustrating changes in mitochondrial metabolites (FDR<0.25, ≥ 1.5-fold) in CD8+ T cells treated with or without 2.5 mM malate. Three biological replicates, each with two technical replicates, were analyzed. Shades of yellow and blue indicate increased or decreased metabolite levels (see color scale). C1-C3: Control; M1-M3: Malate-treated. (h) NAD: NADH ratio in CD8+ T cells treated with or without 2.5 mM malate. Ten biological replicates were analyzed. P-values were calculated using a two-tailed unpaired Student’s t-test. (i) Reactive oxygen species (ROS) analysis with CD8+ T-cells divided into four groups: Pyocyanin (positive control) treated, N-acetylcysteine (negative control) treated, Untreated, and 2.5 mM Malate treated. n=3 biological replicates, each with five technical replicates were analyzed for each condition. P-values were calculated using two-tailed Wilcoxon matched-pairs signed rank test.

**Figure 7.**
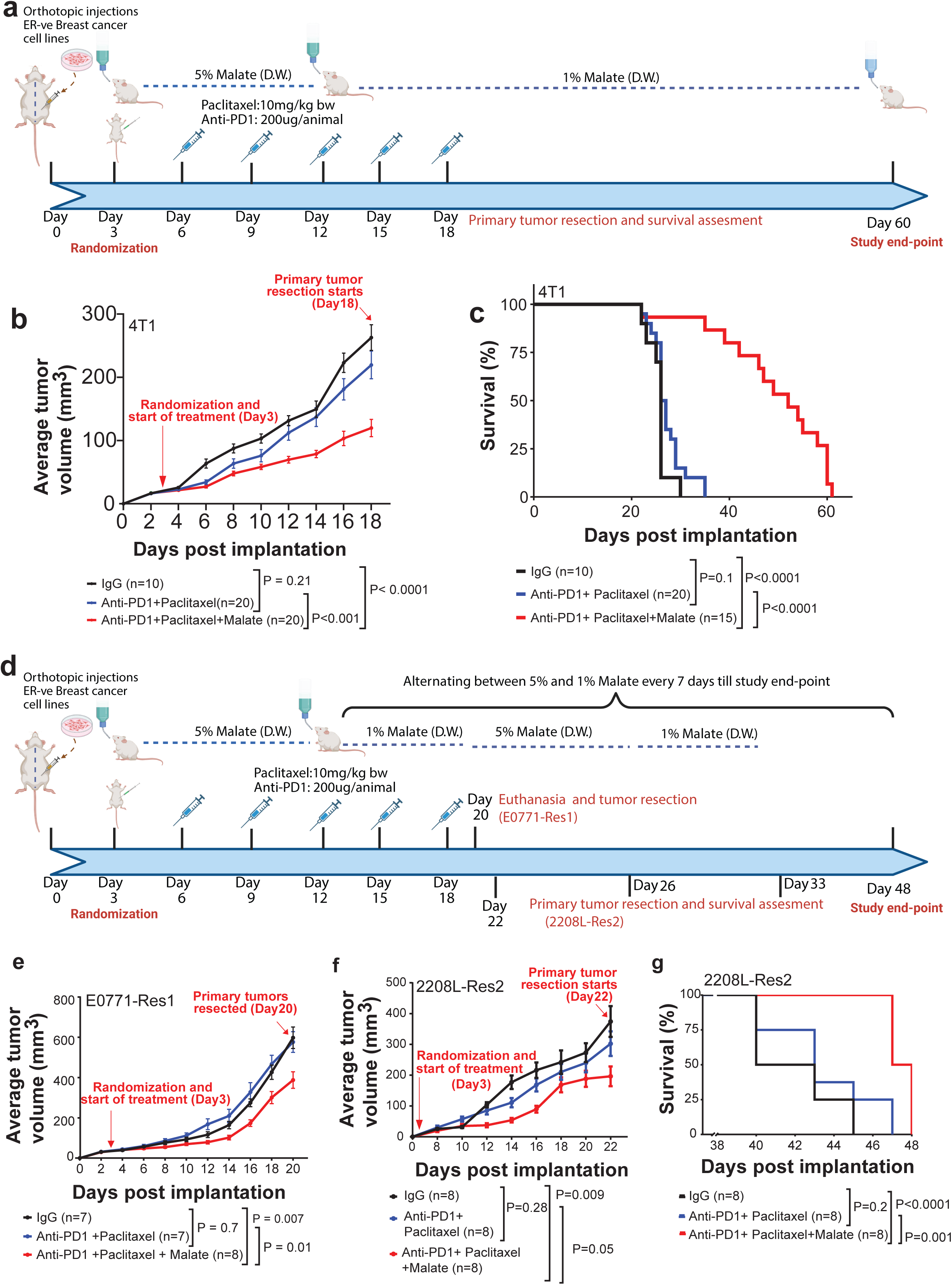
Malate enhances the efficacy of chemo-immunotherapy in vivo. (a) This schematic shows the experimental setup to study the effect of malate on the efficacy of PTX and anti-PD1 treatments in 4T1 tumors implanted in syngeneic wild-type BALB/cJ mice. Tumors were injected into the breast fat pad, and animals were randomized into four treatment groups on day 3, when tumors became palpable. The mice received three different regimens: 1) IgG (control), 2) PTX at 10 mg/kg with anti-PD1 at 200 μg/animal on every 3 days for six cycles (from day3 - day18), and 3) a combination of malate (5% for 12 days, followed by 1% until day 60), PTX, and anti-PD1 administered as mentioned above. Primary tumors were resected once they reached 300-350 mm³, beginning on day 18. Animals were monitored after resection with the endpoint defined by IACUC-approved early euthanasia criteria or in some cases unexpected deaths. (b) Tumor growth analysis of 4T1 cells implanted into syngeneic wild-type BALB/cJ mice, treated with IgG, PTX + anti-PD1, or the combination of malate, PTX, and anti-PD1 as described in panel a. Statistical significance was assessed using multiple unpaired Student’s t-test for each specified time point. The sample sizes are provided. (c) Kaplan-Meier survival analysis of the animals injected with 4T1 cells across the three treatment groups mentioned above was performed, with the endpoint defined by IACUC-approved early euthanasia criteria or, in some cases, unexpected deaths. Statistical significance is determined by the log-rank (Mantel-Cox) test. Sample sizes are indicated in the graph. (d) Similar to panel a, but for E0771-Res1 and 2208L Res2 cells that had acquired resistance to PTX and anti-PD1. The treatment doses and timing are the same as in panel a, except that malate levels were alternated weekly between 1% and 5% after the 12-day treatment period. The E0771 Res1 tumors were all resected on the same day (Day 20) following euthanasia. 2208 Res2 tumors were resected starting on Day 24, and the animals were monitored for post-surgical survival. (e) Tumor growth analysis of E0771-Res1 cells implanted into syngeneic wild-type B6 mice, treated with vehicle, malate, PTX + anti-PD1, or the combination of malate, PTX, and anti-PD1. The mice received three different regimens: 1) IgG (control); 2) PTX at 10 mg/kg with anti-PD1 at 200 μg/animal every 3 days for six cycles (from day3 - day18); and 3) a combination of malate (5% for 12 days, then 1% and 5% malate on alternate weeks until the end of the study), PTX, and anti-PD1 administered as mentioned above. Animals were euthanized, and all primary tumors were resected once the tumors in any group reached a volume of 600-650mm³, i.e Day20. (f) Tumor growth was analyzed in 2208L-Res2 cells implanted into syngeneic wild-type Balb/cJ mice. The treatment followed the same protocol as in panel e. Primary tumors were surgically removed once they reached 600-650 mm³, starting on day 24. Animals were monitored after resection, with the endpoint defined by IACUC-approved early euthanasia criteria or, in some cases, unexpected deaths. A multiple unpaired Student’s t-test was used to assess statistical significance at each specified time point. The sample sizes are provided. (g) Kaplan-Meier survival analysis of the animals injected with 2208L-Res2 cells across the three treatment groups mentioned above was performed, with the endpoint defined by IACUC-approved early euthanasia criteria or in some cases unexpected deaths. Statistical significance is determined by the log-rank (Mantel-Cox) test. Sample sizes are indicated in the graph.

To gain mechanistic insights, we performed bulk RNA-seq analysis of CD8+ T cells obtained from co-cultures described in **Fig. 6a**. As shown in **Fig. 6c**, CD8+ T cells exposed to either AADAT KD cells or malate-supplemented control cells displayed significantly altered, overlapping gene expression signatures. Gene Set Enrichment Analysis (GSEA) of this common set of altered genes revealed enrichment for inflammatory pathways, including those describing TNF-α signaling and the IFN-γ response (**Figs. 6d**). Consistent with this, elevated intracellular levels of both TNF-α and IFN-γ were confirmed by flow cytometry in CD8+ T cells treated with malate (**Figs. 6e, f**), suggesting a possible mechanism for malate-induced activation of CD8+ T cells.

To further assess whether malate-induced activation of inflammatory pathways in CD8+ T cells results from metabolic rewiring, we examined the metabolome of malate-treated CD8+ T cells. Compared to untreated CD8+ T cells, malate-treated T cells exhibited significantly increased glycolytic and TCA cycle metabolites, a higher NAD⁺: NADH ratio, and elevated levels of ROS (**Figs. 6g-i**), all of which indicate improved metabolic fitness that could enhance CD8+ T-cell effector function. Collectively, these data demonstrate that malate supplementation mimics AADAT KD by promoting CD8⁺ T cell effector function through enhanced metabolic fitness and activation of the inflammatory pathway, thereby driving anti-tumor immunity.

### Malate Enhances the Efficacy of Standard-of-Care Immunochemotherapy in TNBC Models

Given the association between lower AADAT expression and improved immunotherapy response in patient data, and our discovery that malate’s immunostimulatory effects enhance immunotherapy efficacy when AADAT is knocked down, we investigated whether malate supplementation could overcome resistance in TNBC tumors unresponsive to standard immunochemotherapy. To examine this, 5% malate was administered daily in drinking water for 12 days to BALB/cJ female mice bearing orthotopic de novo immunotherapy resistant 4T1 TNBC tumors, in combination with anti–PD-1 (200µg, intraperitoneal injection) and PTX (10 mg/kg, intra-orbital injection), given for six cycles every 3 days (**Fig. 7a**). On day 12 after tumor implantation, malate supplementation was lowered to 1% in both the malate-only group and the group receiving malate combined with chemo-immunotherapy until the study concluded **(Fig. 7a).** Remarkably, combining malate with anti–PD-1 and PTX significantly decreased tumor volume and increased survival compared to vehicle, or anti–PD-1 plus PTX treatment groups. (**Figs. 7b, c**).

Additionally, we conducted similar experiments with E0771 and 2208L cells that were made resistant to PTX and anti-PD1 in vitro, referred to as E0771-Res1 and 2208-L Res2, respectively ^34^. These cells were orthotopically injected into syngeneic wild-type BALB/cJ mice, which were treated with a similar regimen (**Fig. 7d**). However, malate levels after Day 12 were alternated weekly between 1% and 5% until the study concluded, as these cells were inherently resistant to chemo-immunotherapy agents **(Fig. 7d).** Importantly, like the results in the 4T1 model, the combination of malate with anti-PD1 and PTX resulted in significantly reduced tumor volumes in both these resistant models (**Figs. 7e, f**), and increased survival (**Fig. 7g**) compared to vehicle-treated or chemo-immunotherapy-treated controls. Furthermore, histopathological examination revealed that animals with 2208L-Res2 tumors treated with a combination of malate, anti-PD1, and paclitaxel exhibited no signs of toxicity in the liver or lungs. (**Fig. S9**).

Overall, these results indicate that malate reverses immunotherapy resistance, enhances both local tumor control and systemic anti-tumor immunity, while improving survival.

These findings collectively show that specific TNBC subgroups with high AADAT expression maintain a CoQ_10_-supported redox imbalance, which prevents the buildup and export of malate. Lower malate levels in the tumor microenvironment (TME) reprogram CD8+ T cell metabolism, leading to an exhausted phenotype (**Fig. 8**). Conversely, adding malate to TNBC tumors improves the effectiveness of standard immune and chemotherapy by enhancing T cell activation and infiltration, reducing tumor growth, increasing survival, and providing a clinically feasible approach to improve patient outcomes in TNBC (**Fig. 8**).

**Figure 8.**
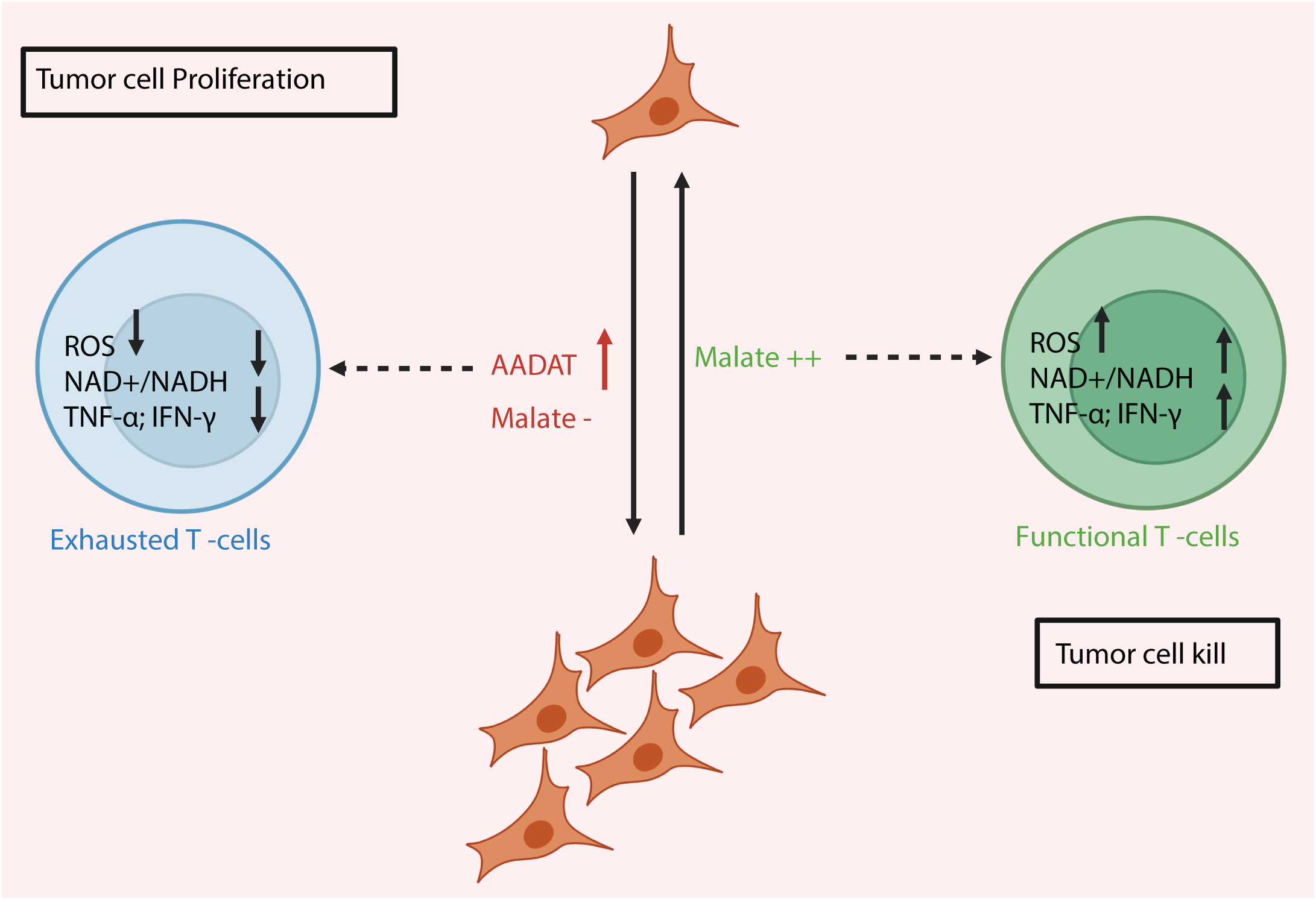
Schematic model describing the mechanistic basis of AADAT-induced TNBC prognosis suggested by our study. Overall, the study describes subtypes of TNBC that have elevated levels of the mitochondrial enzyme AADAT and secrete lower levels of the metabolite malate into the tumor microenvironment (TME), making it “cold” and immunosuppressed, characterized by a predominance of exhausted T cells. Supplementing exogenous malate in vivo or increasing intra-tumor malate levels in patient TNBC tumors are associated with a hot, immune-active tumor microenvironment, marked by a higher prevalence of functional T cells.

## Discussion

The aggressive nature, high recurrence rates, and limited actionable targets of TNBC continue to pose significant challenges despite advancements in immunochemotherapy, underscoring an urgent need for novel therapeutic strategies^35^. While immune checkpoint inhibitors (ICIs) like pembrolizumab, in combination with chemotherapy, have demonstrated survival benefits, a substantial proportion of patients still experience disease progression or recurrence^36^. This report highlights a novel immune-metabolic axis involving AADAT and malate, which significantly impacts the tumor immune microenvironment (TIME) and therapeutic response in TNBC. Our findings position AADAT as a critical driver of immune exclusion and identify malate as a potent, clinically translatable immunomodulatory adjuvant, offering a promising strategy to overcome resistance and improve outcomes in this challenging disease.

The kynurenine pathway (KP) of tryptophan metabolism is a well-established regulator of immune responses, frequently co-opted by tumors to promote immune evasion^37^. Canonical KP enzymes such as indoleamine-2,3-dioxygenase 1 (IDO1) and IDO2 are known to induce immune resistance by depleting tryptophan and generating immunosuppressive kynurenines, often correlating with an influx of T cells that are subsequently rendered dysfunctional or exhausted within the immunosuppressive TME^38^. This study unveils AADAT as a distinct and previously unappreciated player within the KP, exhibiting a unique inverse correlation with CD8⁺ T cell infiltration and poor prognosis in breast cancer, particularly TNBC. This stands in stark contrast to the positive correlation observed between IDO1, IDO2, and HAAO expression and CD8⁺ tumor-infiltrating lymphocytes (TILs). The established understanding of IDO1 and IDO2 suggests that they suppress the immune response by depleting tryptophan and producing immunosuppressive kynurenines, often correlating with an increase in TILs, which is interpreted as a compensatory immune response or an active, albeit suppressed, immune microenvironment^39^.

AADAT’s inverse correlation, which is uniquely linked to decreased CD8⁺ TIL infiltration, indicates a different mechanism from KP. This mechanism involves reprogramming tumor mitochondria, which subsequently leads to downstream events resulting in T cell exhaustion. This deviation from the canonical KP immunosuppression model represents a significant novelty for understanding immune evasion and designing targeted therapeutic interventions. The clinical relevance of AADAT in TNBC is further underscored by its elevated expression in malignant breast tumors, particularly in aggressive HER2-like and TNBC subtypes, and its consistent prediction of worse clinical outcomes across multiple patient datasets^40^. Given TNBC’s reliance on metabolic reprogramming and its aggressive nature with limited therapeutic targets, identifying AADAT as a key metabolic enzyme that promotes tumor growth and immune evasion positions it as a significant novel therapeutic target^35^.

Direct evidence of AADAT’s pro-tumorigenic role was established through genetic knockdown experiments, where depletion of AADAT in TNBC cell lines significantly delayed tumor formation and growth in immunocompetent mice. This tumor-suppressive effect was primarily mediated by CD8⁺ T cells, as AADAT knockdown did not delay tumor initiation in CD8 knockout mice. However, although established AADAT knockdown tumors grew more slowly, even in CD8 knockout mice, this suggests the presence of additional CD8-independent mechanisms. AADAT’s role in shaping the tumor immune microenvironment extends beyond CD8⁺ T cells, as evidenced by flow cytometry, which reveals increased infiltration of macrophages and CD4⁺ T cells following AADAT knockdown. Furthermore, inducible AADAT depletion after tumor establishment significantly reduced spontaneous pulmonary metastases post-resection. While many TME factors contribute to immunosuppression, AADAT’s unique correlation with reduced CD8⁺ TILs suggests it actively contributes to an “immune desert” phenotype, which presents a more challenging barrier to immunotherapy than an “inflamed but suppressed” TME ^41^. This implies that AADAT inhibition might be particularly effective in converting immunologically “cold” tumors to “hot” ones, thereby enhancing anti-tumor immunity. The translational relevance of AADAT inhibition in sensitizing TNBC to immunotherapy is supported by clinical and preclinical data, including a melanoma dataset (GSE91061) where low AADAT expression correlated with improved immunotherapy response^42^. Experimentally, AADAT depletion synergized with anti-PD-1 and anti-CTLA-4 combination therapy, leading to significantly reduced tumor growth. Imaging mass cytometry demonstrated increased interactions between tumor cells and CD8⁺ T cells in AADAT knockdown tumors, highlighting AADAT as a promising target to enhance the efficacy of immune checkpoint blockade (ICB) in TNBC, a critical need given the variable response rates to current ICB strategies^43^.

While AADAT is a primary enzyme in the kynurenine and lysine catabolism pathways, it also exhibits aminotransferase activity in the tyrosine metabolism pathway, underscoring the importance of considering this broader metabolic impact when interpreting our findings. This positions AADAT as a critical node where amino acid metabolism intersects with immune regulation in TNBC, aligning with the broader concept of metabolic vulnerabilities in cancer cells that hijack normal metabolic pathways to create an immunosuppressive environment^43^.

Unbiased metabolomics revealed a profound impact of AADAT on mitochondrial function, potentially via CoQ_10_, a critical cofactor in the electron transport chain and a potent antioxidant ^44^. Specifically, increased malate levels were observed in conditioned media following AADAT knockdown in TNBC cells, accompanied by impaired mitochondrial respiration, increased intracellular malate-to-oxaloacetate ratios (a surrogate for mitochondrial NADH: NAD+^33^), and decreased ATP production. Crucially, CoQ_10_ supplementation normalized malate levels in AADAT knockdown cells. The finding that AADAT directly regulates CoQ_10_ levels, thereby impacting mitochondrial function and immune evasion, represents a novel mechanistic link. While CoQ_10_’s role in mitochondrial health and immunity is well-established, and its deficiency has been linked to various diseases, including cancer^45^, identifying AADAT as an upstream regulator of CoQ_10_ provides a new target for modulating this critical pathway. Interestingly, studies have indicated that high CoQ pathway activity can correlate with worse prognosis and lower immune infiltration in breast cancer^46^, aligning with the current findings and suggesting that dysregulation of CoQ_10_ (e.g., AADAT-mediated reduction leading to compensatory malate secretion) might be beneficial for anti-tumor immunity. This suggests a complex interplay in which optimal CoQ_10_ levels are crucial and perturbing them in the context of AADAT inhibition may be a therapeutic option. The compensatory secretion of malate following AADAT knockdown and CoQ_10_ impairment suggests a metabolic adaptation by tumor cells, highlighting how disruption of one metabolic pathway leads to the extracellular accumulation of another metabolite (malate).

We believe that the metabolic adaptation leading to malate accumulation involves two separate processes, both regulated by AADAT. AADAT/KATII is known to produce α-KG via the TCA cycle by utilizing several glucogenic amino acids^25^. Knocking down AADAT could block α-KG from entering the TCA cycle and from reversing the activity of the α-KG/malate antiporter, thereby decreasing the import of cytosolic malate into mitochondria and increasing its export, both of which are limited by α-KG import. Additionally, as mentioned earlier, AADAT reduces CoQ_10_ levels, leading to reductive stress (high NADH/NAD), which may enhance malate production via mitochondrial Malate Dehydrogenase 2 (MDH2). These downstream effects of AADAT—namely, increased mitochondrial malate synthesis and decreased cytosolic malate import—could lead to a net buildup of intracellular malate, subsequently exported into the tumor microenvironment by tumor cells. Supporting this, our study shows that exogenous CoQ_10_ supplementation can normalize both intracellular and secreted malate levels in tumor cells. Future experiments measuring tumor CoQ_10_ levels and assessing the ability of external CoQ_10_ to alleviate reductive stress will be key in confirming this novel metabolic reprogramming, which results in increased malate secretion into the tumor microenvironment.

Strikingly, malate supplementation mimicked the anti-tumor effects of AADAT knockdown in immunotherapy-resistant preclinical models. Malate treatment reduced tumor growth and enhanced CD8⁺ T cell-mediated killing of TNBC cells in co-culture assays, establishing malate as a direct effector of the observed anti-tumor immunity. The specific role of malate was confirmed by the absence of CAR-T cell cytotoxicity when fumarate was used, which serves as a substrate for malate in the TCA cycle and was also secreted into the conditioned medium (CM) of AADAT KD containing E0771-Ova+ cells (**Fig. 4a**). Additionally, among the metabolites that were elevated in the CM of AADAT KD containing E0771-Ova+ cells, only malate showed overlapping spatial distribution with functional T cells in patient tumors. Furthermore, intra-tumoral malate levels were significantly linked to patient outcomes after chemotherapy. These findings prompted further investigation into the immunostimulatory properties of malate.

Bulk RNA-seq and flow cytometry revealed that malate treatment enriched for inflammatory pathways, including TNF-α and IFN-γ, and increased glycolysis, TCA cycle metabolites, the NAD⁺: NADH ratio, and reactive oxygen species (ROS) levels in CD8⁺ T cells ^47^. These metabolic shifts are consistent with enhanced effector function and metabolic fitness, which are critical for robust anti-tumor immunity^48^.

By demonstrating that AADAT knockdown in tumors impairs mitochondrial respiration (via CoQ_10_ reduction) and that malate can restore T cell metabolic fitness, the study uncovers a new upstream regulatory mechanism for T cell function, potentially preventing T cell exhaustion. This establishes a sophisticated, multi-step connection between the tumor’s metabolism and the immune cell’s function.

In a highly relevant preclinical setting, adding malate to drinking water significantly reversed resistance to standard treatment regimens that combine anti-PD-1 and PTX in TNBC tumors. Malate supplementation led to notable decreases in tumor size and enhanced survival in the orthotopic 4T1 model, which was de novo resistant to immunotherapy, as well as in the E0771-Res1 and 2208-L Res 2 TNBC models that had developed resistance to PTX and anti-PD-1 ^34^ Notably, our preliminary unpublished findings suggest that E0771-Res1 and 2208-L Res 2 follow distinct pathways to acquire resistance to immunotherapy. This demonstrates malate’s potential as a non-toxic, potent metabolic adjuvant to reverse anti-PD1 resistance by enhancing T cell function. This is evident in patient-derived TNBC tumors, which show a positive correlation between intra-tumoral malate levels and clusters containing functional T cells, and a negative correlation with clusters of exhausted T cells. Most importantly, TNBC patients with high intra-tumoral malate levels had improved 10-year survival following standard-of-care chemotherapy compared with those with low malate levels. Notably, malate has been shown to inhibit cancer growth in other contexts, for instance, by inhibiting 6-phosphogluconate dehydrogenase (6PGD) and DNA synthesis in lung cancer ^49^.

These findings position malate as a novel, clinically translatable adjuvant that may overcome resistance mechanisms in TNBC. It further supports malate’s role as a biomarker for a “hot” immune microenvironment and its functional contribution to anti-tumor immunity in patients. The findings directly address the significant challenges in TNBC treatment, characterized by limited therapeutic targets and a poor prognosis ^35^. By identifying AADAT as a novel target and malate as an effective adjuvant for immunochemotherapy, this research offers a promising new strategy to improve outcomes in this aggressive subtype, particularly for patients who may not respond optimally to current ICB plus chemotherapy regimens. This provides a new, potentially well-tolerated approach to enhance outcomes for a patient population that urgently requires improved therapies. Provocatively, it also establishes a basis for developing dietary strategies to address immunotherapy resistance in TNBC.

In summary, this study identifies AADAT as a novel, critical regulator of the tumor immune microenvironment in TNBC, driving immune exclusion and promoting tumor progression. A unique immune-metabolic axis is unveiled, in which AADAT modulates mitochondrial function via CoQ_10_, leading to compensatory malate secretion when AADAT is reduced. Crucially, the evidence demonstrates that malate acts as a potent immunomodulator, enhancing CD8⁺ T cell effector function and sensitizing immunotherapy-resistant TNBC to standard-of-care immunochemotherapy, ultimately improving survival in preclinical models. The robust clinical validation of intra-tumoral malate as a biomarker for immune activation and improved patient outcomes underscores the profound translational potential of these findings. This research provides a compelling rationale for targeting AADAT and for exploring malate supplementation as a novel, accessible, non-toxic, and effective adjuvant strategy to enhance anti-tumor immunity and improve clinical outcomes for patients with TNBC.

## Materials and Methods

### Patient cohorts

The NCI cohort, which includes matched transcriptomics and immunohistochemistry data from 67 breast cancer patients collected by the National Cancer Institute, as previously described, was analyzed to identify KP enzymes associated with immune cell infiltration [19]. AADAT protein expression in TNBC patients was measured by IHC (Rabbit Polyclonal Antibody, Cat No: TA317921 from Origene, Rockville, MD) and related to immune cell marker expression in the Baylor Scott & White Health (BSWH) dataset, collected as previously described, and in a commercially available tissue microarray (US Biomax Inc, Cat# BR2082a, Rockville, MD) [20]. Publicly available METABRIC and TCGA breast cancer datasets were obtained as described previously and analyzed to associate AADAT mRNA expression with clinical outcomes [20,21]. For all datasets, samples with missing data were excluded.

All human studies were conducted under protocols 300009407 (University of Alabama at Birmingham) and at Roswell Park Comprehensive Cancer Center, with patient consent obtained from Roswell Park. Tissue cores from 31 tumors were analyzed. Alongside self-reported race, we collected clinical data including receptor status, tumor grade, stage, chemotherapy history, and 10-year follow-up data. All tumors were obtained during surgery. Subsequently, in the adjuvant setting, all patients were treated at the same hospital following standard care protocols.

Tissues were deparaffinized in xylene and rehydrated through graded alcohols. For antigen retrieval, the slide was pressure-cooked for 10 minutes. Endogenous peroxidase activity was quenched with 3% hydrogen peroxide for 5 minutes. Slides were blocked with 3% goat serum, then incubated with AADAT antibody (10 µg/mL, Cat# TA331923, Origene, Rockville, MD) at room temperature for 1 hour in humidity chambers. The HRP-conjugated goat anti-mouse/anti-rabbit secondary antibody (Jackson Immunoresearch Laboratories Inc., West Grove, PA) was applied for 40 minutes. The antigen-antibody complex was visualized using diaminobenzidine (Sigma-Aldrich, St. Louis, MO) for 7 minutes. Slides were counterstained with hematoxylin (Sigma-Aldrich, St. Louis, MO). Positive controls were included in each staining run; negative controls were prepared by omitting the primary antibody. Slides were then dehydrated in alcohols, cleared in three xylene baths, and mounted with Permount (Sigma-Aldrich, St. Louis, MO). Histology scores were assigned by Dr. Chandandeep, a clinical breast pathologist, based on multiplying the intensity and extent of IHC staining.

### Cell lines and cell culture

E0771, E0771-Res1 and E0771-ova+ cells (gift of Dr. Xiang Zhang) were cultured in RPMI-1640 medium (Thermo Fisher, Waltham, MA) supplemented with 10 mM HEPES buffer (Thermo Fisher). 4T1 and 2208L-Res2 cells ^34^ (gift of Dr. Xiang Zhang) were grown in DMEM medium (Corning, Corning, NY). All media contained 10% Fetal Bovine Serum (Gibco, Cat # A52094-01) and 1% Penicillin/Streptomycin solution (Corning, Cat # 30-002-Cl). Cell lines were confirmed by Short-Tandem Repeat analysis at MD Anderson Cancer Center and cultured in a humidified incubator at 37°C with 5% CO2. All cells were routinely checked for mycoplasma contamination using the MycoAlert Detection Kit (Lonza, Cat# LT07-418). To generate stable Aadat knockdown, E0771 and 4T1 cells were infected with lentiviral vectors containing two independent shRNAs targeting this gene, obtained from the BCM Cell-based Assay Screening Service (C-BASS, Houston, TX). To re-express Aadat, E0771 and 4T1 cells were infected with lentiviral vectors expressing the Aadat open reading frame (GeneCopoeia, Cat# EX-Mm06255-Lv151). In all cell lines, selection for knockdown was performed with puromycin (0.5 μg/ml for E0771 cells, 1 μg/ml for 4T1 cells), or for re-expression with neomycin (1 μg/ml), for at least one week. Inducible Aadat knockdown in E0771, E0771-ova+ and 4T1 cells was achieved using a lentiviral vector containing doxycycline-inducible shRNA targeting Aadat (C-BASS) Cells were induced with 2 μg/ml doxycycline and 4 μg/ml puromycin. shRNA details are mentioned in **Supplementary Table 5**. All the cell lines used in this study to generate genetic knockdown of AADAT and its re-expression are summarized in **Fig. S10**.

### RNA extraction and qRT-PCR

Procedures were performed as described in our previous publication^50^. Primers used in this study are listed in **Supplementary Table 6**.

### Metabolomics to analyze Kynurenine/Kynurenic acid ratio

Three replicates of 0.3 x 10^6 E0771 cells were plated and cultured overnight in standard media. The cells were then incubated with media containing 200 µM L-Tryptophan (13C(U); 15N2, Cambridge Isotope Laboratories, Cat# CNLM-2475-H, Tewksbury, MA) for 24 hours. After this period, cells were washed with PBS and stored at −80°C. Metabolites were extracted using a liquid-liquid method and analyzed by high-throughput LC-MS/MS as previously described (23,24). The tryptophan pathway metabolites were separated on a ZORBAX Eclipse XDB C-18 HPLC column under positive ionization mode. For ESI positive mode, mobile phases were 0.1% formic acid in water (A) and acetonitrile (B). The gradient flow was: 0-2 min 2% B; 2-12 min 10% B; 12-17 min 80% B; then re-equilibrated until 25 min back to 2% B. The flow rate was 0.3 ml/min, and the injection volume was 10 µL. Data acquisition was performed via multiple reaction monitoring (MRM) with a 6495 Triple Quadrupole MS coupled to an HPLC system (Agilent Technologies, Santa Clara, CA) using Agilent Mass Hunter Software (25). Peak integration and data analysis were carried out with Agilent Mass Hunter Quantitative Analysis software. The total metabolite pool was calculated by summing labeled and unlabeled fractions for each metabolite.

### Targeted metabolomics for malate and oxaloacetate in tumor cell-conditioned media

TCA cycle metabolites, including malate and oxaloacetate, were extracted from cell culture media, with pooled samples serving as quality controls, following the previously described extraction protocol^50–52^. Malate was separated using a Luna 3 µm NH2 (100 Å) HPLC column (Phenomenex, Torrance, CA), with mobile phases consisting of 5 mM ammonium acetate in water (A) at pH 9.9 and 100% acetonitrile (B). Metabolite separation was carried out on an Agilent 1290 Infinity HPLC system, with data collected on an Agilent 6495B Triple Quadrupole mass spectrometer (Agilent Technologies, Santa Clara, CA) in MRM mode with negative ionization. Data analysis involved Agilent Mass Hunter Quantitation software, including manual verification of integrations and peak evaluation across samples. Results were normalized using spiked isotopically labeled internal standards and log2-transformed. Differential metabolites were identified using the Benjamini-Hochberg method with an FDR cutoff of 0.25, following Student’s t-test.

### Untargeted metabolomics in tumor cell-conditioned media

For untargeted metabolomics, three volumes of methanol/acetonitrile (50/50, v/v) were added to cell culture media. Samples were vortexed for 5 min, kept at −20°C for 10 min, then centrifuged at 4°C and 15,000 rpm for 10 min. Extracts were dried using a GeneVac EZ-2 Plus SpeedVac and reconstituted in 100 µL of methanol/water (50/50, v/v). 10 µL of supernatant from each sample was pooled to create QC samples. Separation was performed using a Thermo Scientific Vanquish Horizon UHPLC system. HPLC analysis employed a Waters ACQUITY HSS T3 reversed-phase column for reverse-phase separation and a Waters ACQUITY BEH amide HILIC column for HILIC separation. Reversed-phase mobile phases were 0.1% formic acid in water (A) and methanol (B). HILIC mobile phases were 0.1% formic acid in ammonium formate, with varying solvent compositions. Gradient flow: 0–3 min at 1% B; 4–11 min at 50% B; 12–21 min at 95% B; re-equilibration for 30 min. Columns operated at 50°C, with a 300 μL/min flow rate and a 2 μL injection volume. Data acquired on a Thermo Orbitrap IQ-X and processed with Compound Discoverer 3.3.3.2 for peak detection and ID, using retention time, MS/MS databases, mzCloud, NIST 2020 HRMS, and HMDB (<5 ppm). QC runs every 10 samples as a reference. Data analysis included log2 transformation and median IQR normalization. Differential metabolites identified via Student’s t-test (p-values) and FDR correction (FDR<0.25).

### Statistical analysis for unbiased metabolomics

After data acquisition, the missing values for metabolites were imputed using the K nearest-neighbor (KNN) method. Then the data were log2 transformed and followed by the median IQR normalization. The compound-by-compound t-test was applied to identify the top differentially regulated metabolites that passed the nominal P value of p<0.05, followed by the Benjamini-Hochberg procedure for false discovery rate (FDR<0.25) correction, accounting for multiple comparisons ^53^.

### Pathway Enrichment for Differential Metabolites

We performed a pathway enrichment analysis via hypergeometric enrichment for core-enriched metabolites against Hallmark gene sets in the Molecular Signatures Database (MSigDB). A nominal *P*<0.05 and a false discovery rate (FDR) <0.1% were used as thresholds for significance of Enrichment.

### Proteomics

Cell pellets were lysed in 50 mM ammonium bicarbonate with 1 mM CaCl 2 through 3 minutes of sonication. Then, 25 μg of protein was digested with 0. 5 μg of trypsin/Lys-C for 12 hours at 37 ° C. Tryptic peptides were fractionated using a homemade reverse-phase C 18 column in a pipette tip. The peptides were eluted and separated with a stepwise gradient of increasing acetonitrile (2%, 4%, 6%, 8%, 10%, 12%, 14%, 16%, 18%, 20%, 22%, 24%, 26%, 28%, 30%) at pH 10. These fractions were combined into five groups (2 + 12 + 12, 4 + 14 + 24, 6 + 16 + 26, 8 + 18 + 28, 10 + 20 + 30) and vacuum dried. The dried peptides were analyzed on Easy-nLC 1000 nanoflow HPLC coupled with Orbitrap Fusion mass spectrometers (Thermo Fisher Scientific). The setup included a 2 cm × 100 μm trap column and a 5 cm × 150 μm separation column packed with 1. 9 μm Reprosil-Pur Basic C 18 beads. Peptides were separated using a 75-minute discontinuous gradient of 4–26% acetonitrile with 0. 0.1% formic acid at an 800 nl/min flow rate. The mass spectrometer operated under Xcalibur software version 4.1. 1, combining data-dependent acquisition (DDA) and Parallel Reaction Monitoring (PRM). In DDA mode, the top 3 peaks in the full MS scan (300–1400 m/z) at 120, 000 resolution were selected for fragmentation. MS/MS spectra via higher-energy collisional dissociation (HCD) were obtained in rapid scan mode in the ion trap. For PRM, seven specific AADAT peptides were monitored: EIYELAR (+ 2, m/z 447. 447.2463), FEDDLIKR EIYELAR (+ 2, m/z 581. 581.2844; + 3, m/z 345. 859), FLTATSLAR (+ 2, m/z 490. 490.2891), GLAEWHVPK (+ 2, m/z 518. 518.7893), SAAFTVENGSTIR (+ 3, m/z 451. 451.5726), TTADILSK (+ 2, m/z 424. 424.7461), QLIEEK (+ 2, m/z 380. 380.2205). The MS/MS spectra were searched against the Kynurenine/alpha-aminoadipate aminotransferase (UniProtKB/Swiss-Prot: Q 9 WVM 8. 1) protein sequence in Proteome Discoverer 2. 1 (Thermo Fisher), using the Mascot algorithm (Mascot 2. 4, Matrix Science). Allowed parameters included a precursor mass tolerance of 20 ppm, fragment mass tolerance of 0. 5 Da, up to two missed cleavages, and dynamic modifications such as N-terminal acetylation and methionine oxidation. PRM target peptides were manually validated.

### Mice

C57Bl/6J (WT, B6) were obtained from the Center for Comparative Medicine at BCM. B6.129S2-Cd8a^tm1Mak^/J (B6.CD8KO) were a gift from Dr. Xiang Zhang and bred in our facilities. BALB/cJ mice were obtained from Jackson Laboratory. CD8KO BALB/cJ mice were a gift from Dr. Xiang Zhang and bred in our facilities.

### Breast tumor models

Orthotopic tumors were generated by injecting 50 μl of breast cancer cells (0.5 x 10^6 E0771, 0.2 x 10^6 4T1, 1.5 × 10^5 E0771-Res1, and 1.2 x 10^5 2208L-Res2) in PBS mixed with 50 μl of Matrigel (Corning, Cat# 354248, Corning, NY) into the fourth mammary fat pad of mice. Bilateral tumors were established for tumor take and growth studies. Tumor growth was monitored three times weekly using calipers, and tumor volume was calculated with the formula V = π/6 × length × width^2.

### In-vitro and In-vivo treatments

The in-vitro treatments used in this study include Malate (or L-(-)-Malic acid, Sigma, Cat no. M7397), N-Acetyl-L-cysteine (NAC, Sigma, Cat no. A9165) and CoQ_10_ (Sigma-Aldrich, Cat no. C9538). Throughout various experimental setups, cells were treated with 2.5 mM Malate, 2.5 mM NAC and 10µM CoQ_10_ for 24 hours unless otherwise specified. For Malate and NAC treatment, a 100mM stock solution of Malate and NAC was prepared by dissolving the commercially available powdered form in molecular biology grade water at room temperature. The final concentration of 2.5mM was achieved by adding the required volume of stock solution in prewarmed media. For CoQ_10_ treatment, a 1 mM CoQ_10_ stock solution was prepared by dissolving CoQ_10_ in 100% ethanol and heating for 5 min to 65°C. 10μl of 1 mM CoQ_10_ was then added to 90 μl Intralipid (Sigma, Cat no.-I141) and heated to 65°C for 5 min. This solution was then added immediately to the cells to achieve a final concentration of 10 µM. In vivo studies involved treating mice with Doxycycline (Dox), anti-PD-1, anti-CTLA-4, PTX, and Malate. These treatments were administered separately or in combination, depending on the study aims. Dox was used to induce shRNA targeting Aadat in breast tumors by providing mice with water containing 2 mg/mL Dox, prepared twice weekly. For immune checkpoint blockade, each animal received intraperitoneal injections of 100 μg anti-CTLA-4 (clone 9D9) and 200 μg anti-PD-1 (clone RMP1-14), or equivalent amounts of isotype-matched control antibodies (mouse IgG2b (clone MPC-11), rat IgG2a (clone 2A3), BioXCel, Lebanon, NH) at the indicated times post-implantation. PTX was administered at 10 mg/kg body weight via intravenous retro-orbital injection. The injectable PTX was prepared by dissolving it in ethanol to achieve a concentration of 20 mg/mL, then mixing it with an equal volume of Cremophor® EL (Millipore, Cat no. 238470) to reach a final concentration of 10 mg/mL. The stock solution was stored at −20°C. On the day of injection, a 2 mg/mL solution was prepared in 1X PBS, and 100 µL of this working solution was injected per animal on the specified days. Five percent (w/v) Malate (Sigma, Cat no. M7397) was provided in drinking water starting from Day 3 of the study (Day 0 being the day of tumor cell injection into the mammary fat pad). The drinking water was replaced with a fresh 5% Malate solution every 24 hours for each cage. For the 4T1 and E0771 orthotopic mouse models, animals were maintained on 1% (w/v) Malate water until the study endpoint, after 12 days of treatment with 5% Malate water. For the E0771-Res1 and 2208L-Res2 orthotopic mouse model, animals alternated between 1% and 5% Malate water weekly until respective study endpoints, after 12 days of continuous 5% Malate water treatment.

### Immunofluorescence analysis

Frozen tumor sections were prepared as previously described (18). Primary antibodies against CD31 (1:100, R&D Systems, Cat# AF3628, Minneapolis, MN), VE-Cadherin (1:100, R&D Systems, Cat# AF1002, Minneapolis, MN), and NG2 (1:200, Sigma-Aldrich, Cat# AB5320, St. Louis, MO) were incubated with the samples overnight at 4°C. Subsequently, donkey anti-goat IgG, Alexa Fluor 488 (1:500, Jackson ImmunoResearch, West Grove, PA) or donkey anti-rabbit IgG, Alexa Fluor 555 (1:500, Invitrogen, Waltham, MA) were applied for 2 hours at room temperature prior to imaging.

### Flow cytometry of resected tumors

After resection, tumors were dissociated into single cells using the mouse Tumor Dissociation Kit (Miltenyi Biotec, Cat# 130-096-730, Auburn, CA). Cells were incubated on ice for 10 minutes with CD32 FcR blocker (1:100, Tonbo, clone 2.4G2, San Diego, CA). Subsequently, they were stained with conjugated antibodies for 25 minutes on ice in the dark using FACS buffer (PBS with 1% FBS), then washed twice with the same buffer. Data acquisition was performed on a BD LSR Fortessa, and analysis was done with FlowJo v10.0. To quantify absolute numbers of tumor-infiltrating immune cells, liquid counting beads (BD Biosciences, Franklin Lakes, NJ) were employed. The antibodies used, diluted 1:200, are listed in **Supplementary Table 7**.

### Seahorse XF Mito Stress Assay

Mitochondrial respiration was evaluated using the Seahorse XF Analyzer (Agilent Technologies) according to the manufacturer’s protocol. E0771-ova+ control and AADAT-iKD cells (both iKD1 and iKD2) were seeded in 9-10 wells per experimental run. A total of 3 runs were performed with cells taken from different passages. For each well, 20,000 cells were seeded in 100 µL of complete growth medium per well in an XF Cell Culture Microplate and incubated for 24 hours at 37 °C. The sensor cartridge was hydrated overnight in XF Calibrant at 37 °C in a non-CO₂ incubator. On the day of the assay, cells were washed and equilibrated with Seahorse assay medium (XF Base Medium supplemented with 1 mM sodium pyruvate, 2 mM glutamine, and 10 mM glucose) and incubated for 45–60 minutes at 37 °C in a non-CO₂ incubator. Oligomycin (1 µM), FCCP (1 µM), and rotenone/antimycin A (0.5 µM) were prepared in assay medium and loaded into Ports A, B, and C, respectively. Following calibration, the assay was performed according to standard procedures. Data were normalized with protein concentration per sample (µg/µl) in Wave 2.6.0. Data were analysed using Wave software (Agilent Technologies).

### T-cell kill assay

Mouse spleen was isolated after sacrificing a 6–8-week-old female OT-1 mouse obtained from The Jackson Laboratory (C57BL/6-Tg(TcraTcrb)1100Mjb/J, Strain #: 003831). The tissue was immediately washed with 1X PBS supplemented with 2% FBS and 1% Penicillin-Streptomycin (hereafter referred to as FACS buffer). Mechanical disruption of the tissue was performed to obtain a single-cell suspension with FACS buffer. The cell suspension was then filtered through a cell strainer (pluriSelect USA pluriStrainer Mini 10 µm (Cell Strainer), Cat. No. 43-10010-60) and collected in a 5 ml polystyrene tube. The strained cell suspension was centrifuged at 500g for 5 minutes at room temperature. The pelleted cells were incubated in 1X RBC lysis buffer (TONBO biosciences, Cat No. TNB-4300) for 10 minutes at room temperature. After another centrifugation under the same conditions, FCδ receptors were blocked by adding Purified Anti-Mouse CD16 / CD32 (FC shield) from Cytek Biosciences, Cat. No. 70-0161-U500. Negative selection of CD8+ T cells was performed by incubating the cells with a 1:100 dilution of biotin-labeled anti-mouse CD4 (TONBO Biosciences Cat. # 30-0041-U100), CD11c (TONBO Biosciences Cat. # 30-0114-U100), CD11b, CD45R, Ly-6G, and Ly-6C antibodies (BD PharmigenTM, Biotin Mouse Lineage Panel, Cat. No. 559971) at 4°C for 15 minutes. This was followed by centrifugation and resuspension of the pellet in 2 ml of FACS buffer with a 1:40 dilution of Streptavidin Particles Plus-DM (BD IMagTM, Cat. # 557812). Magnetic separation of these beads finally yields a CD8+ T cell population in the cell suspension. The cells were plated with growth media and T cell activation beads (25 µl per 10^6 T cells) (Thermo Fisher Scientific, DynabeadsTM Mouse T-Activator CD3/CD28, Cat. No. 11456D). The T cell growth media consisted of 9 ml RPMI, 1 ml heat-inactivated FBS, 40 µl 1:1000 β-mercaptoethanol, 4 µl of 10 ng/µl IL-2, and 100 µl Penicillin-Streptomycin solution. After 48 hours of activation, activated CD8+ T cells were co-cultured with tumor cells (1:5 ratio) and/or with treatment conditions (as detailed in the respective result sections and figures). After 24 hours of co-culture and treatment (where applicable), the entire well of co-cultured cells was harvested for flow cytometry analysis to assess tumor cell survival and T cell populations by staining with 1:200 dilutions of CD45 (Biolegend, Cat #103155), CD8 (Cytek, Cat # 35-0081-U500), and CD3e antibodies (Biolegend, Cat # 155703). Liquid counting beads (CountBrightTM Plus Absolute Counting Beads, Invitrogen, Cat. No. C36995) were used to quantify the absolute number of tumor and CD8+ T cells. Data were acquired using a BD LSR Fortessa and analyzed with FlowJo v10.0.

### PDXO and CAR T-cell co-culture

Breast Cancer PDXO Establishment: Preparation of patient-derived xenograft organoids (PDXOs) was achieved using 10-15 tumor fragments measuring 2-3 mm^3^ in 5 mL of warm advanced DMEM/F12 medium supplemented with Miltenyi tumor dissociation kit enzymes (Miltenyi Biotec, Auburn) and 10 mM Y-27632. Dissociation was performed using a GentleMACS Octo Dissociator with heaters (Miltenyi Biotec, Auburn) using the pre-set tough tumor type pre-set TDK program. Organotypic fragments were collected by centrifugation at 300 x g for 3 minutes. Fragments were washed in advanced DMEM/F12 medium and centrifuged to increase the organoid to single cell ratio as described in Scherer et al. 2023 (PMID: 37402170). Organoids were counted and embedded in organoid Matrigel (Corning, Glendale). PDXOs were maintained in 6-well plates (Genesee Scientific, El Cajon, CA, USA) and cultured in 200-μl domes of Matrigel matrix for organoid culture (Corning, Corning, NY, USA). PDXOs were covered with Advanced DMEM/F12 (Thermo Fisher, Waltham, MA, USA) supplemented with 5% FBS, 10 mM HEPES (Thermo Fisher), 1X Glutamax (Thermo Fisher), 1 μg/ml hydrocortisone (Sigma-Aldrich, Burlington, MA, USA), 50 μg/ml gentamicin (Genesee Scientific), 10 ng/ml hEGF (Sigma-Aldrich), and 10 μM Y-27632 (Selleck Chemicals, Houston, TX, USA). Organoids were considered established once they were authenticated though STR analysis, had a stable doubling time, and were devoid of stromal cells.

We obtained healthy donor peripheral blood mononuclear cells (PBMCs) through an institutional review board (IRB)-approved protocol at Baylor College of medicine. PBMCs were isolated using Lymphoprep according to manufacturer’s instructions (Axis-Shield PoC AS, Dundee, Scotland). T cells were activated in 24-well non-tissue culture-treated plates coated with OKT3 (1 mg/mL; Ortho Biotech, Bridgewater, NJ) and NA/LE anti-human CD28 antibodies (1 mg/mL; BD Biosciences, San Jose, CA) at a density of 1 × 10^6^ cells per well. H84T CAR-T cells post transduction were expanded in conventional CTL media (45% RPMI-1640 media (Hyclone Laboratories, Marlborough, MA), 45% Click’s medium (Irvine Scientific, Santa Ana, CA), 10% heat-inactivated FBS (Hyclone Laboratories, Marlborough, MA) and 2 mmol/ glutaMAX (Gibco by Life Technologies, Carlsbad, CA)) supplemented with 10ng/ml recombinant human IL-7 and IL-15 cytokines. Retroviral transduction was performed as previously described^54^. The vector encoding the HER.2 directed CAR (2^nd^ generation HER2.28z; clone FRP5) was a kind gift from Dr. Stephen Gottschalk. Transduction efficiency was measured by flow cytometry using a chimeric Erb2-Fc fusion protein (R&D Systems, Minneapolis, MN) with AF-647 anti-Fc antibody (Southern Biotech, Birmingham, AL).

Organoids were expanded prior to seeding in a microplate by culturing approximately twelve 75 ml domes in a 10 cm dish at a density of 1 × 10^^5^ – 2 × 10^^5^ cells per dome. Following 7 days of growth, organoids were collected by dispase (50 units/mL) solution (i.e. Dispase stock supplemented with 20% FBS and 10 mM Y-27632) treatment for 10-15 minutes at 37 °C. Dispase solution was washed off, and organoids were strained through a 100 mm strainer to ensure uniform size prior to plating. 500 organoids were seeded in suspension alone or in co-culture with HER2-directed CAR T or non-transduced T cells (ratio of 1 organoid to 50 T cells) stained with CFSE (ThermoScientific, Cat C34554) into 96-well plates pre-coated with a 15 mL base layer of Matrigel in a volume of 100 mL complete medium supplemented with 5% Matrigel. Complete medium was supplemented with either 2.5 mM Malate or Fumarate and their corresponding vehicle controls (i.e. water or ethanol, respectively). Growth was tracked using green integrated intensity and quantified over time using the IncuCyte software.

### T-cell RNA Sequencing and Analysis

The CD8+ T cell pellets were harvested from the 1:5 T cell: tumor cell co-culture as described in the ‘T cell kill activity’ methodology section. The cells were then washed with Phosphate Buffer Saline (PBS). Three million cells were harvested and counted per replicate (n=3). RNA extraction was performed using the Aurum™ Total RNA Mini Kit (Bio-Rad). Libraries were prepared using the ABclonal mRNA-seq Library Prep Kit for Illumina (poly (A) selection). RNA integrity and quantification were confirmed on the AATI Fragment Analyzer and quantitative PCR. Quality assessment and sequencing were performed by Novogene Corporation, Sacramento. Sequencing was performed using Illumina NovaSeq X Plus (PE150).

### RNA-seq Data Processing and Differential Expression Analysis

First, FAstQC was used to assess the quality of raw RNA-seq reads (https://www.bioinformatics.babraham.ac.uk/projects/fastqc/). It was followed by trimming of low-quality bases and adapter sequences using Trim Galore (https://github.com/FelixKrueger/TrimGalore). Cleaned reads were then aligned to the human reference genome (GRCm38/mm38) using HISAT2 ^55^ with default parameters. The resulting SAM files were converted to BAM format, sorted, and indexed using SAMtools ^56^. Gene-level read counts were quantified using featureCounts ^57^ from Subread. Only uniquely mapped reads overlapping exons were counted toward gene-level expression estimates. The raw count matrix was subsequently imported into DESeq2 ^58^ in R for normalization and differential expression analysis. DESeq2’s default pipeline, which employs a negative binomial generalized linear model, was used to estimate size factors and dispersions, followed by Wald tests for pairwise comparisons between experimental conditions. P-values were adjusted for multiple testing using the Benjamini–Hochberg false discovery rate (FDR) method. Genes with an adjusted p-value (FDR) < 0.10 and |log₂ fold change| ≥ 1 were considered significantly differentially expressed.

### Reactive Oxygen Species (ROS) analysis

The ROS analysis of T cells, which are suspension cells, was performed using the ROS-ID® Hypoxia/Oxidative Stress Detection Kit from Enzo (Cat no.-ENZ-51042). The T cells were harvested from the spleen of 8-10-week-old female C57Bl/6J (WT, B6) mice and activated as described in the ‘T cell kill assay’ section. The activated T cells were then cultured under four conditions: positive control (Pyocyanine treatment at a later stage), negative control (2.5 mM N-Acetyl Cysteine), untreated, and Malate-treated (2.5 mM Malate). The negative control and Malate-treated cells were grown in 2.5 mM NAC and Malate, respectively, for 24 hours. Simultaneously, the untreated and positive control cells were cultured in T-cell medium. After 24 hours, the cells were centrifuged at 400 x g for 5 minutes and the supernatant was removed. Cells from all culture conditions, except the positive control, were then resuspended in 200 µl of ROS-ID® Hypoxia/Oxidative Stress Detection Mix (6 µl of the oxidative stress reagent provided in the kit in 10 ml of 1X PBS) and incubated under standard cell culture conditions (37°C, 5% CO2) for 3.5 hours. The positive control cells were centrifuged and incubated in 200 µl of detection mix along with 500 µM Pyocyanin under the same conditions for 30 minutes. After the incubation, the cells were centrifuged at 400 x g for 5 minutes to remove the detection mix (and Pyocyanin, for positive control), then washed in 1X PBS. Finally, the cells were resuspended in 500 µl of 1X PBS and analyzed by flow cytometry. Data were acquired using BD LSR Fortessa and analyzed with FlowJo v10.0. ROS detection was performed using a filter set compatible with Fluorescein (Ex/Em: 490/525 nm).

### NAD+/NADH assay

The T-cells were harvested from the spleen of C57Bl/6J (WT, B6) 8-10-week-old female mice and activated as described in the ‘T cell kill assay’ section. The activated T-cells were then cultured under two conditions: untreated and malate-treated (2.5 mM malate) for 24 hours. NAD/NADH levels of these cells under the respective treatment groups were measured using the NAD/NADH-Glo™ Assay kit (Promega, Catalog no. G9071). 1 × 10^4 cells were plated per well in 50 µl of the respective culture media in a 96-well white-walled tissue culture plate. Then, 50 µl of NAD/NADH-Glo™ Detection reagent (prepared as instructed in the kit) was added to the wells containing the cells. The plate was gently shaken to mix and lyse the cells. It was then incubated for 60 minutes at room temperature, and luminescence was measured using a luminometer.

### T-cell cytokine analysis

The levels of T-cell intracellular cytokines (TNF-α and IFN-δ) were measured using flow cytometry. The T-cells were activated as described in the ‘T cell kill assay’ section. The activated T-cells were then cultured with or without 2.5 mM Malate for 24 hours. The CD8+ activated T cells from the two treatment groups were harvested, followed by blocking FCγ receptors as previously described. The nuclei were then stained with 1 µl/ml of Ghost Dye™ UV 450 (Cytek, Catalog no. 13-0868) in FACS buffer for 30 minutes at 4°C protected from light. After washing the cells 1-2 times with FACS buffer, they were incubated in Fixation/Permeabilization solution (BD Cytofix/Cytoperm™ Fixation/Permeabilization Kit, Catalog no. 554714) for 20 minutes at 4°C. The cells were then washed twice with 1 ml of 1X BD Perm/Wash™ buffer. Subsequently, they were centrifuged at 500 x g for 5 minutes. This was followed by incubating the cells in 50 µl of Perm/Wash™ buffer (BD Cytofix/Cytoperm™ Fixation/Permeabilization Kit, Catalog no. 554714) containing a 1:200 dilution of PE anti-mouse TNF-α and Alexa Fluor® 700 anti-mouse IFN-γ (Biolegend, Catalog no. 506305 and 505823 respectively) for 30 minutes at 4°C protected from light. Wash the cells twice with 1ml of 1X BD Perm/Wash™ buffer and then resuspend the cells in FACS buffer prior to flow cytometric analysis.

### Metabolomics of CD8+ T cells and conditioned media

The CD8+ T cells were isolated from spleen of 3 female WT C57Bl/6J mice and activated as described above in the ‘T cell kill assay’ section. Post the 48hours of activation, the CD8+ T cells were cultured for 24 hours under 2 independent conditions, vehicle and 2.5mM Malate. The CD8+ T cells isolated from 3 independent animals were cultured in 2 wells providing 2 technical replicates for each of the 3 biological replicates. Each well was seeded with 5 X 10^5^ cells. After 24h treatment, whole well content (Cells + Media) was centrifuged at 500-600 X g for 10 minutes. 1ml of the supernatant (Conditioned media) was collected and flash froze in Liquid N_2_, followed by storage at −80°C. The whole cell pellet was rinsed gently with 1ml of 150mM ammonium acetate and the incubated in 700µl of 100% ice-cold methanol for 10-15 mins at 4°C. 300 µl of ice-cold HPLC-grade water was then added and the tubes were flash froze in Liquid N_2._ Later, the cell samples were stored at −80°C till the time samples were processed. At the time of assay, 30 ul of media sample was pipetted to a 1.5ml Eppendorf tube. After adding 120ul of methanol, the samples were vortexed and centrifuged at 20000 rcf for 10 minutes. The supernatant was pipetted to a glass vial and dried under a gentle nitrogen flow. The cell extract samples were dried under a gentle nitrogen flow. Dried samples (medium and cell pellets) were reconstituted in 50 ul of80% methanol and added 50 µL of 105 mM 1-Ethyl-3-(3-dimethylaminopropyl) carbodiimide (EDC) (methanol solution), 50 µL of 175 mM 3-NPH (75 % methanol solution), and 50 µL of 6% pyridine in methanol. After vertexing, the samples were put in 4 ^0^C for 30 minutes. The samples were dried after derivatization and reconstituted in 50% methanol pending LC/MS analysis. The samples were analyzed with UPLC-TQ absolute system (Waters). Metabolites separation was performed on a BEH C18 column (100mm in length and 1mm in ID, Waters). Metabolites were detected in Multiple Reaction Monitoring (MRM) mode. The data were collected in MassLynx and analyzed in TargetLynx.

### Imaging mass cytometry (Mass CyTOF)

31 TNBC FFPE tumors, each with a 1 mm core, were analyzed from the Roswell Park TNBC TMA (clinical data summarized in Supplementary Table 1). Regions of Interest (ROIs) were chosen by Dr. Gutierrez. In this cohort, ROIs measured 600 μm by 600 μm, covering the entire 1 mm core. IMC analysis was performed by the Houston Methodist Research Institute’s Immune Monitoring Core. Sample preparation involved staining tissues with metal-tagged antibodies verified by a pathologist, optimized for CyTOF® imaging. A total of 32 metal-tagged antibodies were used to examine the samples (see Supplementary Table 2), enabling detailed analysis of immune, stromal, and tumor cell heterogeneity, as well as various cell subsets and functional phenotypes within the TME. All antibodies were prepared following the manufacturer’s protocols from Standard BioTools, measured for absorbance, and stored in Candor PBS Antibody Stabilization solution at 4 °C. FFPE tissue sections underwent baking, dewaxing in xylene, rehydration with graded alcohols, and heat-induced epitope retrieval using an EZ-Retriever System at 95 °C with a Tris-Tween20 buffer at pH 9 for 20 min. After blocking with 3% BSA in TBS, sections were incubated overnight with a master antibody mix, washed, and stained for nuclei with Cell-ID Intercalator (Standard BioTools). Finally, sections were washed, air-dried, and stored for ablation. Data acquisition was performed through ablation using the Hyperion system (Standard BioTools). The data from Roswell Park TNBC cohort is detailed in our previous publication ^59^.

### Image acquisition by Hyperion and data analysis

Image acquisition was carried out in collaboration with the Flow Cytometry core at Methodist Hospital Research Institute. Tissue analysis involved using a Helios time-of-flight mass cytometer connected to the Hyperion Imaging System (Fluidigm). Before acquisition, the imaging system was auto-tuned with a 3-element tuning slide (Fluidigm) following the manufacturer’s instructions. ROIs for imaging were chosen. After flushing the ablation chamber with Helium, tissue sections were ablated spot-by-spot using a UV laser at 200 Hz and 1μm resolution. The data were saved in Fluidigm’s MCD format and exported as 16-bit OME TIFF files for further analysis.

### Processing of the Imaging Mass Cytometry data

The IMC data were stored in MCD files and initially processed with imctools to generate multichannel TIFF images. Cells were then segmented using Steinbock, with DeepCell chosen as the segmentation algorithm, based on Ir191 and Ir193 channels for nuclei and surface markers for cell membranes. Using these cell masks, we created a cell-by-protein matrix that summarized pixel intensities for each cell and marker. The cell position was defined by the centroid of each mask. Data were log-transformed and z-scored, first across all cells and then across all proteins. We performed K-means clustering with 20 clusters, setting nstart=100,000 for robust results. Differential proteins between clusters were identified through one-vs-one comparisons, and clusters were named after their differentially expressed markers.

### Spatial cell-cell interaction analysis from IMC data

We initially created a Delauney spatial graph connecting individual cells within each ROI. Subsequently, we conducted a spatial proximity analysis using the Giotto pipeline with the 20 cell-type cluster labels. The Cell Proximity Enrichment feature in Giotto, involving 1000 random simulations, detected cell-type interaction enrichment or depletion relative to randomized labels, where cell types are shuffled. This process produced a z-score for each cell-type pair, reflecting either enrichment (positive z-score) or depletion (negative z-score) of interactions for each ROI. We performed group comparisons between AADAT knockdown and wildtype conditions using a linear mixed model that accounts for variability among animals and the repeated measures from multiple ROIs per animal.

### Sample preparation for IMS data

TMA sections were dewaxed with 2 times 3 minutes with fresh xylene and allowed to air dry. Fiducial marks were added at the corners of the slide, and an optical image was acquired at 4800 dpi using an Epson Perfection V600 flatbed scanner. The slide was coated with 10 mg/mL 1,5-diaminonaphthalene in 50% acetonitrile using an HTX M5 Robotic Reagent Sprayer over 5 passes with the following parameters: a flow rate of 0.010 mL/min, a track speed of 1200 mm/min, a track spacing of 3 mm, a nozzle height of 40 mm, a nozzle temperature of 60°C, and a crisscross track pattern.

### IMS data collection

Metabolite image data was collected in negative ion mode at 50 µm resolution on a Bruker timsTOF fleX QTOF mass spectrometer using FlexImaging 5.1. Instrument parameters were optimized for metabolites as follows: an *m/z* range of 50-600, TIMS OFF, a Funnel 1 RF of 75.0 Vpp, a Funnel 2 RF of 100.0 Vpp, a Multipole RF of 150.0 Vpp, a Collision Energy of 10.0 eV, a Collision RF of 500.0 Vpp, a Quadrupole Ion Energy of 5.0 eV, a Transfer Time of 50.0 µs, and a Pre Pulse Storage of 5.0 µs. A total of 300 laser shots were summed per pixel.

### IMS data analysis

Image data were visualized using SCiLS Lab 2025b and Root Mean Square normalized before further analysis. Peaks were manually selected from the average spectrum with an integration window of ±15 ppm. Putative metabolite IDs were assigned using the MetaboScape 2025 plugin for SCiLS, allowing a 10-ppm mass tolerance. Pixel data for the entire TMA were exported to a .csv file using the SCiLS Lab API for R to allow for integration with IMC data.

### Association between T cell clusters and malate

The relationship between malate levels and various cell-type proportions, focusing on functional T cells and exhausted T cells, was examined using binned data. First, all spatial IMS-IMC spots were sorted by malate concentration. We then divided these spots into percentile bins (every 5th percentile, totaling 20 bins). For each bin, we calculated the average malate level per spot from IMS data and the average cell-type proportions per spot from IMC data at the same position. The mean malate level was plotted against the mean cell-type abundance across all bins. A trend line was added to show the association direction, and Pearson correlation was used to measure its strength.

### Spatial distribution of malate and the cell types

Spots were ranked based on their malate levels, and the top and bottom 10% were selected and subsetted for each ROI. For each ROI, the malate concentration at the spot level was plotted against the corresponding cell type proportion. Spots with malate levels above zero are shown in yellow, with the color intensity proportional to the concentration. Similarly, the cell type concentration for spots with malate present is shown in red, with the intensity indicating the abundance.

### Survival analysis and Cox proportional hazard analysis

Survival analysis was conducted with the survival R package and visualized using the autoplot function from ggplot2. Patients were grouped based on their average malate levels. Kaplan-Meier curves were then generated using R’s survival library. For Cox Proportional Hazards analysis, we linked spatial malate levels to patients’ 10-year survival status in the Roswell Park TNBC cohort. Each cohort included patient attributes such as age, stage, and grade. We developed multivariate models to examine how spatial malate levels relate to survival, while controlling for covariates such as age and stage. The model was specified as “Surv(time, status) ∼ malateHighSpots + age + stage,” where malateHighSpots is the count of high-malate spots per patient derived from spatial IMS data. Hazard ratios and P-values were then calculated.

### Statistical analysis

Data analysis was performed using GraphPad Prism (v10.6.1) or R. Sample sizes and the specific statistical tests are detailed in the figures or their legends. A p-value under 0.05 was deemed statistically significant. Unless specified otherwise, data are shown as mean ± SEM. Both in vitro and in vivo datasets were tested for normality using the Shapiro-Wilk test, and parametric tests were only applied to data passing this test. All in vitro experiments were conducted in triplicate. Biological and technical replicates are clearly identified in the figure legends for all experiments.

## Ethics approval

The Institutional Animal Care and Use Committee of Baylor College of Medicine approved all animal studies. All the human studies were performed under IRB (Institutional Review Board) protocols 020-393 and 130559 (Baylor Scott and White Hospital), H-28445 (Baylor College of Medicine), and Roswell Park Comprehensive Cancer Center (RPCCC). RPCCC samples were used with patient consent. The approved BSW IRB protocol 020-393 waived the requirement of authorization based on 45 CFR 164.521(i)(2)(ii) and determined informed consent is not required as allowed under 45 CFR 46.116 (g).

## Availability of data and materials

All data supporting this study are available upon request. The metabolomics data have been submitted to the NIH Metabolomics Workbench under Study ID ST004340. The data can be accessed directly via its Project DOI: http://dx.doi.org/10.21228/M8XR8W. The transcriptomics data for T cells is available in GEO under the accession GSE317623.

## Consent for publication

All authors give consent for the publication of the manuscript.

## Consent for publication

All authors give consent for the publication of the manuscript.

## Author contributions

MC, FG, XHZ, and AS designed the study. MC, FG, SS, UR, SMT, YQ, LEEF, HV, MKM, JHP, WZ, LT, LY, YG, BWS, SYJ, BK, VP, NC, EHS, BAK, IJK, and NP performed experiments. DV, BP, and QZ analyzed data. NM, DJ, JRA, AR, CG, ARO, CM, GMD, and CA performed histopathology or provided reagents for the study. MC, FG, and AS drafted the manuscript. MC, FG, BAK, IJK, NP, MKM, AE, AAD, QZ, XHFZ, and AS edited the manuscript.

Co-first author order placed in alphabetical order of their last name.

## Acknowledgements

We thank the Human Tissue Acquisition and Pathology and Cytometry and Cell Sorting Cores at Baylor College of Medicine for their contributions to the study.

