## Supplemental tables and figures for "AADAT-Driven Metabolic Control of Malate and CoQ_10_ Shapes Immune Evasion in Triple-Negative Breast Cancer"

### Slide 1
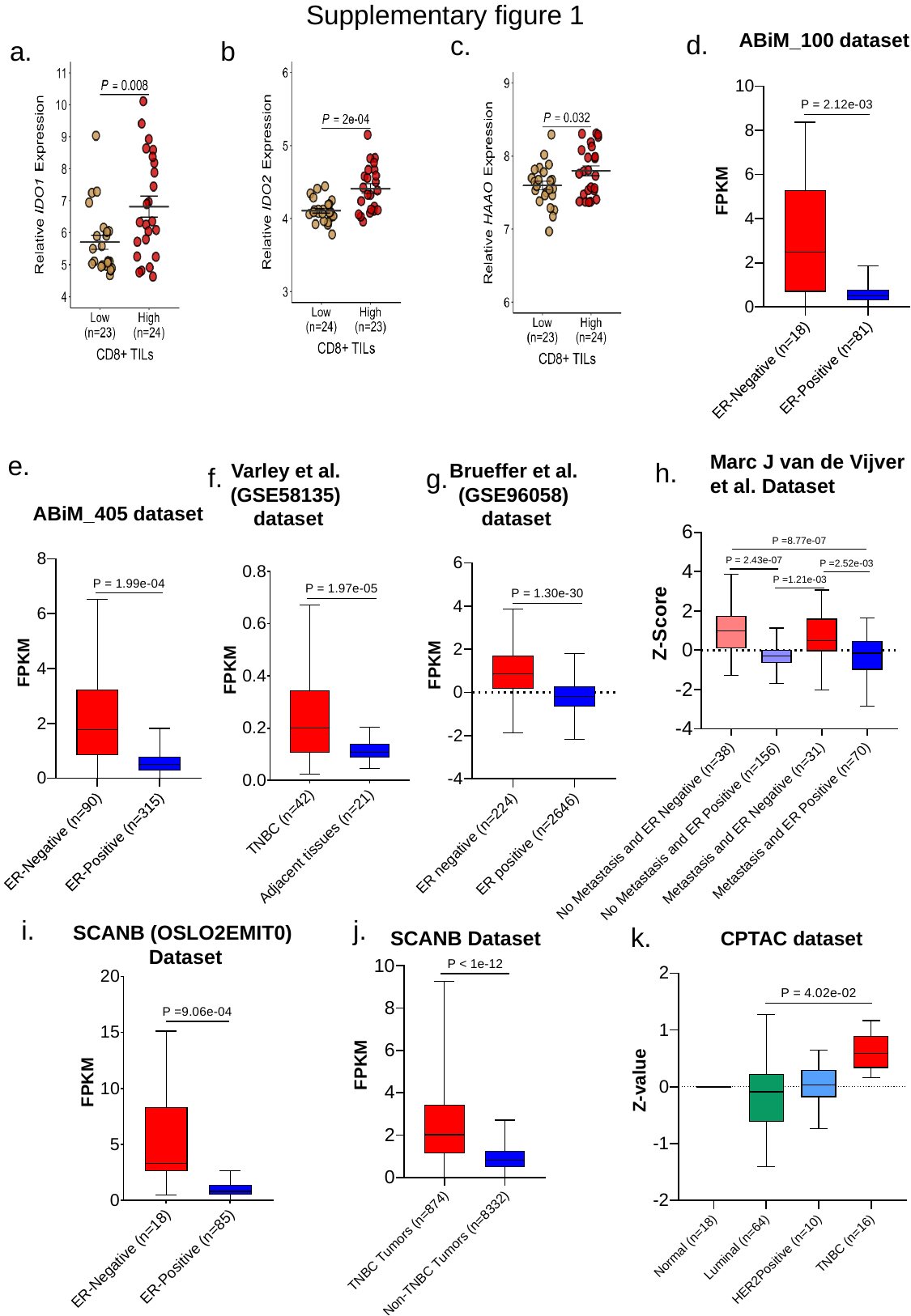

Supplementary figure 1
d.
ABiM_100 dataset
c.
a.
b.
e.
Marc J van de Vijver
et al. Dataset
h.
Varley et al.
(GSE58135)
dataset
Brueffer et al.
(GSE96058)
dataset
f.
g.
ABiM_405 dataset
i.
j.
SCANB (OSLO2EMIT0)
Dataset
k.
SCANB Dataset
CPTAC dataset

### Slide 2
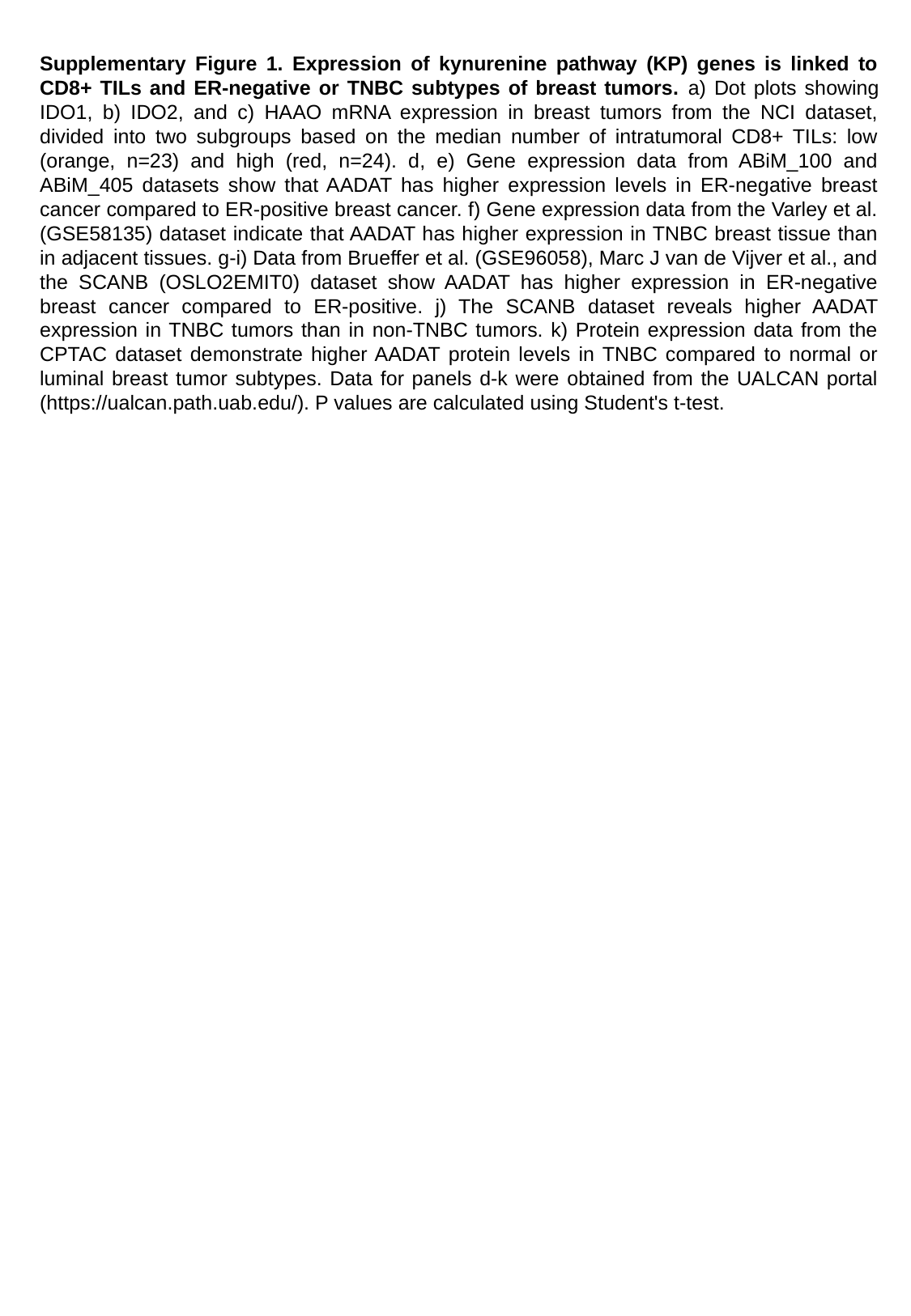

Supplementary Figure 1. Expression of kynurenine pathway (KP) genes is linked to CD8+ TILs and ER-negative or TNBC subtypes of breast tumors. a) Dot plots showing IDO1, b) IDO2, and c) HAAO mRNA expression in breast tumors from the NCI dataset, divided into two subgroups based on the median number of intratumoral CD8+ TILs: low (orange, n=23) and high (red, n=24). d, e) Gene expression data from ABiM_100 and ABiM_405 datasets show that AADAT has higher expression levels in ER-negative breast cancer compared to ER-positive breast cancer. f) Gene expression data from the Varley et al. (GSE58135) dataset indicate that AADAT has higher expression in TNBC breast tissue than in adjacent tissues. g-i) Data from Brueffer et al. (GSE96058), Marc J van de Vijver et al., and the SCANB (OSLO2EMIT0) dataset show AADAT has higher expression in ER-negative breast cancer compared to ER-positive. j) The SCANB dataset reveals higher AADAT expression in TNBC tumors than in non-TNBC tumors. k) Protein expression data from the CPTAC dataset demonstrate higher AADAT protein levels in TNBC compared to normal or luminal breast tumor subtypes. Data for panels d-k were obtained from the UALCAN portal (https://ualcan.path.uab.edu/). P values are calculated using Student's t-test.

### Slide 3
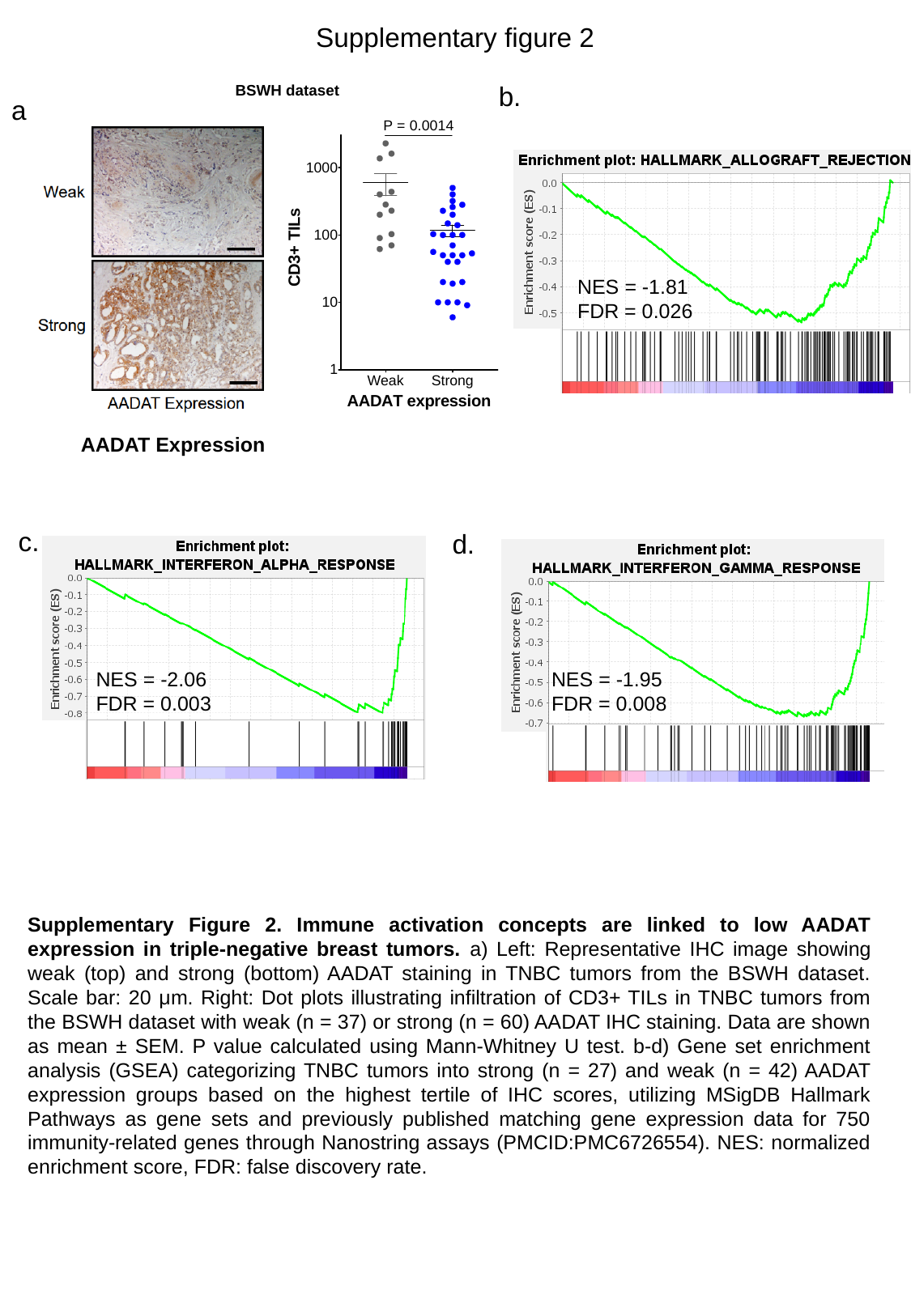

Supplementary figure 2
b.
BSWH dataset
a.
NES = -1.81
FDR = 0.026
AADAT Expression
c.
d.
NES = -2.06
FDR = 0.003
NES = -1.95
FDR = 0.008
Supplementary Figure 2. Immune activation concepts are linked to low AADAT expression in triple-negative breast tumors. a) Left: Representative IHC image showing weak (top) and strong (bottom) AADAT staining in TNBC tumors from the BSWH dataset. Scale bar: 20 μm. Right: Dot plots illustrating infiltration of CD3+ TILs in TNBC tumors from the BSWH dataset with weak (n = 37) or strong (n = 60) AADAT IHC staining. Data are shown as mean ± SEM. P value calculated using Mann-Whitney U test. b-d) Gene set enrichment analysis (GSEA) categorizing TNBC tumors into strong (n = 27) and weak (n = 42) AADAT expression groups based on the highest tertile of IHC scores, utilizing MSigDB Hallmark Pathways as gene sets and previously published matching gene expression data for 750 immunity-related genes through Nanostring assays (PMCID:PMC6726554). NES: normalized enrichment score, FDR: false discovery rate.

### Slide 4
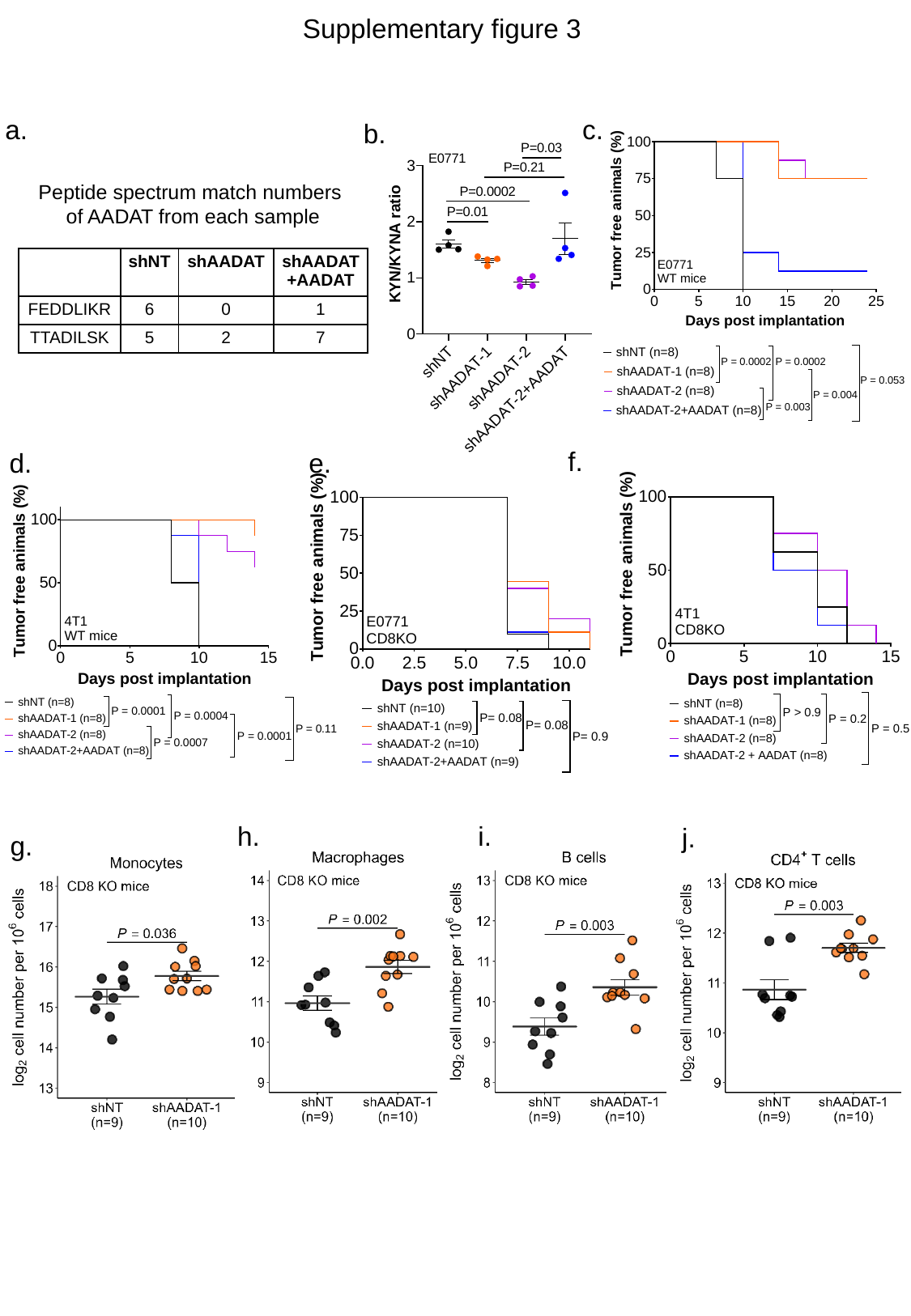

Supplementary figure 3
a.
c.
b.
Peptide spectrum match numbers
of AADAT from each sample
| | shNT | shAADAT | shAADAT+AADAT |
| --- | --- | --- | --- |
| FEDDLIKR | 6 | 0 | 1 |
| TTADILSK | 5 | 2 | 7 |
f.
d.
e.
h.
i.
j.
g.

### Slide 5
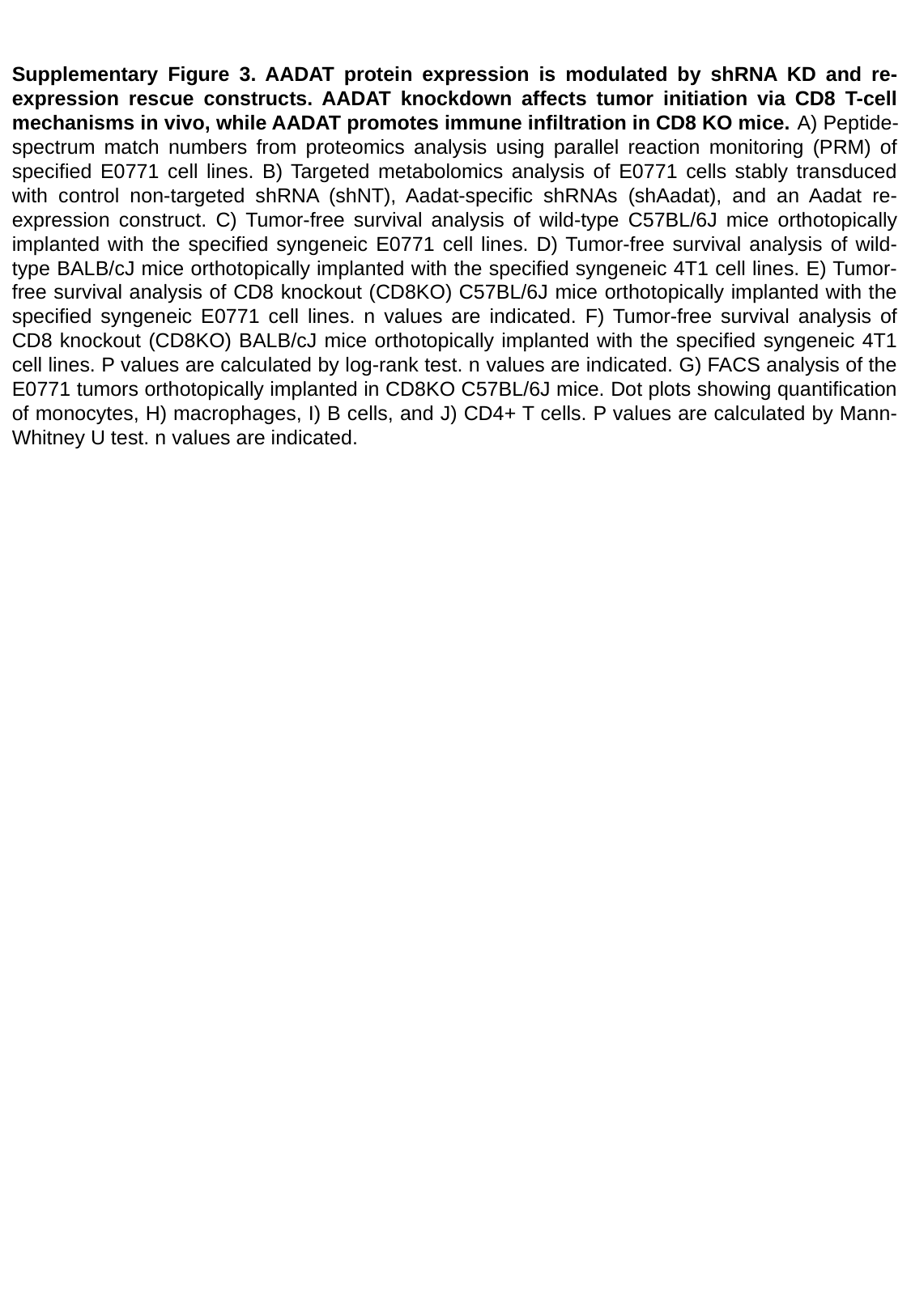

Supplementary Figure 3. AADAT protein expression is modulated by shRNA KD and re-expression rescue constructs. AADAT knockdown affects tumor initiation via CD8 T-cell mechanisms in vivo, while AADAT promotes immune infiltration in CD8 KO mice. A) Peptide-spectrum match numbers from proteomics analysis using parallel reaction monitoring (PRM) of specified E0771 cell lines. B) Targeted metabolomics analysis of E0771 cells stably transduced with control non-targeted shRNA (shNT), Aadat-specific shRNAs (shAadat), and an Aadat re-expression construct. C) Tumor-free survival analysis of wild-type C57BL/6J mice orthotopically implanted with the specified syngeneic E0771 cell lines. D) Tumor-free survival analysis of wild-type BALB/cJ mice orthotopically implanted with the specified syngeneic 4T1 cell lines. E) Tumor-free survival analysis of CD8 knockout (CD8KO) C57BL/6J mice orthotopically implanted with the specified syngeneic E0771 cell lines. n values are indicated. F) Tumor-free survival analysis of CD8 knockout (CD8KO) BALB/cJ mice orthotopically implanted with the specified syngeneic 4T1 cell lines. P values are calculated by log-rank test. n values are indicated. G) FACS analysis of the E0771 tumors orthotopically implanted in CD8KO C57BL/6J mice. Dot plots showing quantification of monocytes, H) macrophages, I) B cells, and J) CD4+ T cells. P values are calculated by Mann-Whitney U test. n values are indicated.

### Slide 6
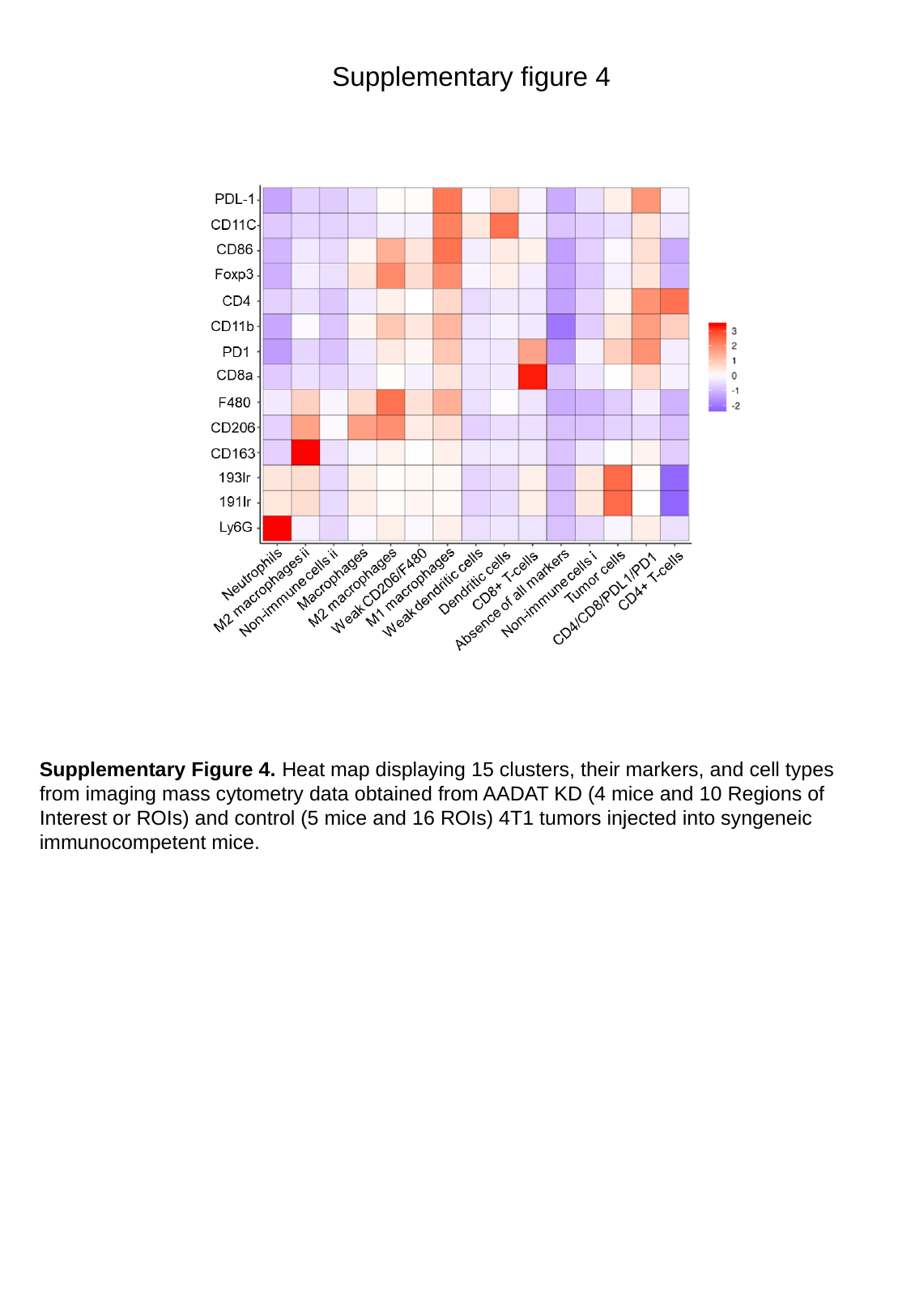

Supplementary figure 4
Supplementary Figure 4. Heat map displaying 15 clusters, their markers, and cell types from imaging mass cytometry data obtained from AADAT KD (4 mice and 10 Regions of Interest or ROIs) and control (5 mice and 16 ROIs) 4T1 tumors injected into syngeneic immunocompetent mice.

### Slide 7
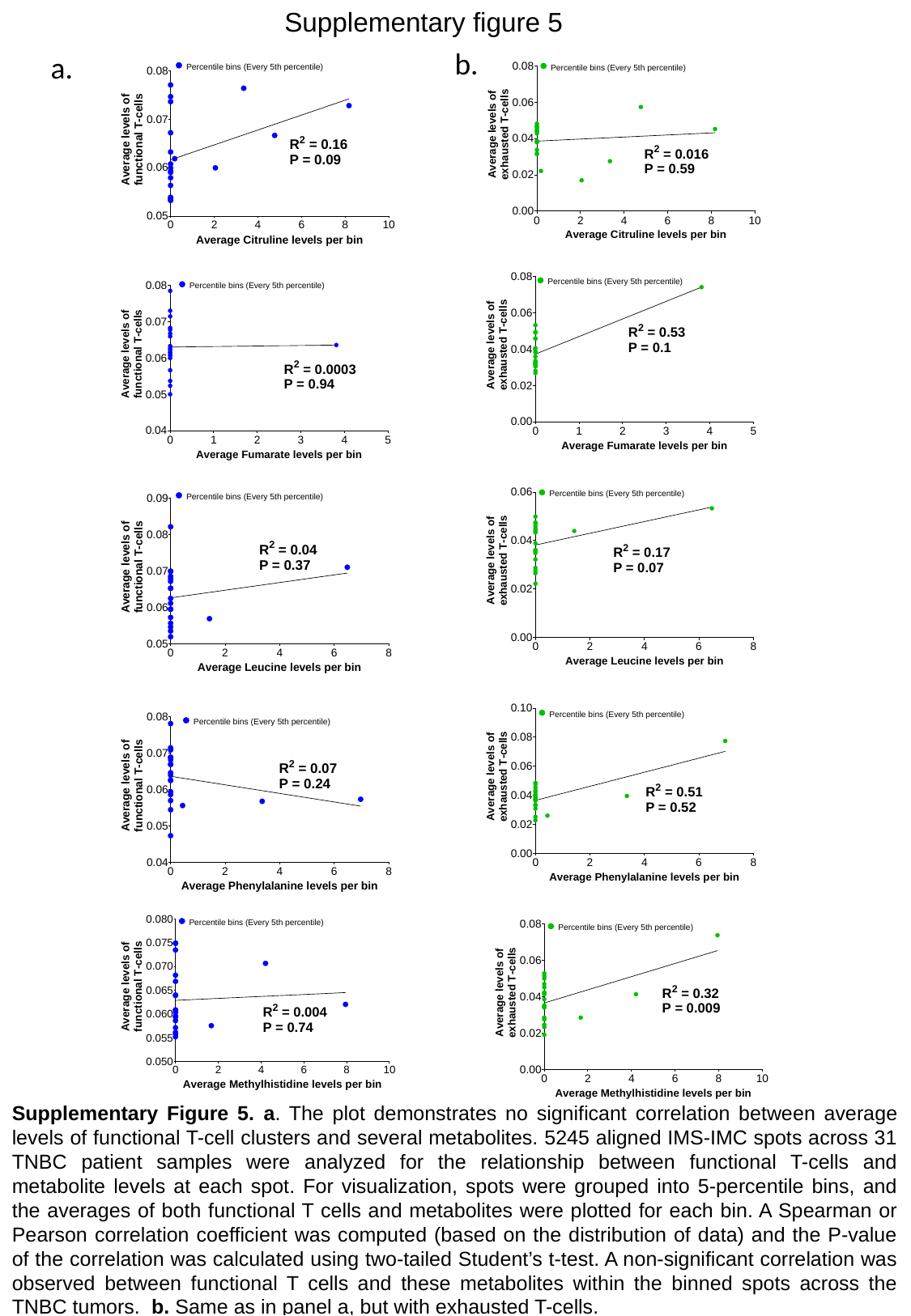

Supplementary figure 5
b.
a.
Supplementary Figure 5. a. The plot demonstrates no significant correlation between average levels of functional T-cell clusters and several metabolites. 5245 aligned IMS-IMC spots across 31 TNBC patient samples were analyzed for the relationship between functional T-cells and metabolite levels at each spot. For visualization, spots were grouped into 5-percentile bins, and the averages of both functional T cells and metabolites were plotted for each bin. A Spearman or Pearson correlation coefficient was computed (based on the distribution of data) and the P-value of the correlation was calculated using two-tailed Student’s t-test. A non-significant correlation was observed between functional T cells and these metabolites within the binned spots across the TNBC tumors. b. Same as in panel a, but with exhausted T-cells.

### Slide 8
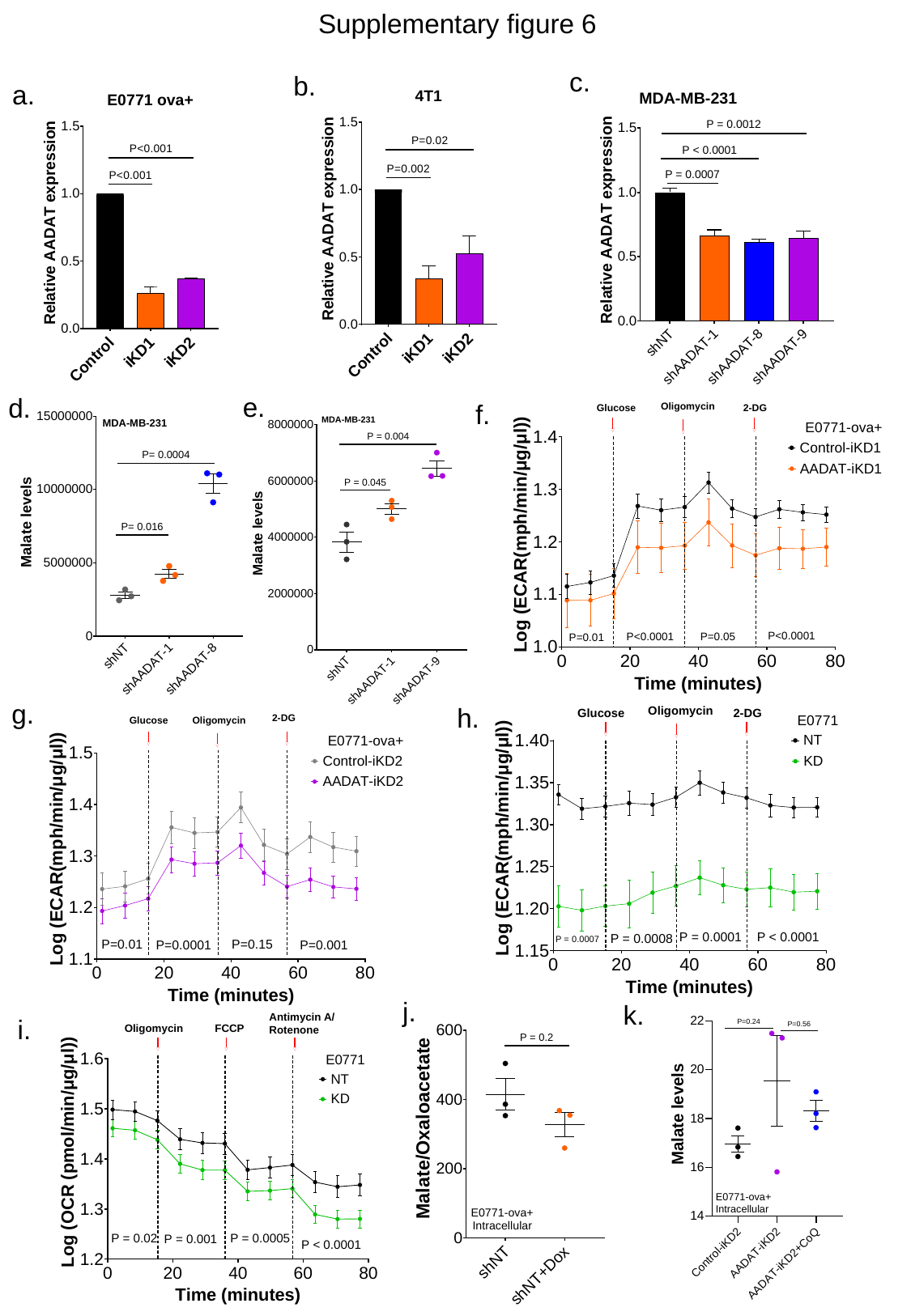

Supplementary figure 6
c.
b.
a.
e.
d.
f.
g.
h.
j.
k.
i.

### Slide 9
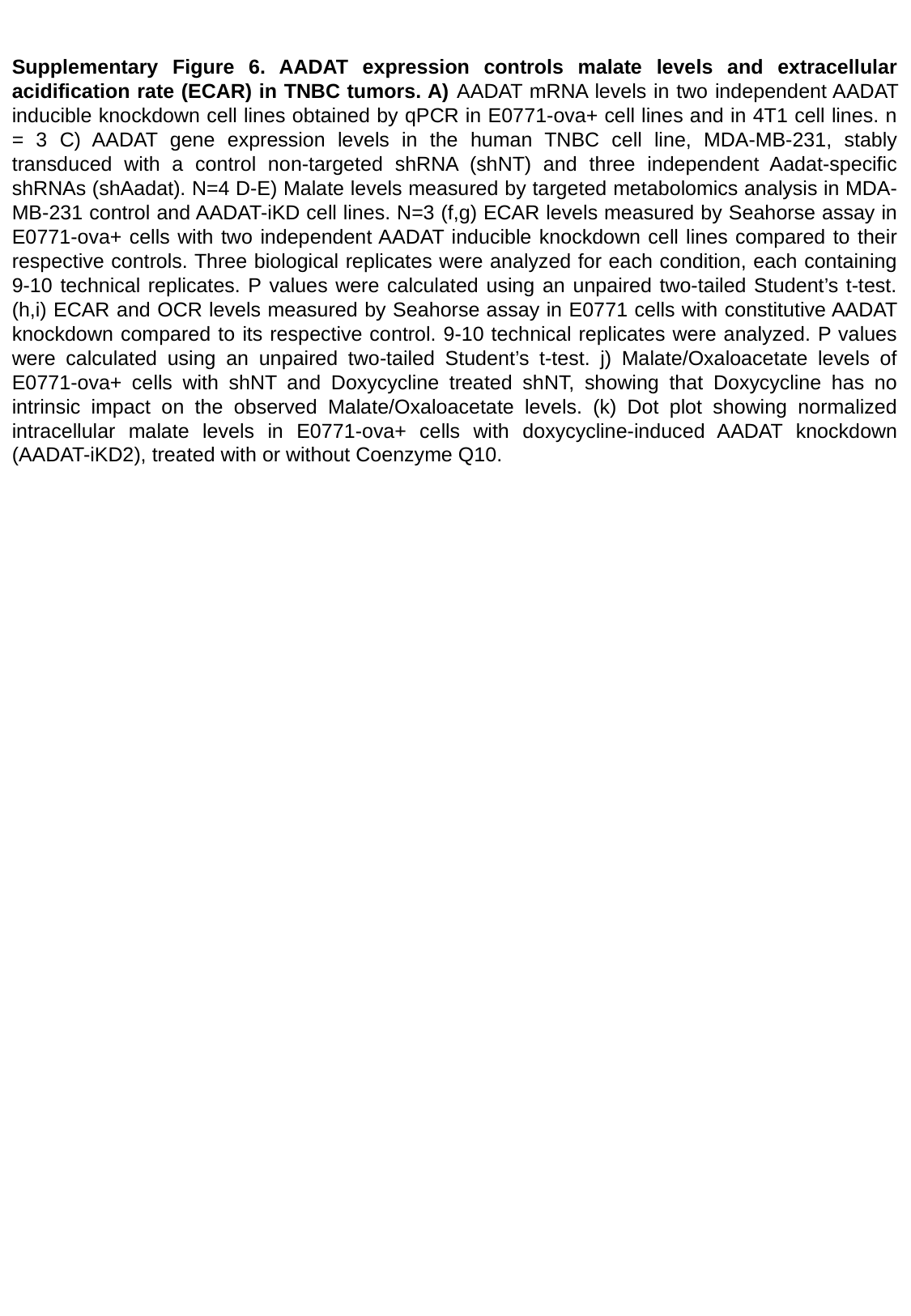

Supplementary Figure 6. AADAT expression controls malate levels and extracellular acidification rate (ECAR) in TNBC tumors. A) AADAT mRNA levels in two independent AADAT inducible knockdown cell lines obtained by qPCR in E0771-ova+ cell lines and in 4T1 cell lines. n = 3 C) AADAT gene expression levels in the human TNBC cell line, MDA-MB-231, stably transduced with a control non-targeted shRNA (shNT) and three independent Aadat-specific shRNAs (shAadat). N=4 D-E) Malate levels measured by targeted metabolomics analysis in MDA-MB-231 control and AADAT-iKD cell lines. N=3 (f,g) ECAR levels measured by Seahorse assay in E0771-ova+ cells with two independent AADAT inducible knockdown cell lines compared to their respective controls. Three biological replicates were analyzed for each condition, each containing 9-10 technical replicates. P values were calculated using an unpaired two-tailed Student’s t-test. (h,i) ECAR and OCR levels measured by Seahorse assay in E0771 cells with constitutive AADAT knockdown compared to its respective control. 9-10 technical replicates were analyzed. P values were calculated using an unpaired two-tailed Student’s t-test. j) Malate/Oxaloacetate levels of E0771-ova+ cells with shNT and Doxycycline treated shNT, showing that Doxycycline has no intrinsic impact on the observed Malate/Oxaloacetate levels. (k) Dot plot showing normalized intracellular malate levels in E0771-ova+ cells with doxycycline-induced AADAT knockdown (AADAT-iKD2), treated with or without Coenzyme Q10.

### Slide 10
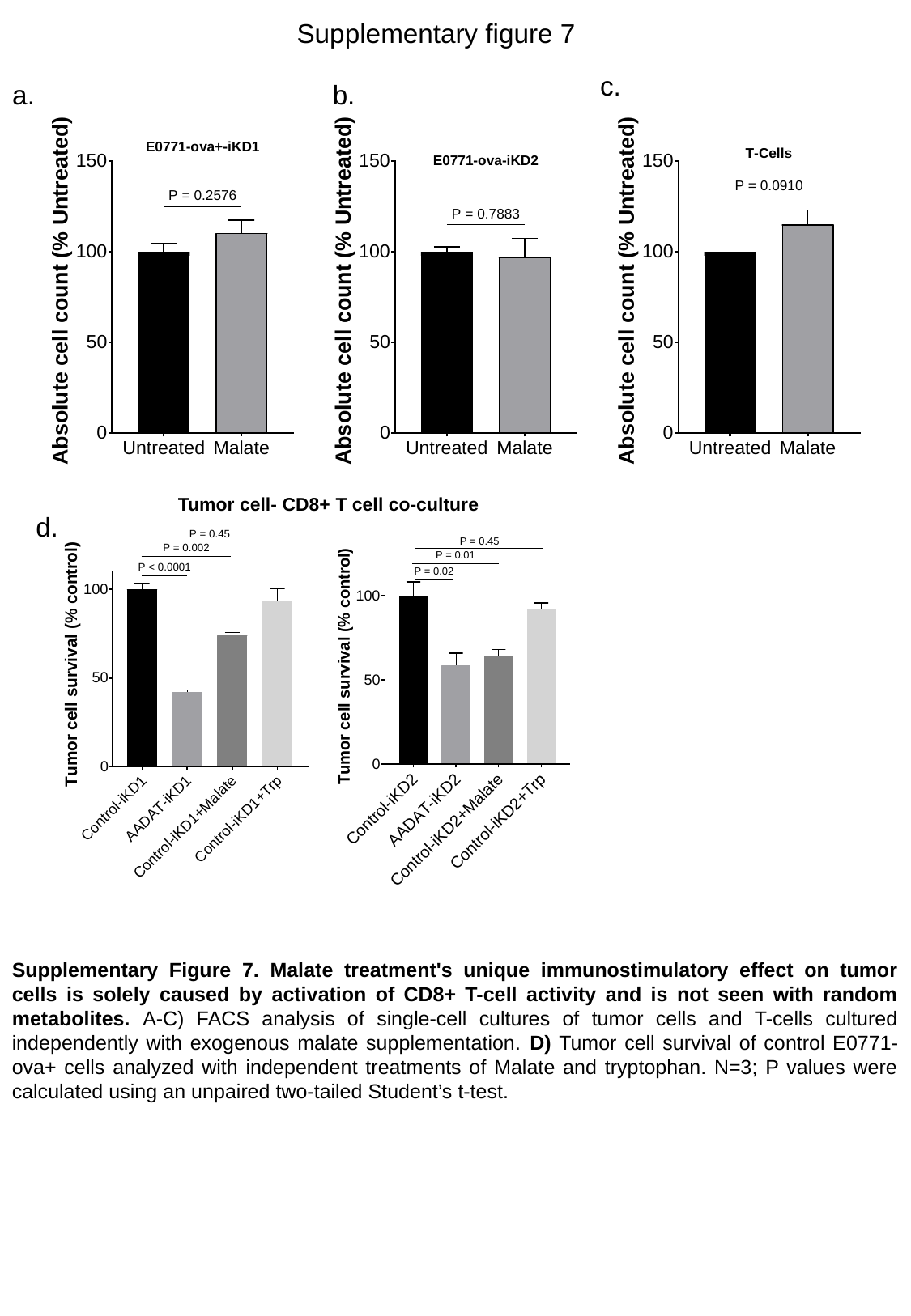

Supplementary figure 7
c.
a.
b.
Tumor cell- CD8+ T cell co-culture
d.
Supplementary Figure 7. Malate treatment's unique immunostimulatory effect on tumor cells is solely caused by activation of CD8+ T-cell activity and is not seen with random metabolites. A-C) FACS analysis of single-cell cultures of tumor cells and T-cells cultured independently with exogenous malate supplementation. D) Tumor cell survival of control E0771-ova+ cells analyzed with independent treatments of Malate and tryptophan. N=3; P values were calculated using an unpaired two-tailed Student’s t-test.

### Slide 11
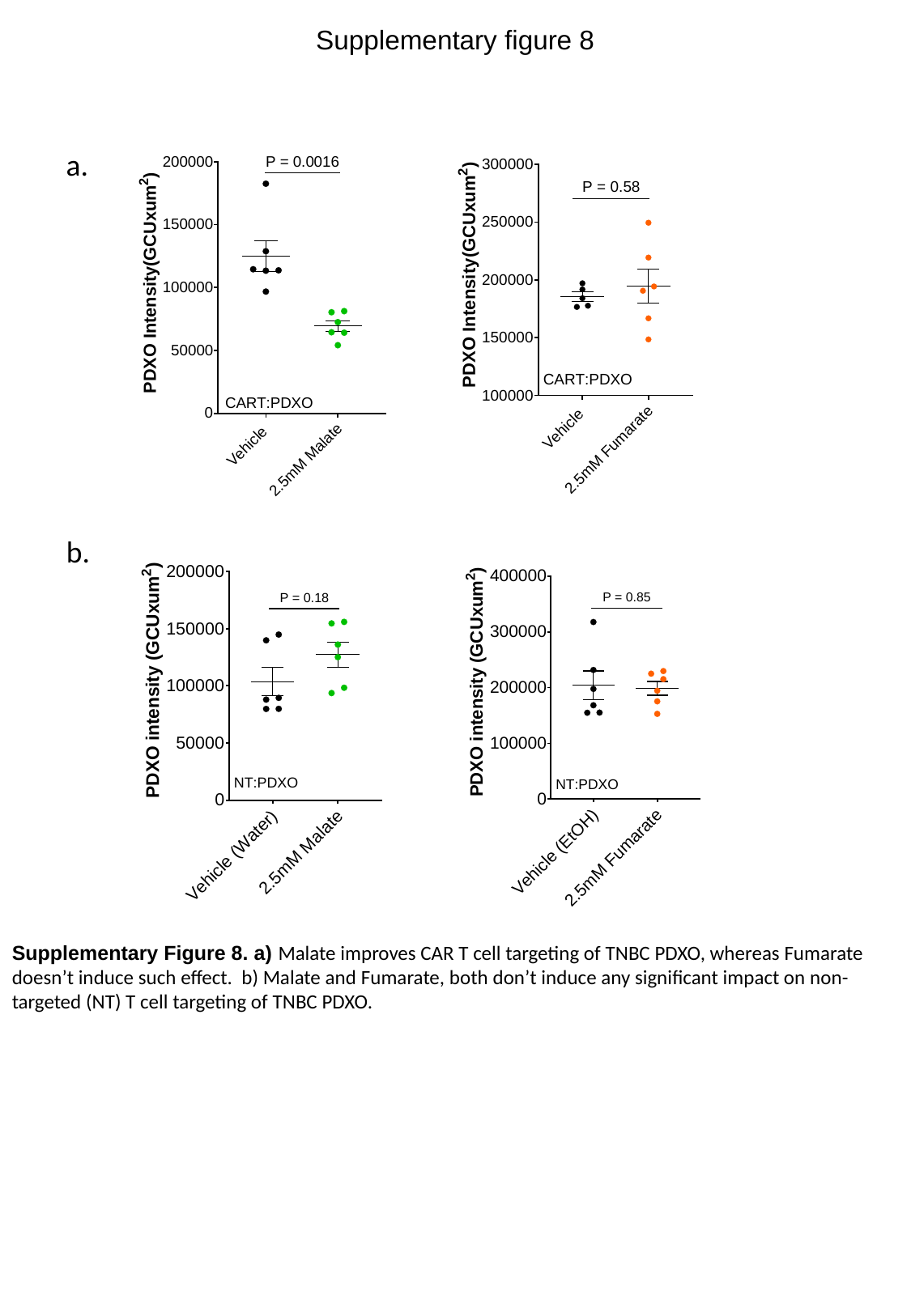

Supplementary figure 8
a.
b.
Supplementary Figure 8. a) Malate improves CAR T cell targeting of TNBC PDXO, whereas Fumarate doesn’t induce such effect. b) Malate and Fumarate, both don’t induce any significant impact on non-targeted (NT) T cell targeting of TNBC PDXO.

### Slide 12
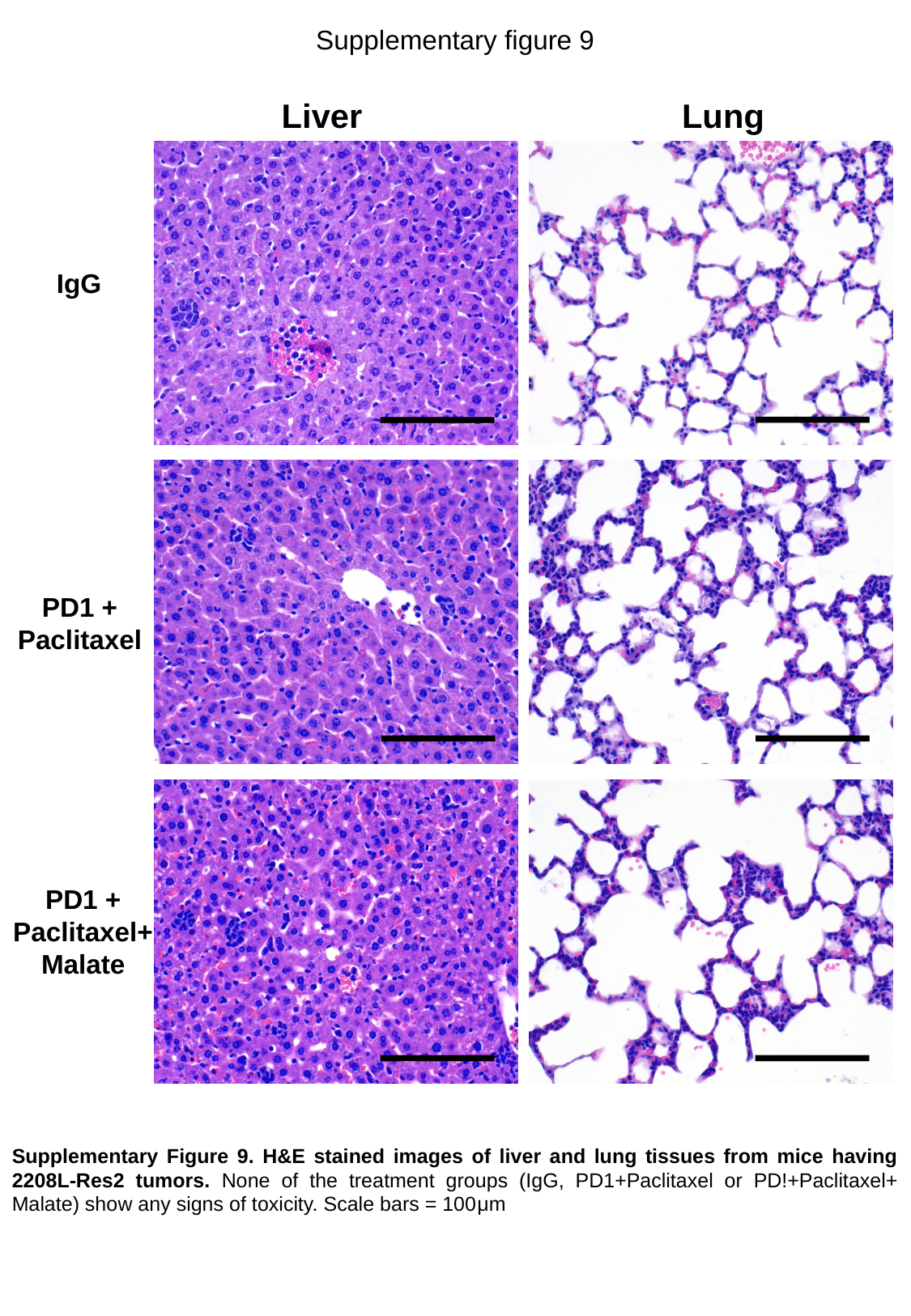

Supplementary figure 9
Liver
Lung
IgG
PD1 +
Paclitaxel
PD1 +
Paclitaxel+
Malate
Supplementary Figure 9. H&E stained images of liver and lung tissues from mice having 2208L-Res2 tumors. None of the treatment groups (IgG, PD1+Paclitaxel or PD!+Paclitaxel+ Malate) show any signs of toxicity. Scale bars = 100μm

### Slide 13
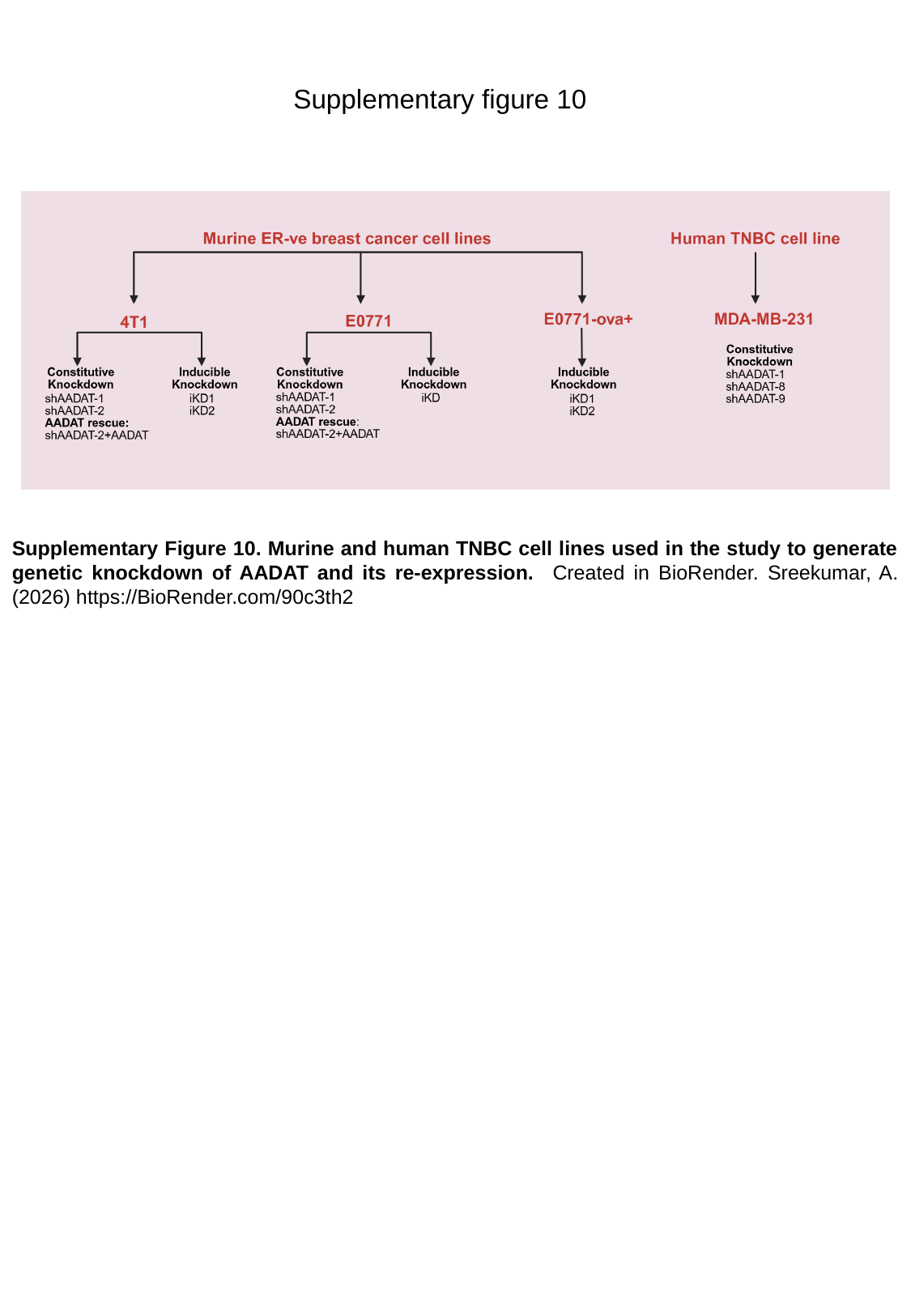

Supplementary figure 10
Supplementary Figure 10. Murine and human TNBC cell lines used in the study to generate genetic knockdown of AADAT and its re-expression. Created in BioRender. Sreekumar, A. (2026) https://BioRender.com/90c3th2

### Slide 14
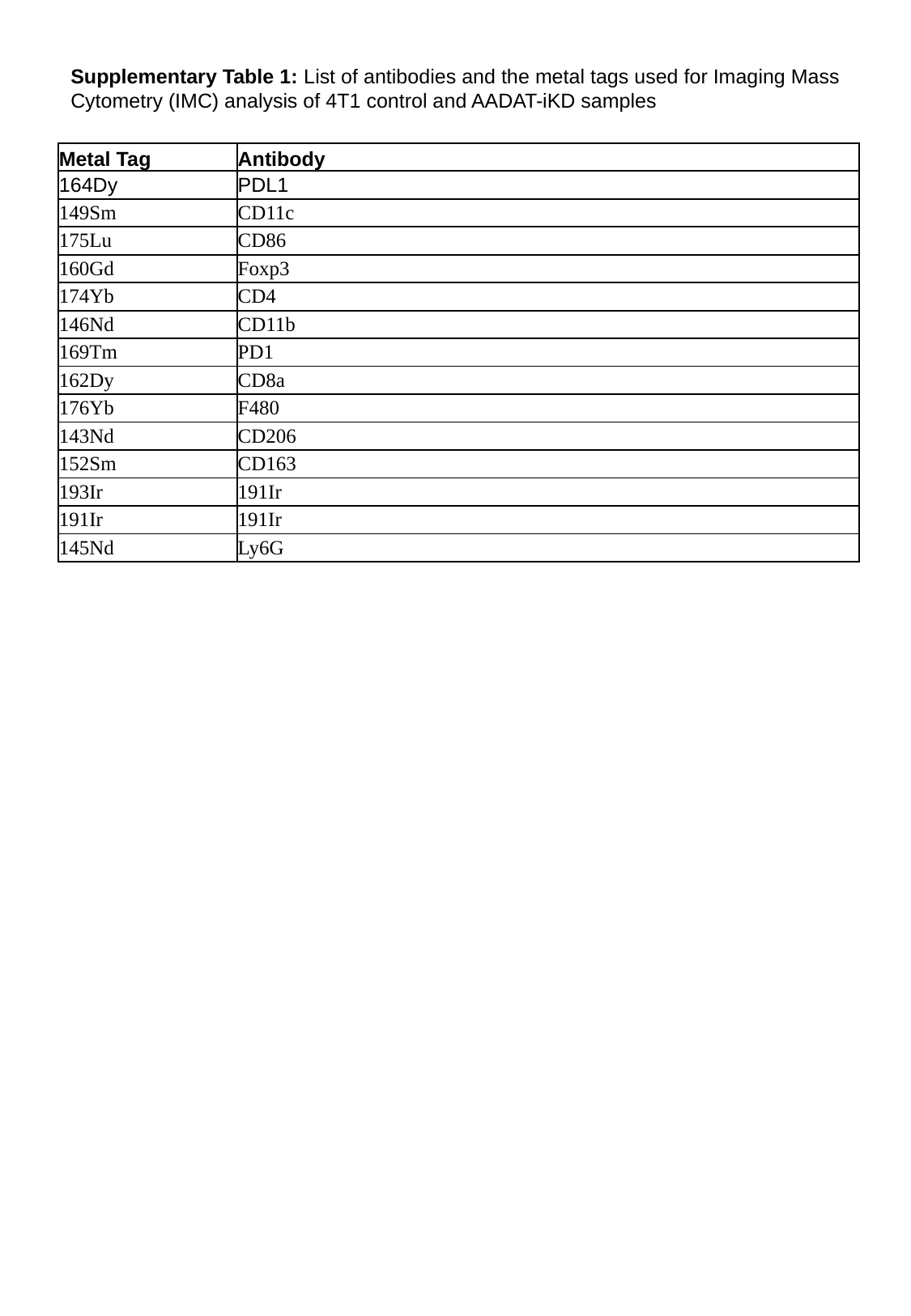

Supplementary Table 1: List of antibodies and the metal tags used for Imaging Mass Cytometry (IMC) analysis of 4T1 control and AADAT-iKD samples
| Metal Tag | Antibody |
| --- | --- |
| 164Dy | PDL1 |
| 149Sm | CD11c |
| 175Lu | CD86 |
| 160Gd | Foxp3 |
| 174Yb | CD4 |
| 146Nd | CD11b |
| 169Tm | PD1 |
| 162Dy | CD8a |
| 176Yb | F480 |
| 143Nd | CD206 |
| 152Sm | CD163 |
| 193Ir | 191Ir |
| 191Ir | 191Ir |
| 145Nd | Ly6G |

### Slide 15
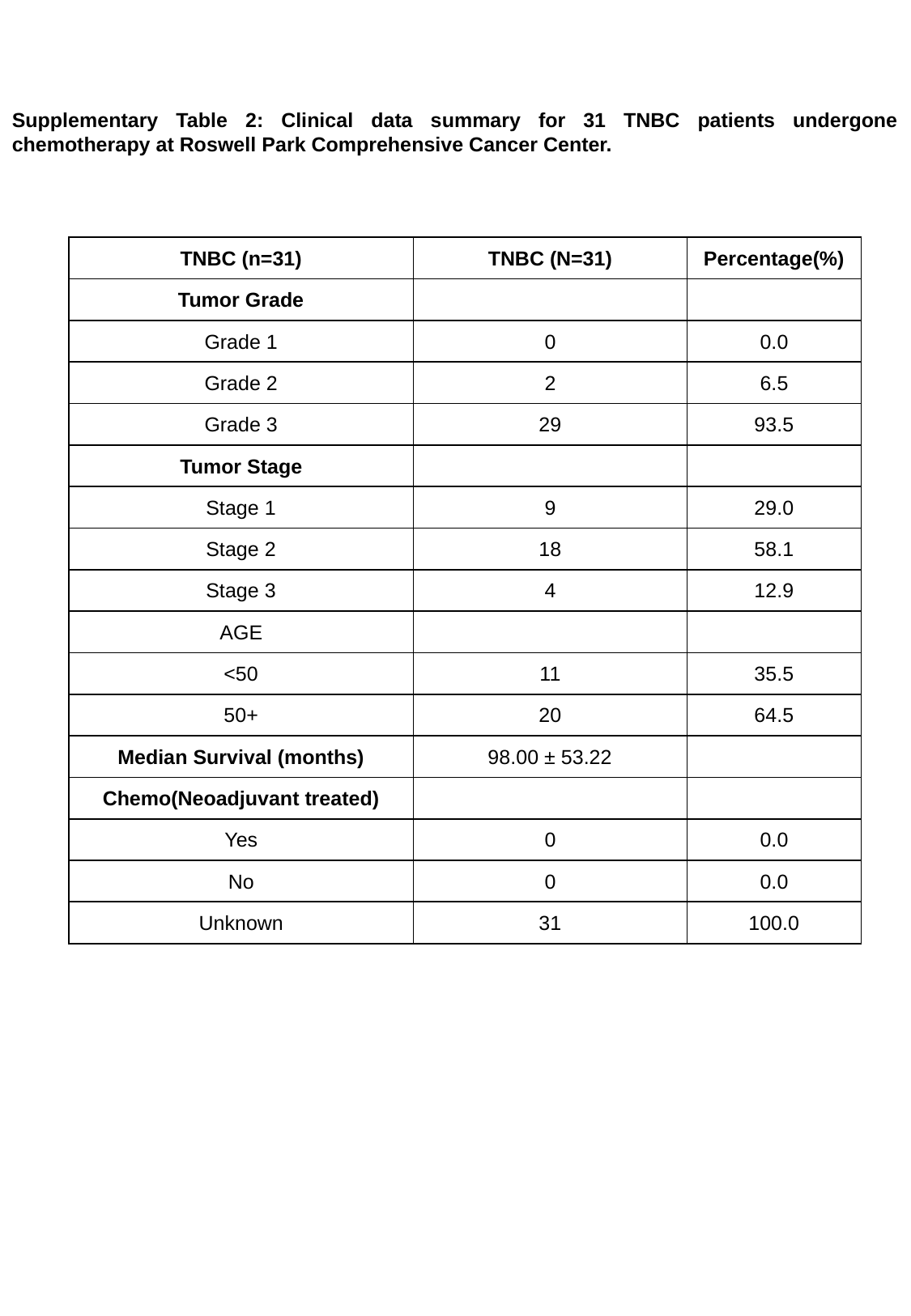

Supplementary Table 2: Clinical data summary for 31 TNBC patients undergone chemotherapy at Roswell Park Comprehensive Cancer Center.
| TNBC (n=31) | TNBC (N=31) | Percentage(%) |
| --- | --- | --- |
| Tumor Grade | | |
| Grade 1 | 0 | 0.0 |
| Grade 2 | 2 | 6.5 |
| Grade 3 | 29 | 93.5 |
| Tumor Stage | | |
| Stage 1 | 9 | 29.0 |
| Stage 2 | 18 | 58.1 |
| Stage 3 | 4 | 12.9 |
| AGE | | |
| <50 | 11 | 35.5 |
| 50+ | 20 | 64.5 |
| Median Survival (months) | 98.00 ± 53.22 | |
| Chemo(Neoadjuvant treated) | | |
| Yes | 0 | 0.0 |
| No | 0 | 0.0 |
| Unknown | 31 | 100.0 |

### Slide 16
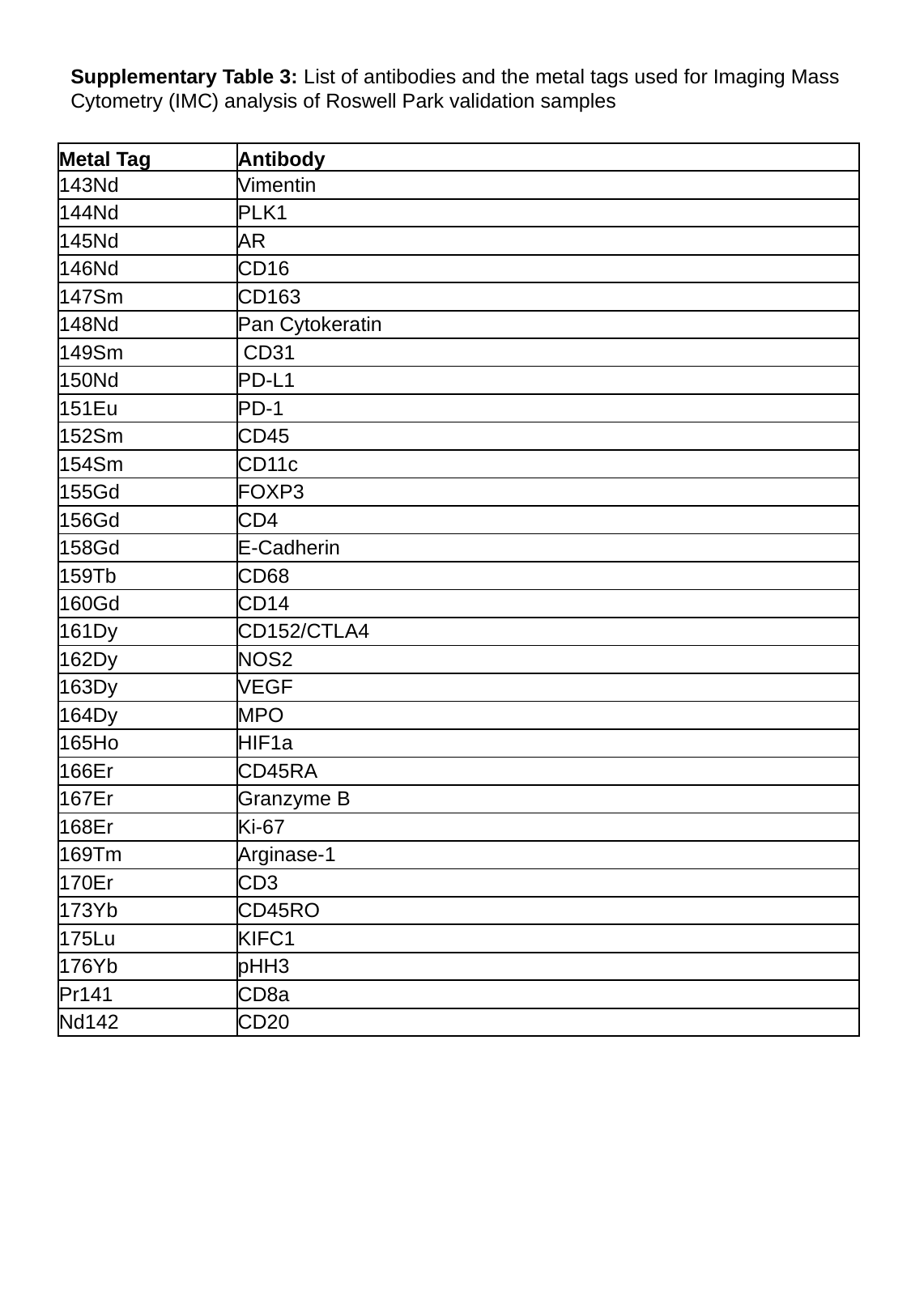

Supplementary Table 3: List of antibodies and the metal tags used for Imaging Mass Cytometry (IMC) analysis of Roswell Park validation samples
| Metal Tag | Antibody |
| --- | --- |
| 143Nd | Vimentin |
| 144Nd | PLK1 |
| 145Nd | AR |
| 146Nd | CD16 |
| 147Sm | CD163 |
| 148Nd | Pan Cytokeratin |
| 149Sm | CD31 |
| 150Nd | PD-L1 |
| 151Eu | PD-1 |
| 152Sm | CD45 |
| 154Sm | CD11c |
| 155Gd | FOXP3 |
| 156Gd | CD4 |
| 158Gd | E-Cadherin |
| 159Tb | CD68 |
| 160Gd | CD14 |
| 161Dy | CD152/CTLA4 |
| 162Dy | NOS2 |
| 163Dy | VEGF |
| 164Dy | MPO |
| 165Ho | HIF1a |
| 166Er | CD45RA |
| 167Er | Granzyme B |
| 168Er | Ki-67 |
| 169Tm | Arginase-1 |
| 170Er | CD3 |
| 173Yb | CD45RO |
| 175Lu | KIFC1 |
| 176Yb | pHH3 |
| Pr141 | CD8a |
| Nd142 | CD20 |

### Slide 17
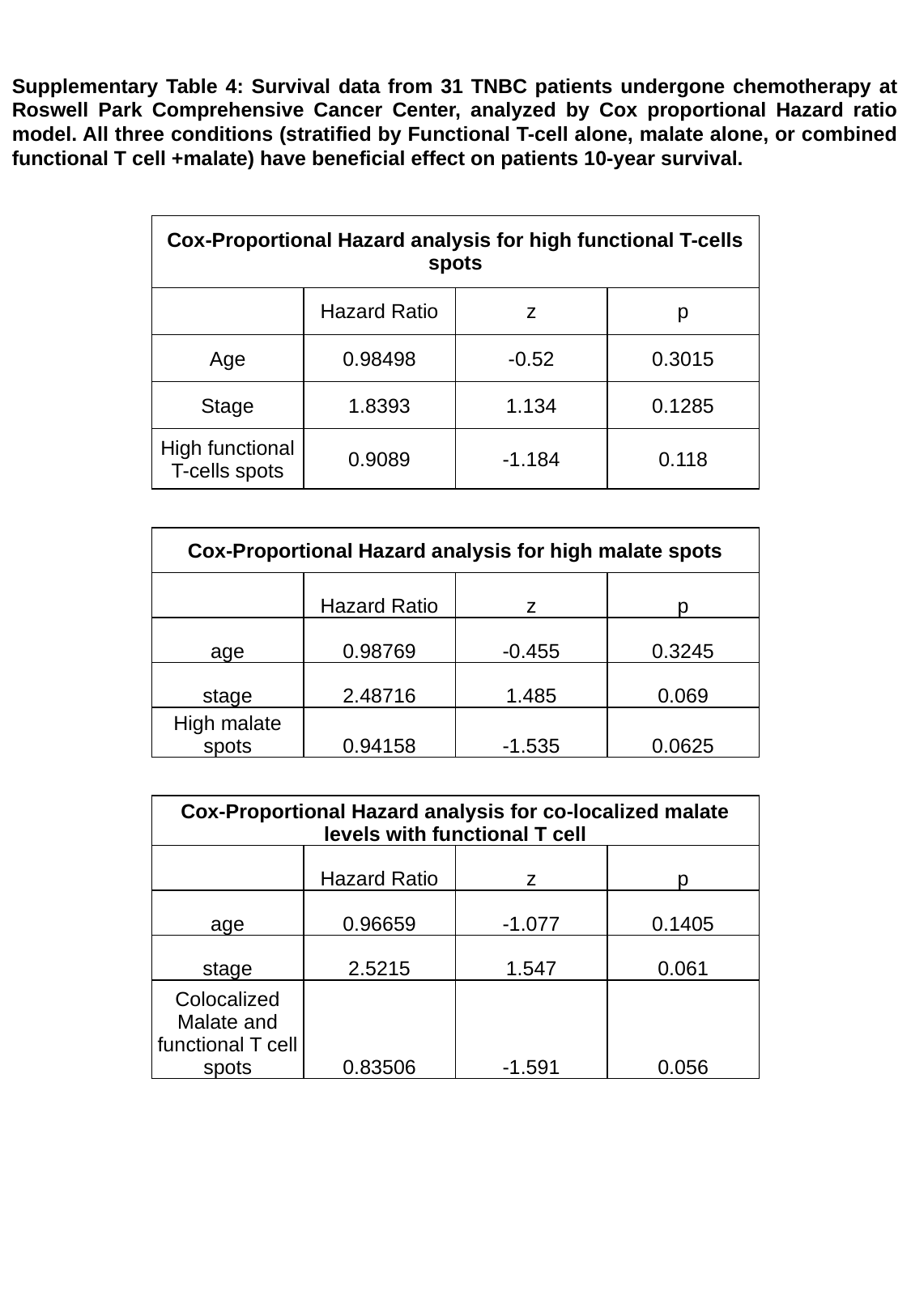

Supplementary Table 4: Survival data from 31 TNBC patients undergone chemotherapy at Roswell Park Comprehensive Cancer Center, analyzed by Cox proportional Hazard ratio model. All three conditions (stratified by Functional T-cell alone, malate alone, or combined functional T cell +malate) have beneficial effect on patients 10-year survival.
| Cox-Proportional Hazard analysis for high functional T-cells spots | | | |
| --- | --- | --- | --- |
| | Hazard Ratio | z | p |
| Age | 0.98498 | -0.52 | 0.3015 |
| Stage | 1.8393 | 1.134 | 0.1285 |
| High functional T-cells spots | 0.9089 | -1.184 | 0.118 |
| Cox-Proportional Hazard analysis for high malate spots | | | |
| --- | --- | --- | --- |
| | Hazard Ratio | z | p |
| age | 0.98769 | -0.455 | 0.3245 |
| stage | 2.48716 | 1.485 | 0.069 |
| High malate spots | 0.94158 | -1.535 | 0.0625 |
| Cox-Proportional Hazard analysis for co-localized malate levels with functional T cell | | | |
| --- | --- | --- | --- |
| | Hazard Ratio | z | p |
| age | 0.96659 | -1.077 | 0.1405 |
| stage | 2.5215 | 1.547 | 0.061 |
| Colocalized Malate and functional T cell spots | 0.83506 | -1.591 | 0.056 |

### Slide 18
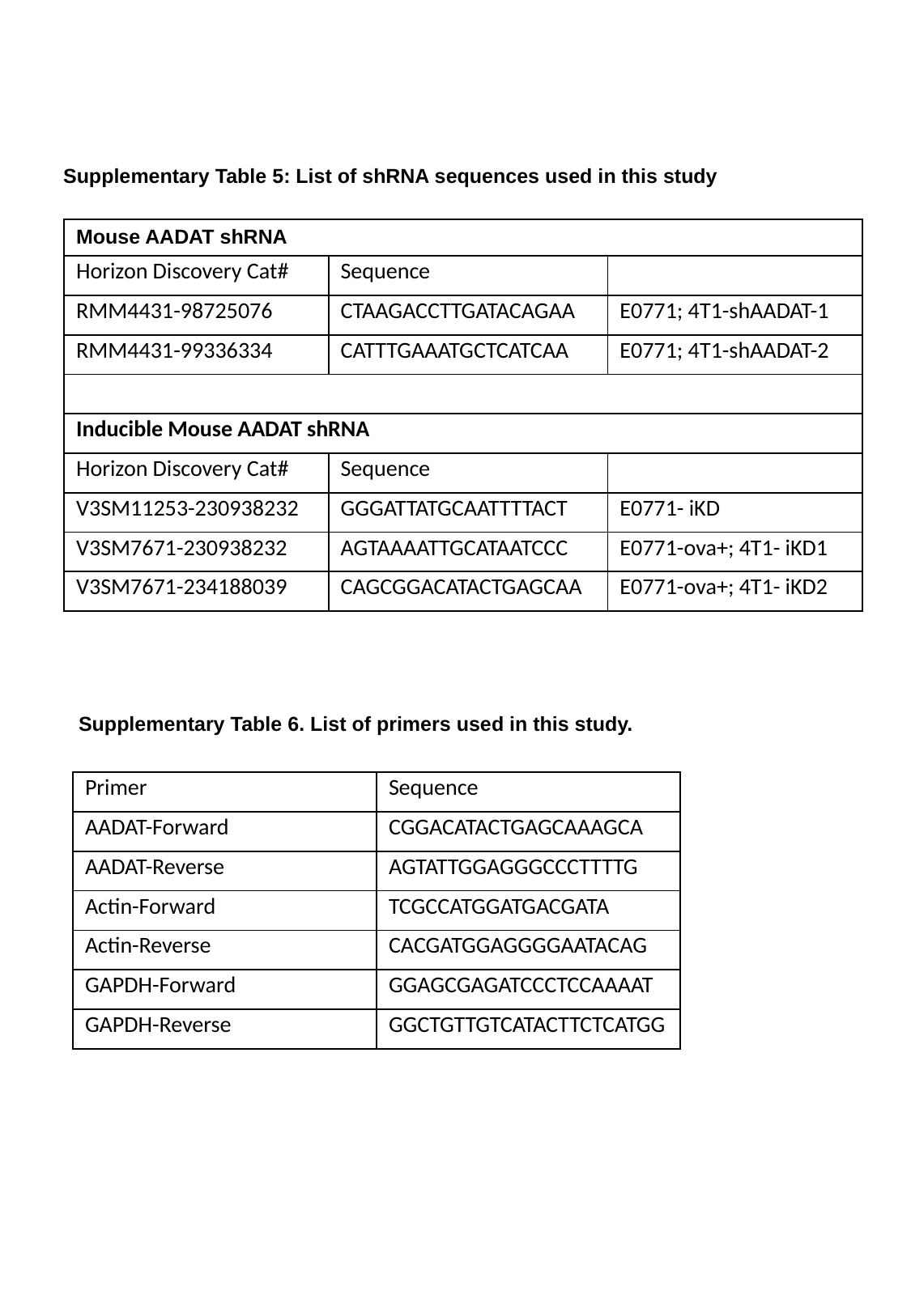

Supplementary Table 5: List of shRNA sequences used in this study
| Mouse AADAT shRNA | | |
| --- | --- | --- |
| Horizon Discovery Cat# | Sequence | |
| RMM4431-98725076 | CTAAGACCTTGATACAGAA | E0771; 4T1-shAADAT-1 |
| RMM4431-99336334 | CATTTGAAATGCTCATCAA | E0771; 4T1-shAADAT-2 |
| Inducible Mouse AADAT shRNA | | |
| Horizon Discovery Cat# | Sequence | |
| V3SM11253-230938232 | GGGATTATGCAATTTTACT | E0771- iKD |
| V3SM7671-230938232 | AGTAAAATTGCATAATCCC | E0771-ova+; 4T1- iKD1 |
| V3SM7671-234188039 | CAGCGGACATACTGAGCAA | E0771-ova+; 4T1- iKD2 |
Supplementary Table 6. List of primers used in this study.
| Primer | Sequence |
| --- | --- |
| AADAT-Forward | CGGACATACTGAGCAAAGCA |
| AADAT-Reverse | AGTATTGGAGGGCCCTTTTG |
| Actin-Forward | TCGCCATGGATGACGATA |
| Actin-Reverse | CACGATGGAGGGGAATACAG |
| GAPDH-Forward | GGAGCGAGATCCCTCCAAAAT |
| GAPDH-Reverse | GGCTGTTGTCATACTTCTCATGG |

### Slide 19
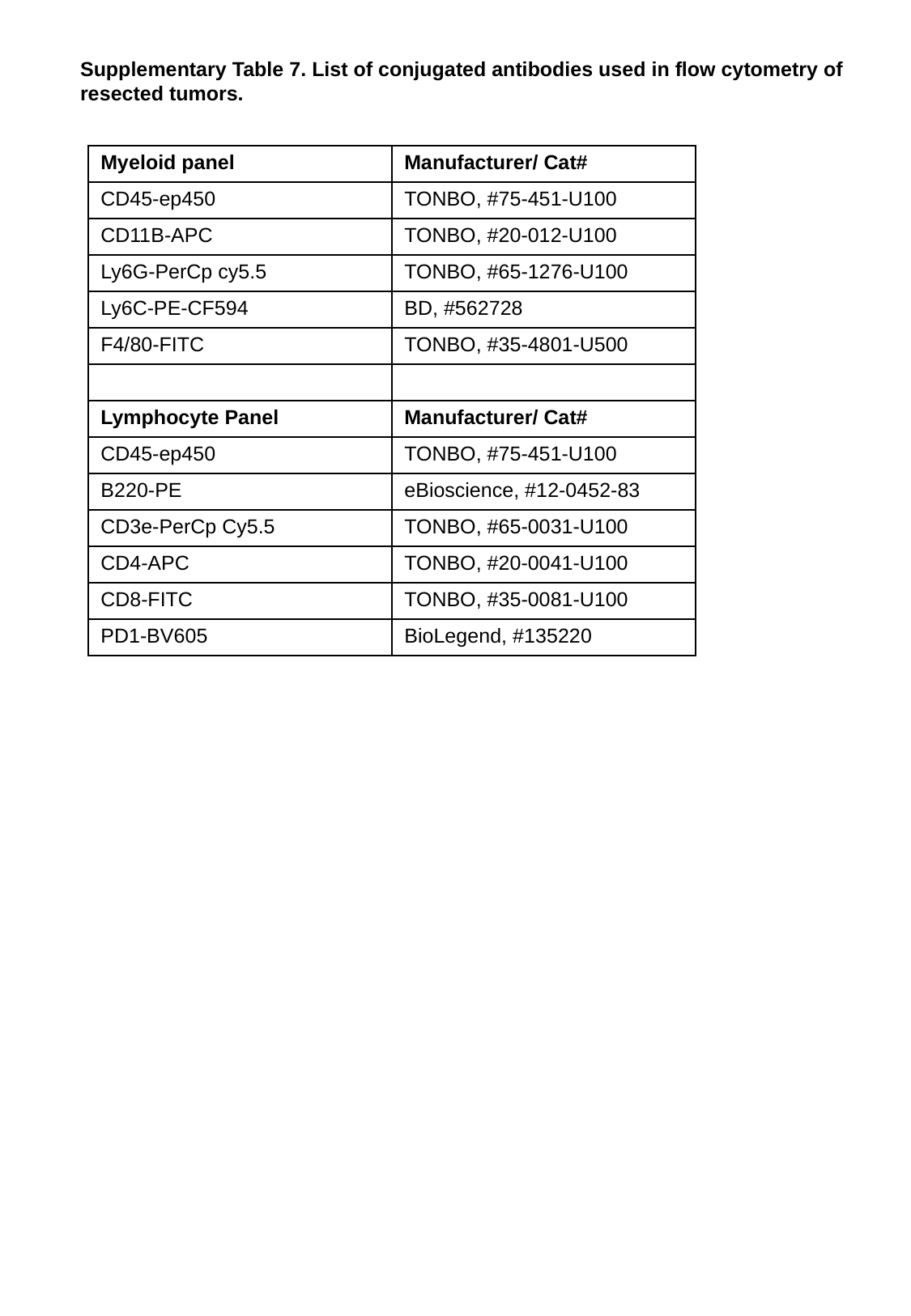

Supplementary Table 7. List of conjugated antibodies used in flow cytometry of resected tumors.
| Myeloid panel | Manufacturer/ Cat# |
| --- | --- |
| CD45-ep450 | TONBO, #75-451-U100 |
| CD11B-APC | TONBO, #20-012-U100 |
| Ly6G-PerCp cy5.5 | TONBO, #65-1276-U100 |
| Ly6C-PE-CF594 | BD, #562728 |
| F4/80-FITC | TONBO, #35-4801-U500 |
| Lymphocyte Panel | Manufacturer/ Cat# |
| CD45-ep450 | TONBO, #75-451-U100 |
| B220-PE | eBioscience, #12-0452-83 |
| CD3e-PerCp Cy5.5 | TONBO, #65-0031-U100 |
| CD4-APC | TONBO, #20-0041-U100 |
| CD8-FITC | TONBO, #35-0081-U100 |
| PD1-BV605 | BioLegend, #135220 |
